# Genetic dietary adaptation in Neandertal, Denisovan and Sapiens revealed by gene copy number variation

**DOI:** 10.1101/2021.10.30.466563

**Authors:** Riccardo Vicedomini, Niccolò Righetti, Lélia Polit, Silvana Condemi, Laura Longo, Alessandra Carbone

## Abstract

Dietary adaptation involves evolving an efficient system to digest food available in an ecosystem. The diet of archaic humans is traditionally reconstructed by isotopic analyses of human remains combined with the faunal assemblages found on the sites, and, recently, from metagenomic analyses of dental calculus. Here, we propose a new computational approach to find the genetic basis for human dietary adaptation. We searched 15 genomes from Neandertal, Denisovan and Early Sapiens for food digestion genes that tend to have more or fewer copies than the modern human reference genome. We identify 50 genes, including 10 gene clusters, with discernible copy number variation (CNV) trends at the population level, from an analysis of the full set of 20,000 human genes. The genomic variation of 19 of these genes shows how metabolic pathways for carbohydrates, lipids, liver lipids and brown fat in archaic humans adapted to metabolize food from animal or plant sources. The remaining 31 genes are all highly expressed in tissues of the digestive apparatus and are involved in immune response, environmental response and obesity. Analysis of the CNV profiles, compared to 64 modern human individuals belonging to distinct ethnic groups in Eurasia, Africa, Oceania, suggests that *Homo sapiens* may have had an evolutionary advantage compared to Neandertal and Denisovan in adapting to cold and temperate ecosystems.

**Significance Statement:** Understanding dietary strategies and foraging behaviors of past populations are among the goals of paleoanthropological and prehistoric studies. Based on a new computational approach, we seek for the genetic basis for human dietary adaptation. Gene copy number variation (CNV) is a major type of structural genome variation that we used as indicator of efficient metabolic processes for food digestion. By analysing billions of short sequences from 15 archaic human genomes and 64 modern human ones, across the full set of 20,000 human genes, we identify 50 genes whose population-wide discernable CNV trends point to lipid metabolism as being crucial for Neandertal, the efficiency in carbohydrate metabolism for Sapiens’ diet, and the importance of lipid metabolism and brown fat metabolism for both Neandertal and early Sapiens compared to modern humans.

**Preprint availability:** an early version, where the CNV analysis of ancient genomes was conducted on the human reference genome hg19, was deposited on BioRxiv on November 2, 2021 (DOI: <u>10.1101/2021.10.30.466563</u>). The current version, which utilises CNV estimates based on the human reference genome hg38 and includes comparisons with modern human populations, was deposited in BioRxiv in November 2024.

## Introduction

After the great dispersion of *Homo sapiens* (*Hs*) from Africa into Eurasia about 60,000 years ago (1), the local Eurasian archaic humans (EAHs), notably the Neandertals (*Hn*) and the Denisovans (*Den*) who populated this geographical region, disappeared. Reasons for the success of *Hs* are debated among specialists, with food intake being one factor to be considered (2,3).

According to the pioneering study of the *AMY1* gene coding for salivary amylase (4,5), high starch diets became staple during the Neolithic, when people became major consumers of cereals and other starch-rich plants, tubers and rhizomes. The inclusion of plants in the diet of *Hs* was demonstrated to provide not only a substantial caloric intake but also essential fatty acids (FAs), proteins, and other vital micronutrients (6–10). By contrast, the diet and ecological niche of EAHs were indicators of low versus high trophic level carnivores (11–14). More recent studies have highlighed that the diet of EAHs is more complex than what was thought for many years and that dietary carbohydrates consumption may have not being totally absent in their diet (15–18).

In this study, we consider differences in diet between EAHs and *Hs* by studying the copy number variation (CNV) of genes involved in digestion and metabolism. The aim of our novel computational approach is to use publicly available complete genomes of archaic humans to identify genomic changes underlying the diverse and evolving ability of humans to efficiently digest and metabolize energetic nutrients. Our CNV data show that humans living in temperate and cold latitudes had access to differentiated food sources. While both *Hn* and *Den* maintained highly carnivory, *Hs* has increased intake of dietary carbohydrates since its oldest occurrence in Eurasia, thus accessing this caloric food long before crop domestication emerged (9,19). This novel information can be interpreted in light of the archaeological and anthropological records of human populations living across Eurasia during the Late Pleistocene.

Genomic CNV has been studied for more than 30 years (20). Initially, it was thought that CNV was rare with a limited impact on the total extent of human genetic variation (21). With recent genomic technologies, thousands of heritable copy number variants (CNVs) within modern populations have been documented, generating considerable interest over the functional significance of gene duplication (21–23). CNV has been demonstrated to influence levels of gene transcription (19,24–26) conferring an adaptive advantage, and some CNVs have been associated with differential susceptibility to complex diseases (27,28). The landscape of past and present CNV in genomes could be of functional significance related to specific phenotypes within and across populations. Only a few genomes of human archaic populations will ever be available, but our CNV analysis shows that genes that maintain their copy number across individuals of a same population while changing copy number across populations can be indicators of adaptation to different environments.

## Results

More than 10,000 ancient nuclear genomes of individuals younger than 20,000 years, and only slightly more than 50 genomes from individuals older than 20,000 years, are available to date in the AADR v54.1 reference repository (29), with additional genomes also reported in the broader literature. Of those, we could only estimate gene CNV for 15 of them due to insufficient sequencing coverage, the elimination of repeated regions, and contamination of most sequenced genomes (see **Materials and Methods**). Still, this is the largest pool of ancient nuclear genomes analyzed in this way (**Figure 1A** and **Table S1**). In this pool are key genomes from individuals at sites spanning temperate and cold latitudes of Eurasia (**Table S2**) including EAHs from the Altai Mountain - Den D3 (30), Altai Neandertal D5 (31) and Chagyrskaya 8 (32) – and the earliest *Hs* from Ust’Ishim (North Siberia) (33). To this Central Asian core sample, we added *Hn* genomes following their east-west distribution, namely Mezmaiskaya 1 (34), Mezmaiskaya 2 (35), Vindija 33.19 (also referred to as Vindija 87) (34), Goyet Q56.1, Spy 94a, and Les Cottés Z4-1514 (35). In addition to Ust’Ishim, modern human genomes belonging to Peştera Muierii 1 (36), Mal’ta (37), to the hunter-gatherers from Motala 12 and Loschbour, and to the earliest northern farmer from Stuttgart (38,39) were analyzed (see **Supplementary Text 1**).

**Figure 1.**
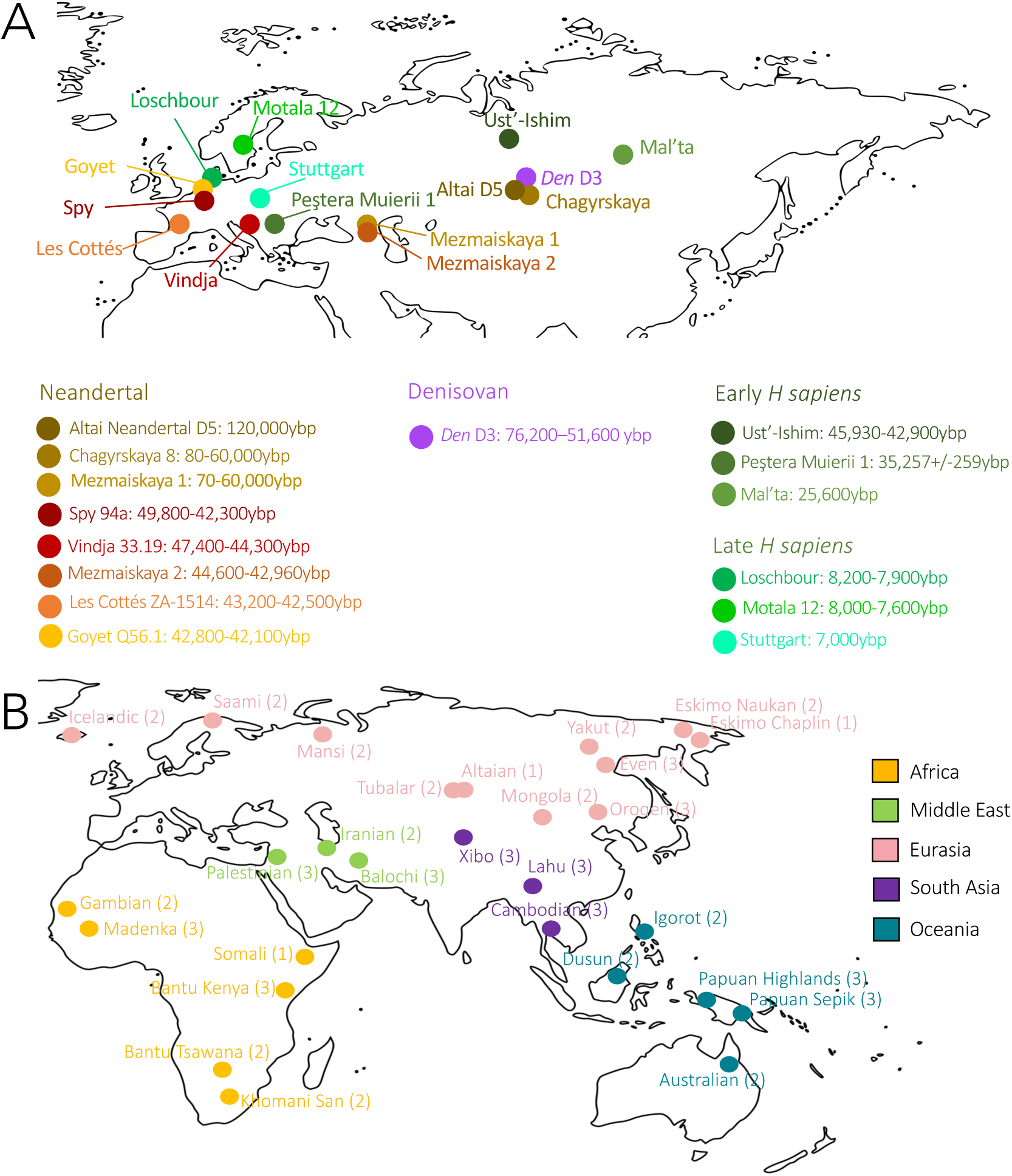
Geographical location of ancient and modern genomes used for gene CNV analysis. **A.** Ancient genomes are from individuals of early and late *Hs* (greens), *Hn* (browns, reds) and *Den* (purple) populations. Darker shades correspond to older time periods. Dates are calibrated. B. Modern genomes are from individuals from 28 ethnic groups of various geographical origins: Africa (yellow), the Middle East (green), Eurasia (pink), South Asia (purple), and Oceania (cyan). The number of individuals from each group is indicated in parentheses.

Building on this extensive analysis of ancient genomes from key Eurasian sites, we extended our comparison to include modern human populations from a wide geographical range. We analyzed 64 genomes from modern individuals across Africa, Eurasia, the Middle East, South Asia, and Oceania, representing 28 diverse ethnic groups with varying dietary practices (**Figure 1B** and **Table S3**) (40,41). This provides a comprehensive framework for understanding genetic variation across regions. This broader comparison is crucial for properly evaluating gene adaptation, as understanding how CNVs in ancient populations compare to those in present-day human groups enables us to more accurately assess evolutionary changes.

### Definition of population-dependent CNVs

To be certain of capturing the genetic changes in a population rather than specifying the genetic background of any individual, we considered genes with a differential copy number in an ancient population. First, gene copy numbers were determined individually for the genomes of *Hs, Hn, Den*, and the modern reference human genome (**Figure 2AB**; see Methods). Then, the *differential copy number of a gene* was defined for each individual as being positive if the genome had more haploid copies than the modern reference genome (GRCh38/hg38 (42)) and negative if it had fewer copies (**Figure 2C**, center). Next, we identified those genes with a positive or negative CNV in more than the half of the individuals of a population (**Figure 2C**, right; see Methods). Practically, this translates as 5 or more of the 8 *Hn* individuals and 4 or more of the 6 *Hs* individuals. Genes not showing a population-dependent CNV were not selected even though they might have been duplicated in one or more genomes within a population. As there is only one *Den* genome, the list of genes with a differential CNV from *Hn* and *Hs* has been supplemented by the CNV estimation from *Den*.

**Figure 2:**
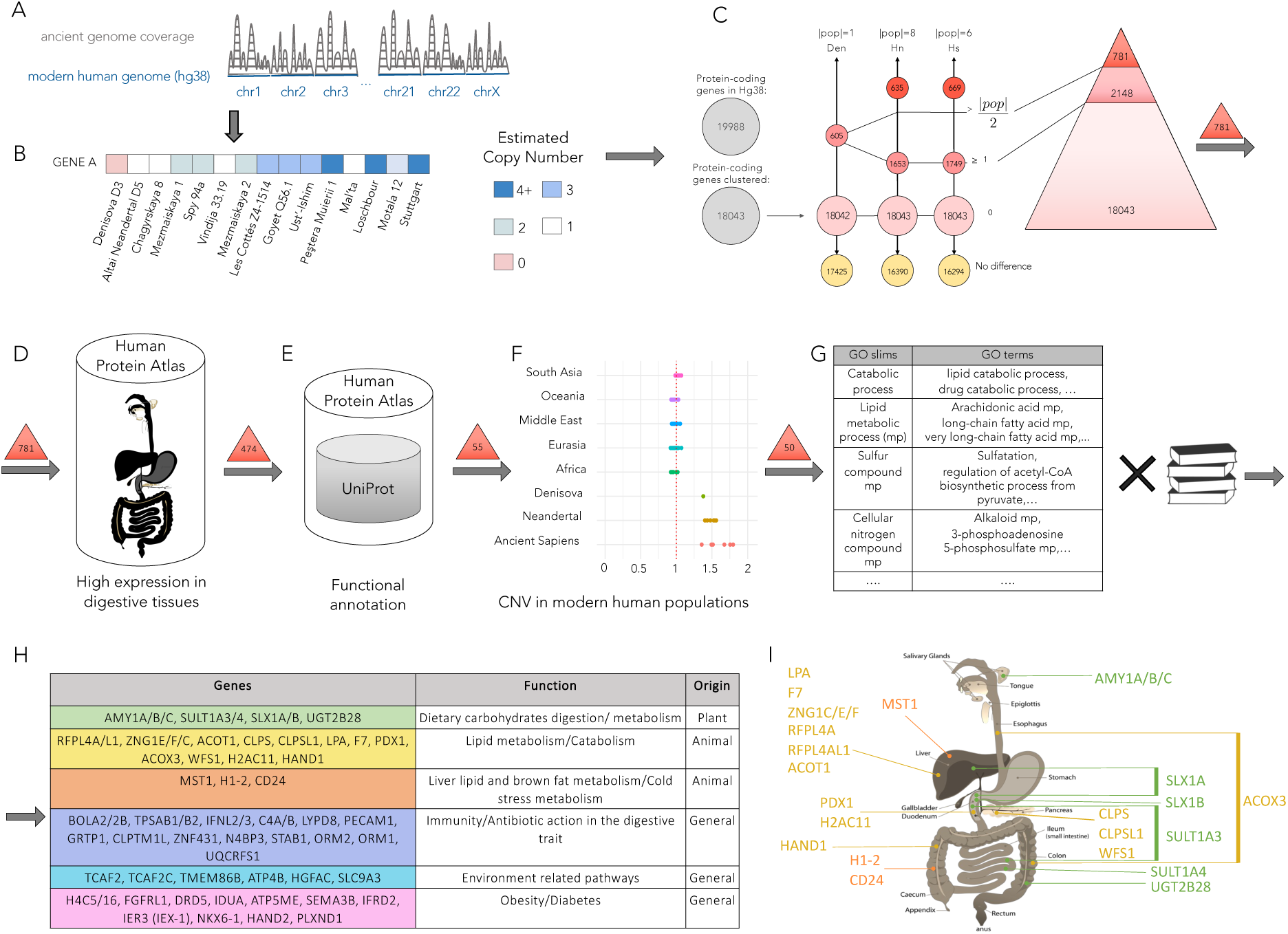
A schematic overview of the unsupervised approach identifying 19 strictly diet-related genes involved in three overarching metabolic functions and 31 accessory diet-related genes. Our automatic pipeline is defined by six main steps illustrated in panels A-F (see Methods and **Supplementary Text 2** for details). **A.** Coverage estimation of chromosomes and genes in ancient genomes after read mapping on the modern human reference genome hg38 comprising 19,988 genes, or 18,043 genes if gene clusters are counted instead. **B.** Gene haploid copy number estimation realized for the 15 ancient genomes provides CNV profiles for 18,043 genes in *Hn* and *Hs* and 18,042 in *Den* (see Methods). **C.** For *Den*, *Hn* and *Hs*, the different pink shaded circles detail the number of genes displaying a differential copy number for >n individuals in the population, where n=0…|pop| and |pop| indicates the total number of individuals in a population; for instance, over the total of 18,043 genes, 1,749 genes in *Hs* display a CNV profile different than hg38 on at least one individual, and 669 genes display a CNV profile greater or smaller than hg38 on at least 4 *Hs* individuals for a total of 6. Based on this analysis, the 18,043 clustered human protein genes (grey circle) are split in three subgroups (triangle): 18,043 genes are analysed across the 15 ancient genomes and hg38 (light pink, n=0), 16,294 (16,390) of which display no differential CNV in *Hs* (*Hn*) genomes with hg38 (yellow); 1,749 (1,653) genes have at least one individual across the *Hs* (*Hn*) population with CNV different from hg38 (pink, 1≤n≤|pop|/2); 781 genes have at least one population which displays more than the half of its individuals with a greater or smaller copy number (red, n>|pop|/2). **D**. From the 781 genes selected in **C**, we screened those with primary expression is digestive system organs using data from the Human Protein Atlas (HPA) database (**Table S4**). **E.** These genes were further filtered through a manual review of UniProt functional annotation reported in HPA to confirm their role in digestion, resulting in a set of 55 genes. **F.** These 55 genes were then compared to CNV estimations from five populations across Africa, Eurasia, South Asia, Middle East and Oceania to assess adaptive trends, narrowing the set to 50 genes. **G**. A GO term and literature-based analysis of these 50 genes identified their metabolic functions and specific roles in digesting plant or animal foods, or in “general” digestion-related processes. The results are shown in **H** (see also **Table S5**). **I.** Of these, 19 strictly diet-related genes are mapped to the digestive organs where they are primarily expressed. [The original drawing of the digestive apparatus was released into the public domain by the author LadyofHats.]

We used an unsupervised approach to analyze the full set of 20,000 human genes for the 15 genomes (**Figure 1A**) to identify a subgroup of 781 genes based on the differential haploid copy number of the gene for a population (**Figure 2ABC**). These 781 genes are expected to highlight population-dependent characteristics of the genomes and constitute the “tip of the iceberg” of the entire collection of human genes. Here, being interested in investigating the human diet, we further filtered them on the basis of two other criteria: their principal expression in organs of the digestive apparatus (**Figure 2DE** and **Table S4**) and a comparison with CNV estimations from five modern human populations to assess their adaptive trends (see **Materials and Methods**; **Figure 2F**). This process led to the identification of 50 genes, which were then analysed using Gene Ontology classification (43) and digestion-related literature (**Figure 2G** and **Table S5**). Of these, 19 genes were involved in three strictly diet-related overarching metabolic functions: carbohydrate, lipid, and liver lipid/brown fat metabolisms. The remaining 31 were associated with auxiliary roles in diet, including functions related to the immune system, cold response, and obesity (**Figure 2HI**). The computational approach is large-scale and allows genome-wide gene analysis while remaining accurate in estimation, as indicated in the discussion and in the detailed analysis of the *AMY1A/B/C* gene cluster, involved in carbohydrate metabolism, compared to *AMY2A* and *AMY2B* genes reported in **Figure S1**. (See **Materials and Methods** and **Supplementary Text 2** for all details.)

The panel of 50 genes is primarily expressed in digestive organs, with most showing expression in a specific organ (**Figure 2G** and **Table S4**), according to data from the Human Protein Atlas (HPA) (44). However, certain genes, such as *ACOT1*, which is involved in fatty acid metabolism and cellular energy balance, and *MST1*, which plays a role in liver lipid and brown fat metabolism, exhibit a much wider expression across multiple tissues. The CNV observed may therefore be an adaption to processes other than digestive activity. We excluded from our analysis any food metabolism genes (45) which are not principally expressed in the digestive apparatus (44), even if they presented a differential CNV in a population. We also excluded those genes that have not yet been associated with a known biological function (46), even though their high expression in digestive organs and differential CNV suggest they might play a role in, or be influenced by, digestion. The stringency of the expression criterion and the functional criterion used here may mean that the current panel of genes is an underestimate of those potentially responsible for CNV-driven food adaption.

For the 50 genes selected with a differential CNV in a population, we checked whether there was a *CNV trend* in the population, that is in 5 or more of the 8 *Hn* individuals and 4 or more of the 6 *Hs* individuals (see Methods). The *AMY1A/B/C* gene cluster (**Table 1A**; **Table S6**), for example, was selected by a negative differential CNV with 7 out of 8 *Hn* individuals having fewer gene copies than modern humans (**Table 1A**). The *Hs* population does not show a positive CNV trend, even though 4 of the 6 genomes have at least the same number of *AMY1A/B/C* gene copies than the modern genome (**Table 1A**). Determining the differential CNV trend of a gene in a population revealed several aspects. The same gene may show different trends between ancient populations: *AMY1A/B/C* and *SULT13/4*, for instance, have fewer gene copies in *Hn* and *Den*, but more or on par in *Hs*. Trends may be positive and negative in a single population: in *Hn*, the differential CNV of *ACOT1* is negative, while that of *ACOX3* and *HAND1* is positive. The same trend was shared by populations: for example, for *TPSAB1/B2* and *IFNL2/3* show a negative trend in both *Hn* and *Hs*, whereas *CLPS* and *CLPSL1* show a positive trend in both *Hn* and *Hs*. As expected, CNV estimations in *Den* very often agree with the *Hn* trend (47). The differential CNV captured chronological trends within populations, as shown for the *HAND1* gene in *Hs*, where duplication appears in Mesolithic (late hunter-gatherer) individuals. Another example are the 5 and 6 copies of *AMY1A/B/C* observed in Loschbour and Stuttgart (Neolithic, early farmers) genomes (**Table 1B**), recently highlighted in (48).

**Table 1.**
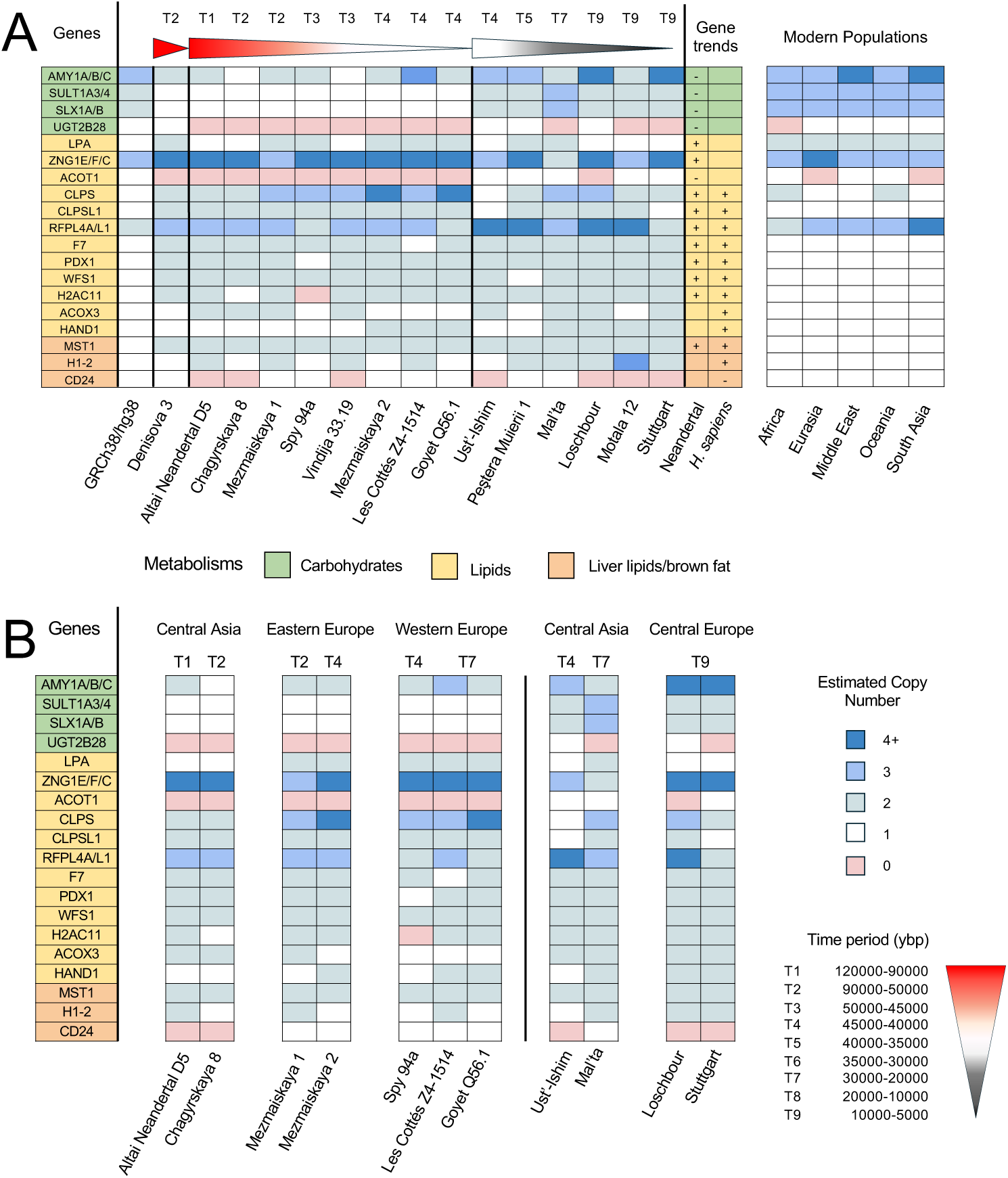
Estimated haploid copy number for genes involved in digestion in ancient genomes and differential CNV analysis in ancient and modern populations. **A.** Copy number of 19 genes (rows) in 15 ancient genomes and the modern human reference (columns). Individuals are grouped according to their population, from left to right, the GRCh38/hg38 reference, *Den, Hn* and *Hs*. Within a population, the order reflects time period and geographical origin. The estimated copy numbers computed for each gene/genome (see **Supplementary Text 2**) are rounded to the closest integer (see **Table S6** for specific values). The order and the color shading for gene names correspond to those used in **Figure 2A**. On the right, the differential CNV trends of each human population are scored for each gene as positive (+), negative (-), not determined (NA) or no differential trend (no symbol). The far right table displays the CNV medians computed for each gene in modern populations across 5 different geographical areas, each representing various ethnic groups (**Table S3** and **Table S7**). Compare to **Figure 3**. B. CNV profiles of ancient genomes in A grouped by geographical area.

Among the 50 genes, we found the well-studied *AMY1A/B/C* gene cluster (4,5,9,10). The negative CNV trend seen in *Hn* and *Den* populations (**Table 1** and **Table S6**) confirms the key role of the cluster in *Hs* adaptation to a different mode of food digestion and metabolism. Some of the remaining 18 genes, such as *ACOT1* (49) and *CLPS* (23), have previously been identified as key players in human dietary adaptation. In this study, we highlight their primary roles and propose an expanded panel of genes for further investigation. The 19 genes cover a broad spectrum of metabolic functions. The most striking feature is that for each overarching metabolic process represented, either all the genes involved in that process present a positive differential CNV for a population or a negative one (see **Table 1A** and **Table S6**, rightmost columns). This is a strong indication that an adaptive force underlies the duplication of these genes in archaic populations. A detailed description of the 50 genes is reported in **Supplementary Text 3.**

It should be highlighted that the dataset of 50 genes has been identified after a comparison with CNV data from modern individuals (**Figures 3** and **Figure S2**). Their CNV was estimated in the same way as for ancient individuals (**Figure 1AB**), and for each gene and each of the five modern human populations, the median CNV distribution (**Tables 1** and **2**) was considered to represent the CNV of the modern population. This comparison revealed that while CNVs in the human reference genome generally align with CNV medians of the modern populations, there are certain populations where they do not. This suggests that the reference human genome alone is insufficient for evaluating CNV variability and that comparison with modern human populations should be integrated into CNV analyses. We excluded five genes showing these differences (**Figure S3ABC**) and identical CNV median for both ancient and modern populations, suggesting no evidence for environmental adaptation for these genes. For instance, the *PGA3/4/5* gene cluster (Pepsinogen A, group 1), involved in the breakdown of animal proteins and highly expressed in the stomach, shows a differential CNV between hg38 and nearly all *Hn* and *Hs* individuals, with more than 4 copies observed as a median in Eurasia, Oceania and South Asia populations, aligning with the high duplication in both *Hn* and *Hs* (**Figure S3CD**). We conclude that this gene is essential for the human diet across both ancient and modern populations; however, its duplication seems unrelated to environmental adaptation in ancient populations.

**Figure 3.**
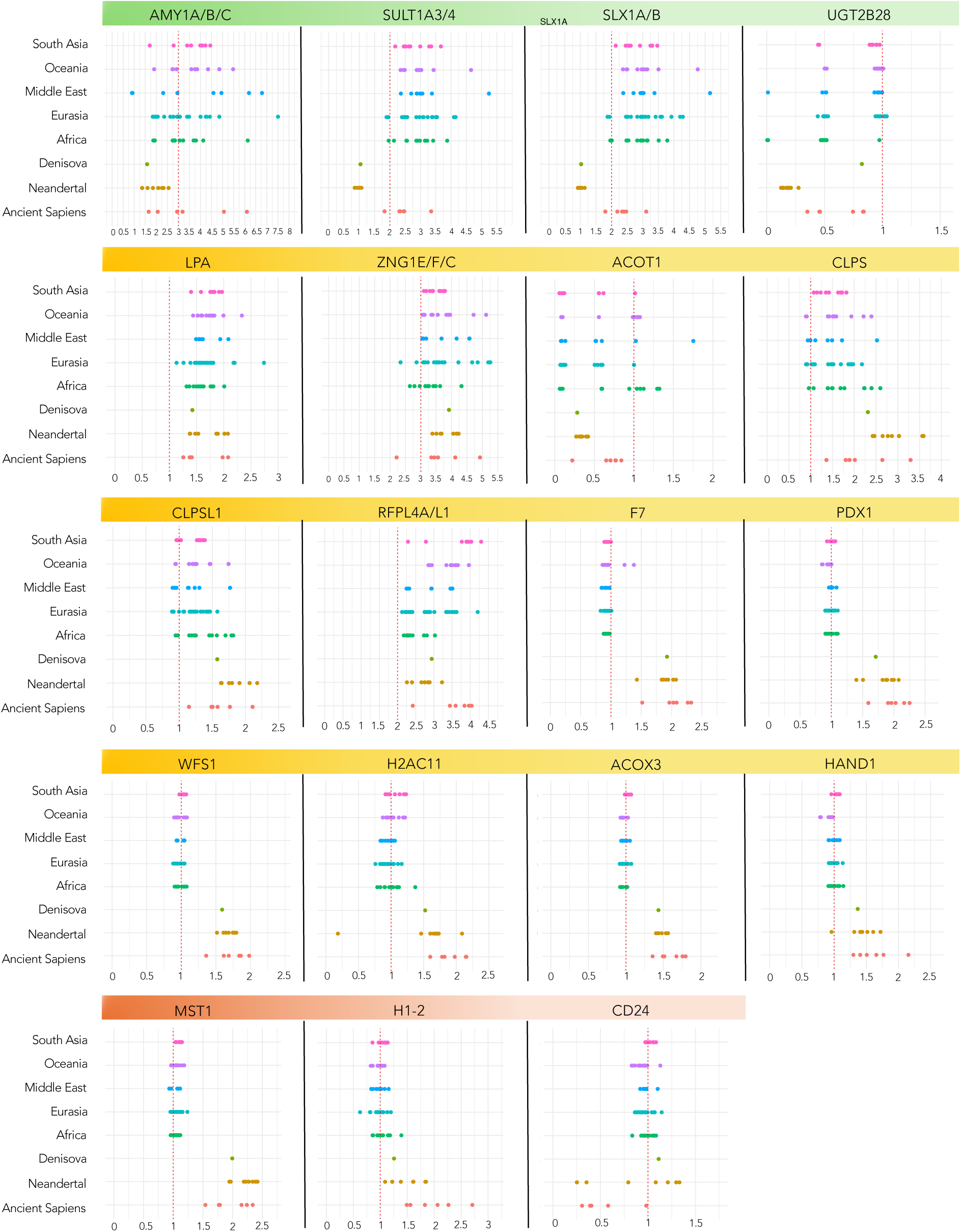
Estimated haploid copy number for the 19 strictly diet-related genes in ancient and modern genomes. For each gene in **Table 1** and for each individual in modern and ancient populations, the estimated CNV is provided. See **Figure S2** for the full set of 50 diet-related genes. The red vertical bar indicates the CNV in the reference genome hg38. Green, yellow and red gene labels correspond to the carbohydrate, lipid, liver lipid/brow fat metabolisms, respectively.

In **Figure 3A** and **Table 1A** (see **Figure 3C** and **Table 2A** for the 31 accessory diet-related genes), the comparison between CNV distributions between ancient and modern populations reveals three main patterns: 1. Modern populations may align with the CNV estimation from the hg38 reference genome. This is observed in lipid metabolism genes such as *F7, PDX1, WFS1, HCAC11, ACOX3*, and *HAND1*, as well as liver lipid/brown fat metabolism genes *MST1, H1-2, CD24*, and the carbohydrate metabolism gene *UGT2B28*. In these cases, the CNV median for ancient populations, including *Hn*, *Den* and *Hs,* differs from that of modern populations. 2. Modern populations can exhibit a wide range of CNV values, with their median aligning with hg38, but diverging from the CNV median of at least one ancient population. This pattern is seen in the carbohydrate metabolism gene *AMY1A/B/C* and lipid metabolism genes *ZNG1E/F/C, ACOT1, CLPS*, and *CLPSL1*. 3. In some cases, modern populations show a wide range of CNV values, but their median does not align with the hg38 CNV estimation, while the CNV median in at least one ancient population differs significantly. This is the case for the carbohydrate metabolism genes *SULT1A3/4* and *SLX1A/B* genes, which display lower CNV values in *Hn* compared to modern humans. Similarly, lipid metabolism genes *LPA* and *RFPL4A/L1* show lower and higher CNV medians in *Hs*, respectively, compared to modern populations.

**Table 2.**
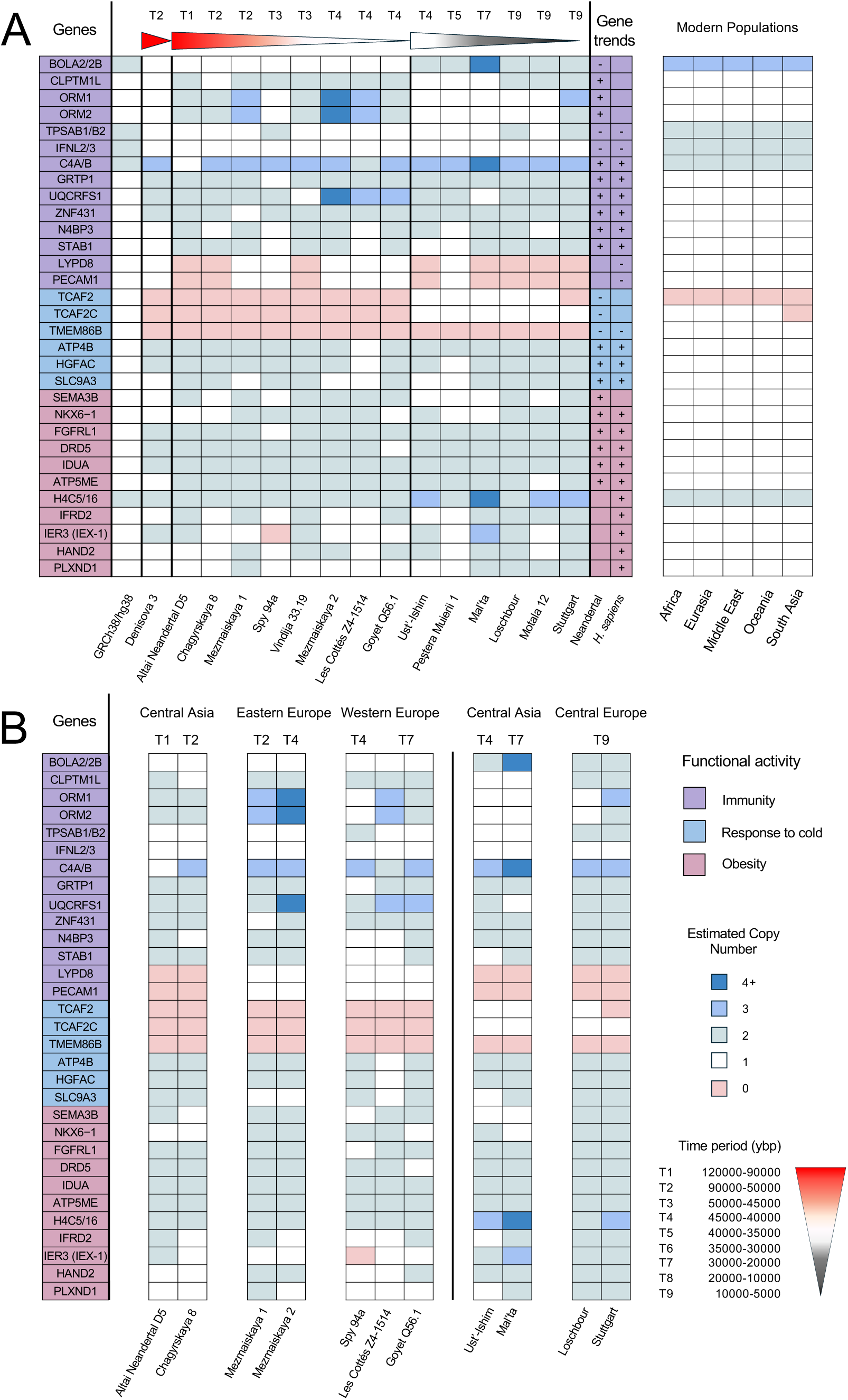
Estimated haploid copy number for digestion-related genes in ancient genomes and differential CNV analysis in ancient and modern populations. Description of the 31 accessory genes related to digestion. Legend as in **Table 1**.

### A multidimensional genetic space defined by CNV

We further test our main hypothesis that dietary changes directly cause a significant reshaping of individual genomes within populations, and that this phenomenon can be decoded today through large-scale analysis of gene copy number. For this we conducted a PCA analysis, which reveals how ancient and modern genomes can be grouped together in gene space. Based on the CNV of the 19 genes involved in digestion listed in **Table 1A**, we visualized and compared the 15 genomes by plotting them in a 19-dimensional space, where each dimension corresponds to a gene. Formally, we represent a genome of an individual with its CNV profile as described by the relevant column in **Table 1A**. In this multidimensional space, two individuals are close when their CNV profiles are similar. To aid visualization, we reduced the space by principal component analysis (PCA) (50). The PCA projection (**Figure 4A)** guarantees the largest variation in the data points through an optimized linear combination of the 19 original dimensions (see **Supplementary Text 2**). We built two 19-dimensional spaces, with (**Figure S4A**) or without (**Figure 4A**) the modern human hg38 profile included.

**Figure 4.**
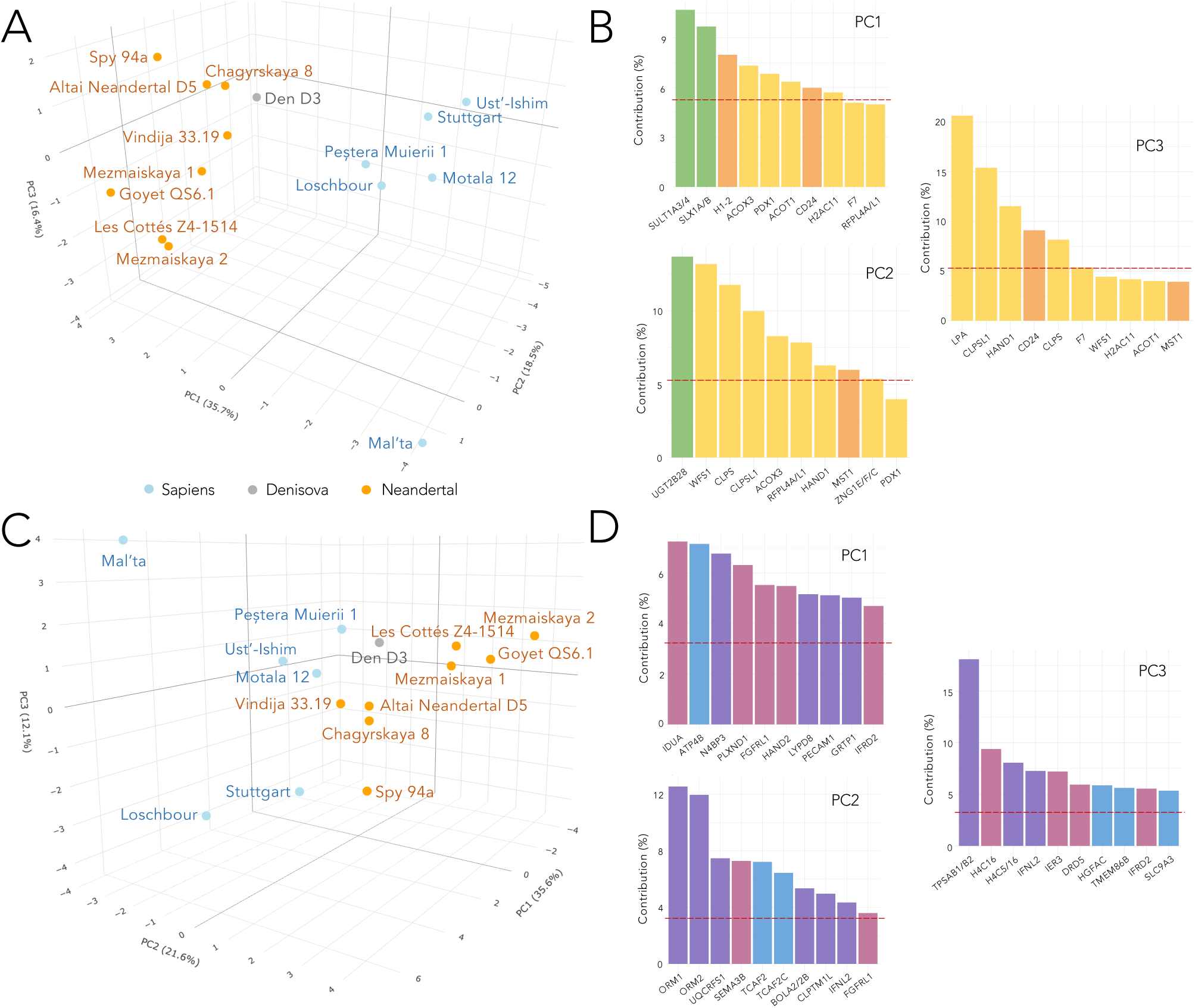
Comparative CNV analysis of *Hn*, *Den* and *Hs* living in the European belt. **A.** Fifteen *Hn*, *Den* and *Hs* individuals are represented in a three-dimensional space obtained by PCA from the 19-dimensional space defined from differential CNV values of the 19 genes involved in metabolic processes (see **Supplementary Text 2**). View of the 3D space where each individual (points) is colored with respect to human populations: *Hn* (orange), *Den* (grey), and *Hs* (light blue). PC1 explains the 35.7% of the variance, PC2 the 18.5% and PC3 the 16.4%. **B.** Contributions of factors for each principal component of the 3D space in **A**. Colored bars indicate the contributions from CNV of genes acting in the three overarching metabolic processes (colors as in Figure 2F). **C.** Same analysis as in **A** but based on the differential CNV values of the 31 accessory genes. PC1 explains the 35.6% of the variance, PC2 the 21.6%, and PC3 12.1%. **D.** Contributions of factors for the definition of each principal component of the 3D space, obtained by PCA for the 31 genes analysis in **C**. See legend in **B**.

In **Figure 4A**, the CNV data clearly separate *Hn* and *Hs* (see the first component PC1 in **Figure 4A**). Den D3 localises close to *Hn*. Classification based on geographical origin (**Figure S5A**) or time period (**Figure S6B**) shows coherent subgroupings. Comparing CNV profiles of the individuals living in the same geographical area (**Table 1B**) illustrates how they are close in the high-dimensional space even when they belong to very different time periods. For instance, Mezmaiskaya 1 and Mezmaiskaya 2 both lived in Eastern Europe (Caucasus) and their similar CNV profiles (**Table 1B**) bring them close together in the CNV space (**Figure S5A**, light blue dots), even though they were not contemporary, living in T2 and T4 respectively.

Analysis of the PCA axes of **Figure 4A** reveals that the most important factors explaining the first dimension (PC1) of the CNV projection separating *Hn* from *Hs*, are genes involved in carbohydrate digestion, *SULT1A3/4* and *SLX1A/B*, and to a minor yet statistically significant extent genes involved in liver lipids metabolism, *H1-2* and *CD24*, and genes involved in lipid metabolism, *ACOX3, PDX1, ACOT1*, and *H2AC11* (**Figure 4B**). The second principal component of the CNV space is mainly accounted for by the gene *UGT2B28* involved in carbohydrate digestion, and the genes involved in lipid metabolism (WFS1, *CLPS, CLPLS1, ACOX3, RFPL4A/L1, HAND1*). Finally, the liver lipid/brown fat metabolism gene *CD24* and the lipid metabolism genes *LPA, CLPSL1, HAND1*, and *CLPS*, make a significant contribution to the third dimension.

In **Figure S4A**, the CNV data clearly separate *Hn, Hs* from the modern genome hg38 (PC1 in **Figure S4AC**) and also separate *Hn* from *Hs* (PC2 in **Figure S4AC**). The first principal component (PC1), separating modern from ancient genomes, is mainly explained by several lipid metabolism genes (*ACOX3, PDX1, WFS1, F7, HAND1, H2AC11, CLPS*), and liver lipid metabolism genes (*MST1* and *H1-2*). The second component (PC2), separating *Hn* and *Hs* genomes, is explained by genes involved in carbohydrate digestion (*SLX1A/B, SULT1A3/4* and UGT2B28). This ensemble of contributions, taken together, underscore the significant dietary shift of modern populations compared to ancient ones, who primarily consumed high-fat and protein-rich animal products (11–15,17,18), whereas modern diets are characterized by a more diverse range of nutrients that include a growing rate of dietary carbohydrates.

The dietary habits of ancient and modern populations are further reflected at the genomic level through a signature based on an independent set of 31 accessory genes. This suggests that the genomic differences between *Hn* and *Hs* related to their diets extend beyond immediate metabolic differences, strongly influencing other biological processes such as immune response, adaptation to cold environments, and obesity. We run our analysis over the complementary set of 31 accessory genes in **Table 2A.** These genes are highly expressed in the digestive apparatus. This analysis provided the same genome split for the first and second PCA components (**Figure 4B**) identified by the set of 19 strictly diet-related genes. Interestingly, the separation of *Hn* from *Hs* (PC2; **Figure 4BD**) is obtained with the main contribution of genes which are involved in immunity (*BOLA2/2B*, *ORM2, UQCRFS1, ORM1, PECAM1, C4B*), response to cold (*TCAF2C, TCAF2*), and obesity (*H4C16*, *IER3*). It is noteworthy that the main factors contributing to the splitting between modern humans from ancient humans (PC1 in **Figure S4BD**) are principally involved in obesity (*IDUA, PLXND1, HAND2, FGLR1*) and immune response (*N4BP3, CLPTM1L*).

To summarize, our analysis of the 19 genes primarily involved in digestive functions and displaying a differential copy number provides an unprecedented basis from which to deduce the adaptive changes between EAHs and *Hs* regarding dietary strategies. This study underscores the critical role of three distinct metabolic processes in shaping the behavior of ancestral populations and suggests that these 19 genes could serve as reliable markers for tracking changes in dietary adaptation over time. The CNV genetic trend (**Figure 4A)** points to lipid metabolism as being crucial for both *Hn* and *Hs*, while efficiency in carbohydrate metabolism influences the diet of *Hs*. Lipid and liver lipid/brown fat metabolisms must have been important for both *Hn* and *Hs* compared to modern humans and were probably already present in the common ancestral lineage.

## Discussion

In this study, we analyzed 15 complete ancient genomes to identify CNV trends in genes across three human populations present in the vast Eurasian territory during Late Pleistocene— *Hn*, *Den* and early *Hs*. By examining the largest number of publically available full genomes suitable for CNV analysis, our study significantly extends the list of genes with unusual copy number in archaic genomes. By remapping ancient genome reads to the modern human reference genome hg38, we accessed the most up-to-date CNV estimations for comparative analysis. Our genome-wide CNV analysis identified all genes with increased or decreased CNVs in ancient populations relative to the human reference genome, uncovering 781 genes of high interest through an unsupervised search. Among these, we focused on genes involved in food digestion and metabolism, revealing the complexity of nutritional adaptation among the three human populations that shared the same environmental conditions in Eurasia at the end of MIS 4 and during MIS 3. From the genomic data, three overarching metabolic pathways related to diet emerged as overrepresented in the 781-gene dataset: carbohydrate metabolism, lipid metabolism, and liver lipid and brown fat metabolism. These findings provide evidence of distinct adaptive nutritional pathways that may underlie the dietary habits of these populations. Most importantly, the study offers a novel perspective to look at longstanding questions, challenging established results with new genetic evidence. It contributes unique insights into the physiology and biology of past humans that cannot be gleaned from the fossils and archeological records. However, it should be noted that functional complexity, epigenetic regulation, and microbiotic interaction make it challenging to precisely determine the roles of these genes in metabolic activities, even in present-day human populations.

### A genome-wide methodological approach based on multiple selective criteria

Distinguishing between adaptive from neutral evolution, especially in hominids which have low effective population sizes and whose available genomes are limited, has been a major challenge (21,23,22,38,49,51,22,52). In this article, we highlight the difficulty in such a selection and propose that a combination of criteria can lead to the identification of a set of candidate genes to play a key role in the dietary adaptation of ancient populations. We search for a pool of genes with differential copy number in ancient populations by analysing, at large-scale, 20 000 human genes with four main selection criteria:

**1.** a CNV differentiation in ancient populations (not individuals). Focusing on “populations” rather than “individuals” helps identify CNVs that are consistent across a group, implying a broader evolutionary significance of the gene copies. CNV differentiation across ancient populations could highlight long-term selection pressures that shaped the genetic landscape over time. The intuition being that CNVs that show marked differentiation in ancient populations might reflect early environmental pressures (e.g., climate, diet, or pathogens) that shaped genetic adaptations crucial for survival.
**2.** a comparison of modern populations to ancient ones, to track how certain CNVs have persisted, expanded, or diminished over time. This comparison helps in distinguishing adaptive CNVs that were beneficial in the past and might still be advantageous in some modern environments, or those that were selected against and have decreased. CNVs that have been maintained or show a selective sweep in modern populations likely reflect ongoing adaptive advantages, such as responses to new environmental factors (e.g., agriculture, modern diets, urbanization, or exposure to new pathogens).
**3.** the high expression of the gene in specific tissues from digestive organs. Tissue-specific high-expression of genes, especially in critical systems like digestive organs, suggests a key functional role. If the CNVs influence gene expression in tissues tied to nutrient absorption, metabolism, or immune responses in the gut, this could indicate an adaptive response to dietary changes, such as the ability to process different food sources (e.g., starch, lactose), or microbial challenges, to fend off gut pathogens.
**4.** the functional annotation of the gene. Functional annotation helps clarify the biological role of the genes affected by CNVs. By linking CNVs to specific functions (e.g., immune response, metabolism, etc.), we can pinpoint whether these genetic changes have played a role in specific traits. For example, if a CNV is associated with genes involved in immune function, this might suggest adaptation to pathogen pressures, which is a key driver of human evolution.

The combination of these criteria strengthens the argument that CNVs can serve as markers for human adaptation. Note that the criteria for gene selection under these multiple markers were very strict and allowed to identify 50 genes related to diet which likely contributed to adaptive changes of ancient populations. The stringency of the criteria may also mean that the current panel of genes is an underestimate of those potentially responsible for CNV-driven food adaption. Our 50 genes should be considered as the core set for digestion.

### High read coverage and CNV estimation in ancient genomes

Our computational approach underscores the critical need of sequencing complete ancient genomes at high coverage, as emphasized by previous studies (48,53). Read coverage analysis of the 50 diet-related genes (**Figures S7 and S8**) demonstrates that while CNV estimation is feasible, mapping sensitivity is highly influenced by coverage depth. High-coverage genomes, including Den D3, Altai Neandertal D5, Chagyrskaya 8, Ust’Ishim, Peştera Muierii, Vindija, Loschbour, and Stuttgart (**Table S1**) exhibit significantly reduced mapping fluctuations compared to lower-coverage genomes, as illustrated by the smoother curves in **Figure S7** for the CLPS gene and **Figure S8** for the AMY1A/B/C cluster. Moreover, sequencing entire genomes, rather than focusing solely on SNPs or regions, enables to uncover significant aspects of human genome evolution, such as gene copy number variations and mutations in coding or regulatory regions—some of which may not be observable in modern populations. Comprehensive sequencing is thus more essential than ever for advancing our understanding of human evolutionary history.

Another key aspect of our approach is the estimation of the haploid CNV values, which are rounded to the nearest integer and used as indicators of duplication. These estimations are provided and can support further analysis of CNV. A key question arises regarding the method’s ability to distinguish between closely related but distinct gene copies. For instance, the highly similar *AMY1* and *AMY2* genes can be correctly differentiated using our haploid genome approach. **Figure S1A** demonstrates our method’s estimation of CNV for *AMY1A/B/C, AMY2A,* and *AMY2B* in both modern and ancient populations. The amylase locus, known for its genotyping challenges, underscores the importance of verifying CNV accuracy, particularly for distinguishing *AMY1* from *AMY2* in diploid genomes. Our results align with previous studies (54), showing that modern populations consistently have one copy each of *AMY2A* and *AMY2B*, while *AMY1A/B/C* is typically duplicated with a median of three copies. Further supporting the accuracy of our haploid estimations, the distribution of paired and unpaired copy numbers in corresponding diploid CNV analyses agrees with earlier studies on modern genomes, as illustrated in **Figure S1B**.

### Comparison to modern populations

Another critical aspect highlighted in this study is the importance of the comparison with modern humans (23,55), which helps us better assess whether gene duplication truly represents adaptation, as argued above, by taking into consideration the *PGA3/4/5* gene. Additional information, such as whether a gene is located in copy number variation hotspots across multiple primates, is also crucial for evaluating the gene’s likelihood of being duplicated (56). This was the case in (57), which explored the duplication of the *PGA3/4/5* gene. In our analysis, we excluded this gene because its duplication in modern populations—facilitated by its location in a hotspot—does not distinguish ancient genomes from modern ones (**Figure S3**). Our large-scale approach, comparing modern and ancient genomes, indirectly captures aspects at the genome organisation level.

### On the generality of the method

While diet plays a significant yet partial role in the evolutionary history of ancient populations, our computational approach is broad and applicable across the entire set of 20,000 genes in our 15 ancient genomes. It provides a foundation for future research into identifying differential CNVs for other functional domains, such as cognition, morphology, sensory perception, or immune response, although further development will be needed to fully explore this potential.

### Unsupervised identification of genes presenting common CNV trends

The 19 strongly diet-related genes identified through our unsupervised approach are involved in three key metabolic processes: carbohydrate metabolism, lipid metabolism, and liver lipid/brown fat metabolism. Each metabolic group includes genes with either positive or negative CNV trends within a population. In our view, this is a strong indication that an adaptive force acted on the duplication of these genes in archaic populations, shaping their metabolic capacities with consequences on dietary habits and adaptation to diverse environmental conditions. Our data also indicate that gene copy numbers can differ between earlier and later individuals within the same population, as seen with *AMY1A/B/C* in *Hn* genomes and *LPA* in *Hs* genomes. These chronological trends can be clarified as more high-coverage genomes become available to this analysis. The 31 accessory diet-related genes fall into three main functional categories: immune response, thermoregulation, and obesity management. Many of these genes show higher or lower copy numbers in both *Hn* and *Hs* compared to modern humans, indicating that different levels of gene expression were likely required in these population.

### Two independent classifications lead to the same signal of CNV trends

The clear separation of *Hn* individuals from *Hs* individuals in the multidimensional spaces defined by our two independent datasets of diet-related genes—categorized as “strictly” and “accessory” diet-related genes—strongly suggests that dietary preferences and adaptations are encoded at the genomic level, even when considering small gene sets comprising only a few dozen genes. Notably, both gene sets also distinguish ancient populations from the modern reference genome, which occupies a separate region in the multidimensional spaces, suggesting a significant shift in dietary patterns in modern humans. Importantly, our computational approach opens new perspectives on the possibility to reconstruct the evolution of the human diet by analysing genomic data over the past 50,000 year, extending evidence of the inclusion of starchy foods in *Hs’* diet well before crops domestication. This aligns with the view that the transition from hunting-gathering to domestication was occurring across protracted conditions, shaped by ecological and behavioural factors that left their mark at the population level rather than on individuals.

### Genomically distinct metabolisms in ancient human populations

Due to limited number of available genomes where the estimation of CNV was possible (20), we mainly focused on *Hn* and *Hs*, with inferences for *Den* where possible. The different CNV trends observed for *Hn* and *Hs* reflect that these hominins relied on different diets, and align well with their evolutionary histories. The genes involved in efficient digestion and **metabolism of dietary carbohydrates** (*AMY1A/B/C, SULT1A3/4, SLX1A* and *UGT2B28*) appear important for *Hs* whereas most genes involved in **lipid metabolism** (*CLPS, CLPSL1, RFPL4A/L1, F7, PDX1, WFS1, H2AC11*) show positive trends for both *Hn* and *Hs*. A few genes involved in lipid metabolism (*LPA, ZNG1E/F/C*, and *ACOT1*) exhibit trends specific to *Hn*, while others (*ACOX3*, *HAND1*) are unique to *Hs*. Conversely, the MST1 gene involved in **liver lipid and brown fat metabolism**, so important for the survival of newborns, appears to be of importance for both *Hn* and early *Hs*, both displaying a positive trend compared to hg38. These CNV trends provide key insights into ancient populations, but integrating them with modern genomic and clinical data allows us to deepen our understanding.

Comparing CNV trends in ancient and modern populations reveals two primary distribution patterns, clearly visible in **Figure 3**: in one, both modern and ancient individuals display limited CNV variation, with modern individuals having a single copy, while ancient individuals have higher copy numbers (*F7, PDX1, WFS1, H2AC11, ACOX3, HAND1*). In the second pattern, *Hs* and/or *Hn* exhibit narrow CNV variation (*ZNG1E/F/C, ACOT1, CLPS1, LPA*), while modern individuals show a broader CNV range. The *CLPS* gene deviates from these patterns, showing a wide CNV range in both modern and ancient populations: modern individuals range from 1 to 3 copies, mostly centered around 1 and 2, whereas *Hn* and *Hs* range from 1 to 4 copies, centered around 2 copies for *Hs* and 3 for *Hn*. The narrow CNV range observed in *Hn* across most genes confirms once more their great homogeneity. Overall, the elevated CNV in archaic individuals aligns with the role of these genes in enhancing nutrient processing and the ability to consume high-fat diets, as seen with *ACOX3* and *PDX1* (**Supplementary Text 3**).

The functional effects of CNV in modern humans—whether their copy numbers match those found in ancient genomes or their expression is altered by environmental factors such as high-fat diets—enable us to hypothesize about their adaptive significance in ancient populations. By investigating CNVs in genes involved in key metabolic processes, we can trace how these variations shaped dietary strategies and survival mechanisms across diverse environments.

**Carbohydrate metabolism** played a pivotal role in the dietary adaptations of *Hs*, as evidenced by the *AMY1A/B/C* genes. Here, broadening the number of genomes investigated since the pivotal studies by Perry and co-authors (4,5) provides further evidence that *AMY1A/B/C* CNV played a vital role in equipping *Hs* to digest and transform dietary carbohydrates and thus benefit from caloric energy for cost-efficient homeostasis. It is relevant here to estimate when the *AMY1A/B/C* duplication occurred (9,10) as the earliest *Hs* in Eurasia, represented in this work by the Ust’-Ishim fossil, already bore three copies per haploid genome. An ancient *AMY1A/B/C* duplication matches the archaeological evidence and provides molecular corroboration that cooked rhizomes could have been consumed as early as 170 ybp in South African caves (58), dovetailing with the occurrence of ground stones used to mechanically tenderize rhizomes and tubers during the Early Upper Paleolithic in Eurasia (15,59–62). Based on our data, it is parsimonious to consider that the emergence of an efficient strategy for starch digestion and metabolism is related to the *Hs* lineage already carrying the *AMY1A/B/C* gene duplication when they moved out of Africa, thus well before crop domestication (9,19). On the other hand, our findings reveal that the earliest *Hn*, such as the Altai Neandertal D5 (dated to 120,000 years ago) and Den D3 (dated to 76,000–51,000 years ago), possess only two copies of the *AMY1* gene (see **Tables 1A** and **S6**). Among the eight *Hn* individuals analyzed, all have either one or two copies. This confirms once again the great homogeneity of *Hn*, which has already been demonstrated many times from an anatomical point of view in terms of bone and dental characters, and suggests that this character was also present in *Den*. The only exception is Les Cottès (dated to 43,000 years ago), which has three copies—potentially due to interbreeding with *Hs*, who may have introduced the third copy, as *Hs* were already present in Europe at that time. As shown in **Figure 3**, the CNV distribution in ancient and modern Sapiens spans a broader range of 1 to more than 5 copies across populations. In ancient individuals, the three copies were already present in the earliest *Hs*, with higher copy numbers becoming more prevalent in later Mesolithic individuals. All five modern populations exhibit a wide range of CNV variation. This stands in sharp contrast to the limited variation in *Hn* and *Den*, where the CNV is consistently concentrated between 1 and 2 copies.

A similar pattern of variation is observed in the *SULT1A3/4* and *SLX1A/B* gene clusters, where *Hn* display only one copy, while modern and ancient *Hs* show a broader CNV range. This indicates that gene duplications likely occurred after the EAH lineages split (9,19). We suggest that these duplications may have been driven by the nutrition changes faced by *Hs* in African territories compared to the temperate and cold environments where *Hn* and *Den* foraged their prey. The CNV of these genes may have made efficient lipid metabolism possible as a response to the cold stress during MIS 3 and moderating susceptibility to ketosis in adults (17)(p. 256). *SULT1A3/4*, involved in carbohydrate metabolism and in thyroid hormone activity, may have contributed to this adaptive trend by increasing cold-induced thermogenesis, to avoid the effects of “polar T_3_ syndrome” (63). The involvement of thyroid hormone in vital pathways warrants further investigation in relation to *SULT1A3/4* gene cluster emergence, chronology and duplication. Overall, CNV in genes involved in oxygen and lipid metabolism provides valuable insights into adaptation to temperate and cold ecosystems. This evidence can help deduce or confirm thermal ranges and their implications for differential nutritional strategies among the three populations.

Genes *MST1* and *H1.2*, which are involved in **liver lipid and brown fat metabolism**, play essential roles in regulating adipose tissue function, crucial for thermogenesis and energy balance. *MST1*, also known as *STK4*, protects the liver from damage due to fasting or high-fat diets, stimulates adipose tissue differentiation (64), and importantly, acts as a physiological suppressor of mitochondrial capacity in brown (BAT), beige, and white (WAT) adipose tissues. Genetic inactivation of *MST1* increases mitochondrial mass and function, providing resistance to metabolic dysfunction induced by high-fat diets (65). These results suggest that, in archaic humans, increased *MST1* CNVs likely enhanced mitochondrial efficiency and metabolic resilience in challenging environmental conditions. *MST1* duplications are consistently observed in Den D3, and across *Hn* and *Hs* genomes, underscoring their significance in metabolic adaptation. On the other hand, the *H1.2* gene, a linker histone variant, regulates the browning of WAT and thermogenic gene expression in beige and brown fat cells, particularly in response to cold environments (66). Increased *H1.2* copy numbers in archaic humans likely provided a finely tuned mechanism for energy storage and release, beneficial for cold survival (66). Notably, higher copy number of H1.2 is seen in all *Hs* genomes except for the oldest out-of-Africa Ust’Ishim genome, indicating that these duplications became more widespread as *Hs* adapted to colder Eurasian climates. Additional genomic changes in *Hs,* such as *CD24* loss, associated with a reduction of WAT (67), *TCAF2* diversification, believed to be a result of positive selection in response to cold climates or dietary adaptations (68), and *MST1* duplication, which also plays a role in physiological responses to oxygen depletion (69) (see **Supplementary Text 3**), further support enhanced brown fat metabolism, providing flexibility and resilience in diverse environments.

### Expanding Nutritional Insights: high-fat diet and nutrient utilisation in *Hn*, *Den* and *Hs*

New genetic evidence identified in this work (**Figure 3**) sheds light on the metabolic adaptations of *Hn*, *Den*, and *Hs* to high-fat diets and nutrient utilization, challenging established perspectives. Genes associated with micronutrient metabolism, including vitamins and minerals (70), exhibit notable duplications in both *Hn* and *Hs*. While animal products and oysters are the richest sources of zinc (71), plant-based sources such as pine nuts, hemp seeds, sesame seeds, chickpeas, and lentils, which were accessible to both *Hn* and *Hs*, also provide significant amounts of zinc. The *RFPL4A/L1* gene cluster produces a protein that binds zinc, and the *ZNG1E/F/C* cluster, which is vital for zinc transfer, exhibits a notable trend of increased CNV in *Hn* and certain *Hs*. Calcium is another important example of a food-related minerals. *WFS1* encodes wolframin, a protein believed to regulate cellular calcium levels, which is essential for various critical cellular functions. The association of *WFS1* with diet, particularly high-fat diets, has been demonstrated (72), showing that having only one functional copy of this gene can cause metabolic impairments. The presence of multiple copies of *WFS1* may have provided an advantage to individuals such as *Hn* and *Hs*, who followed high-fat dietary regimes.

### New information on the *Hn* diet

Our work highlights that *Hn* metabolism was more efficient in the digestion of lipids than carbohydrates. The genetic adaptation to fat metabolism is an archaic pattern present in *Hn*, who were adapted to temperate and cold climates. The CNV trend of genes involved in fat metabolism (**Table 1**) undoubtedly emerged under cooler climatic conditions (even before the timing of Altai Neandertal D5, the oldest Neandertal in our sample) when the *Hn* differentiated. As already pointed out for *Hn* anatomical characteristics (i.e., their low stature resulting from adaptation to the cold) (13,73,74) and their cold adaptation genes (75,76), these adapted traits persisted even during interglacial periods, putatively due to the low genetic diversity of *Hn* (31,34). We found a trend in *Hn,* going from the oldest Altai Neandertal to the later Spy 94a individual, confirming independently what has already been noted from fossil analysis — that Altai Neandertals were highly carnivorous (77–79) while Spy 94a might have added proteins of vegetable origin to his diet (80). The CNV population trend of genes involved in fat and protein metabolism could even explain the peculiar skeletal morphology observed in *Hn*. A large bell-shaped lower thorax and a wide pelvis shaped to accommodate the larger liver and kidneys are an adaptation to a high fat, high protein diet. These organs are highly solicited for lipid and protein metabolism so the genetic evidence coincides with the specific anatomy (81) and high energy needs of *Hn* (13,82).

*Hn* shows a negative trend for genes involved in dietary carbohydrate digestion compared to the modern human reference genome and *Hs*. Nonetheless, the late *Hn* in our sample, Les Cottés ZA-1514, has 3 copies of AMY1A/B/C like the human reference genome, indicative of a less restricted foraging strategy being adopted by this late *Hn* due to the different environmental conditions (8,12,15,16,18,83). A plausible interpretation is that some late *Hn* may have interbred with early dispersals of *Hs*. Further analyses will be feasible once additional high-coverage genomes from southern or coastal *Hn* become available for study.

*CLPS, CLPSL1*, and *LPA* positive CNV trends in *Hn* have been reported to relate to fasting hepatic FA metabolism because they control the lipids balancing oxidative flux and capacity (84). Fats can be digested to form FA, and proteins can be partially transformed into glucose through gluconeogenesis (85,86). Gluconeogenesis is an efficient pathway to allow glucose to directly entering the bloodstream, to fuel the high demand of organs, notably the brain, which in *Hn* is very large (around 1450 cc, much larger than the average brain today) (13). Although gluconeogenesis demands much energy (87), within a highly carnivorous regime it can (i) provide glucose to the brain for a limited time, and (ii) become a profitable pathway to maximize the benefit of protein intake (3) by effectively reducing the amount of nitrogen to avoid protein poisoning (3,85,88) so limiting ketosis in adults (8).

Regarding the beneficial role of FA in the diet, it is well known that nuts have always been present in human diet as shown by the Acheullian site in Israel 700 kyrs ago (89). Yet pine nuts were also available in the Altai Mountains refugia. In certain geographical regions (e.g coastal Mediterranean), it is possible that *Hn* relied on a diet much richer in omega-3 present in wild fish (90). FA are consumed at higher levels in diets incorporating more animal proteins and are critical for brain development, brain metabolism, and inflammatory responses (91).

### A glance into the *Den* diet

The CNV analysis was possible on only one *Den* genome. Den D3 shows an *Hn-like* trend, particularly in genes related to lipid digestion, which is expected given their shared ancestry. Indeed, according to genetic estimates, the *Hs* and *Den* lineages separated from the *Hs* lineage some 800 ka ago, before separating from each other around 400 ka ago (30,47). It is acknowledged that *Den* in Altai are a western extension of a much larger population originating in central and southwestern Asia (92,93) with a less small size population that *Hn* (92). Further analyses will be possible when more *Den* genomes become available.

### A new picture of the *Hs* diet

If it holds true that plants were part of the hominid diet for millions of years during their tropical forest foraging niche (94), the physiology required for efficiently digesting and metabolizing dietary carbohydrates, particularly starch, took time to adapt, likely through the redesigning of core metabolic and physiological processes (7). This took place during the evolution of *Hs* in Africa, where under favorable environmental conditions, they may have experienced the gathering and transformation of plant organs rich in starch (58,95). Our study provides three main contributions linked to dietary carbohydrates metabolism. First, *Hs* and the modern genome hg38 have a higher copy number for the gene *UGT2B28*, and the *AMY1A/B/C, SULT1A3/4* and *SLX1A/B* gene clusters compared to *Hn* and *Den*. Second, our data are consistent with the proposed early emergence of *AMY1* coupling – between 273-475 ybp – therefore after the divergence from *Hn* and well before crop domestication (9,10,19). Third, we identified the role of the *SULT1A3/4* and *SLX1A* gene clusters and of gene *UGT2B28* as main contributing factors explaining the first two dimensions of the CNV space of individuals in **Figure 4**. These three results prompt new questions centered on the importance of dietary carbohydrates in the diet of archaic humans and, in particular, in *Hs*.

### From diet to survival: broader roles of key genes

Lastly, it is important to highlight three genes— *HAND1, BOLA2/2B*, and *F7*—that extend beyond diet and play significant roles in reproduction and survival. *HAND1*, which is duplicated in most *Hs*, has been shown to inhibit the conversion of cholesterol to steroids in trophoblasts, thus playing a vital role in the development and physiological function of the human placenta (96). *BOLA2/2B*, involved in the immune response in the gut, also plays a role in the maturation of cytosolic iron-sulfur proteins. Its duplication in *Hs* alters iron homeostasis, potentially serving an adaptive function by protecting against iron deficiency, which is crucial during embryonic development (97). Further investigation of these genes could provide valuable insights into the interplay between diet and reproduction in the evolution of the *Homo* genus. Together, these genes suggest a complex interplay between diet and reproduction in the evolution of the *Homo* genus, warranting further investigation.

Additionally, the coagulation factor *F7* gene, which is produced in the liver and is vitamin-K dependent, encodes an enzyme of the serine protease class that plays a critical role in the blood coagulation system, facilitating the formation of blood clots. This gene is duplicated in all archaic individuals including *Hn*, *Den* and *Hs*. The duplication of *F7* induces fast blood coagulation, which could have been advantageous for these populations by enabling rapid wound healing and reducing the risk of infection, thereby increasing their chances of survival (75). Additionally, *F7* is also associated with dietary fat intake and cholesterol levels (98), with studies documenting its activity being influenced by diet (99). These findings underscore the broader adaptive roles of these genes, linking nutrition, reproduction, and survival strategies in archaic human populations.

### Conclusion

The different CNV trends observed for *Hn* and *Hs* (and *Den*) reflect that these hominins relied on different sources of energy in their diets. Our unsupervised analysis of the CNV of multiple genes adds direct lines of evidence that confirm the biological adaptation of *Hn* to a narrow-spectrum diet and highlight the genetic changes facilitating the liver lipid pathway in both *Hn* and *Hs* living in the same nutritional environment of the cold Eurasia. The increasing duplication of *AMY1A/B/C*, *SULT1A3/4* and *SLX1A/B* genes evident in *Hs* is consistent with broad-based foraging strategies grounded, on the behavioral side, by the intentional transformation of starch-rich storage organs through the use of grounding stones, and this occurred much before crops domestication (59), started with the Neolithic (61,62). Different nutritional strategies among the three human populations inhabiting the temperate and cold latitudes during the crucial timing of their overlapping could have contributed to *Hs* better fitness leading to a demographic advantage that allowed them to conquer Eurasia and then the world.

## Materials and Methods

**An unsupervised procedure for the identification of genes involved in digestion presenting a CNV trend within an ancient population.** The methodology (**Figure 2**) is based on several distinct computational steps:

1. Selection of all available genomes that could allow for an estimation of the haploid gene copy number.
2. Mapping of the selected ancient genomes to the modern human reference genome GRCh38/hg38 (42).
3. Clustering of recent paralogs and estimation of the haploid gene copy number for genes of the genomes identified in 1.
4. Identification of genes whose number of haploid copies in *Hn*, *Hs* or *Den* is higher or lower than in the modern reference genome GRCh38/hg38 for more than the half of individuals in the population. This means that a selected gene should display a *differential CNV* for at least 5 individuals in *Hn* or at least 4 individuals in *Hs*. This step identified 781 genes exhibiting population-dependent variation in copy number.
5. Selection of a subset of genes based on Human Protein Atlas tissue expression data and information on functional annotation in UniProt.
6. Comparison of CNV data from 28 diverse modern human ethnic groups to evaluate differences with ancient populations.
7. Gene Ontology (GO) and literature-based analysis of the final set of genes for a functional characterisation.

Steps 1-3 focus on read mapping and data preparation for the 19,988 genes identified across 15 ancient genomes. In steps 4-6, the dataset is refined by filtering the 19,988 genes down to 50 genes and gene clusters. Finally, step 7 links the selected genes to their functional characterization. Full details of the procedure are provided in **Supplementary Text 2**.

**Figure 5A** illustrates the analysis of the AMY1A/B/C gene cluster across modern and ancient individuals (see **Figure S1** for an extended analysis). In modern populations, *AMY1A/B/C* is duplicated, with a median of three haploid copies. When considering diploid CNV estimations, the distribution of pair and unpair diploid copy number aligns with prior analyses (54) of modern genomes, as illustrated in Figure 5B. CNV estimation of AMY1A/B/C of each genome is derived by summing the average read coverage over each AMY1 gene copy in hg38. Figure 5C displays this read coverage across each copy for the *Hs* genome of Ust’Ishim and the *Hn* genomes of Vindija 33.19 and Mezmaiskaya 2 (see **Figure S8** for mapping on all ancient genomes). Ust’Ishim and Vindija 33.19 are high-coverage genomes, showing less sensitivity to mapping fluctuations compared to the lower-coverage Mezmaiskaya 2 genome.

**Figure 5.**
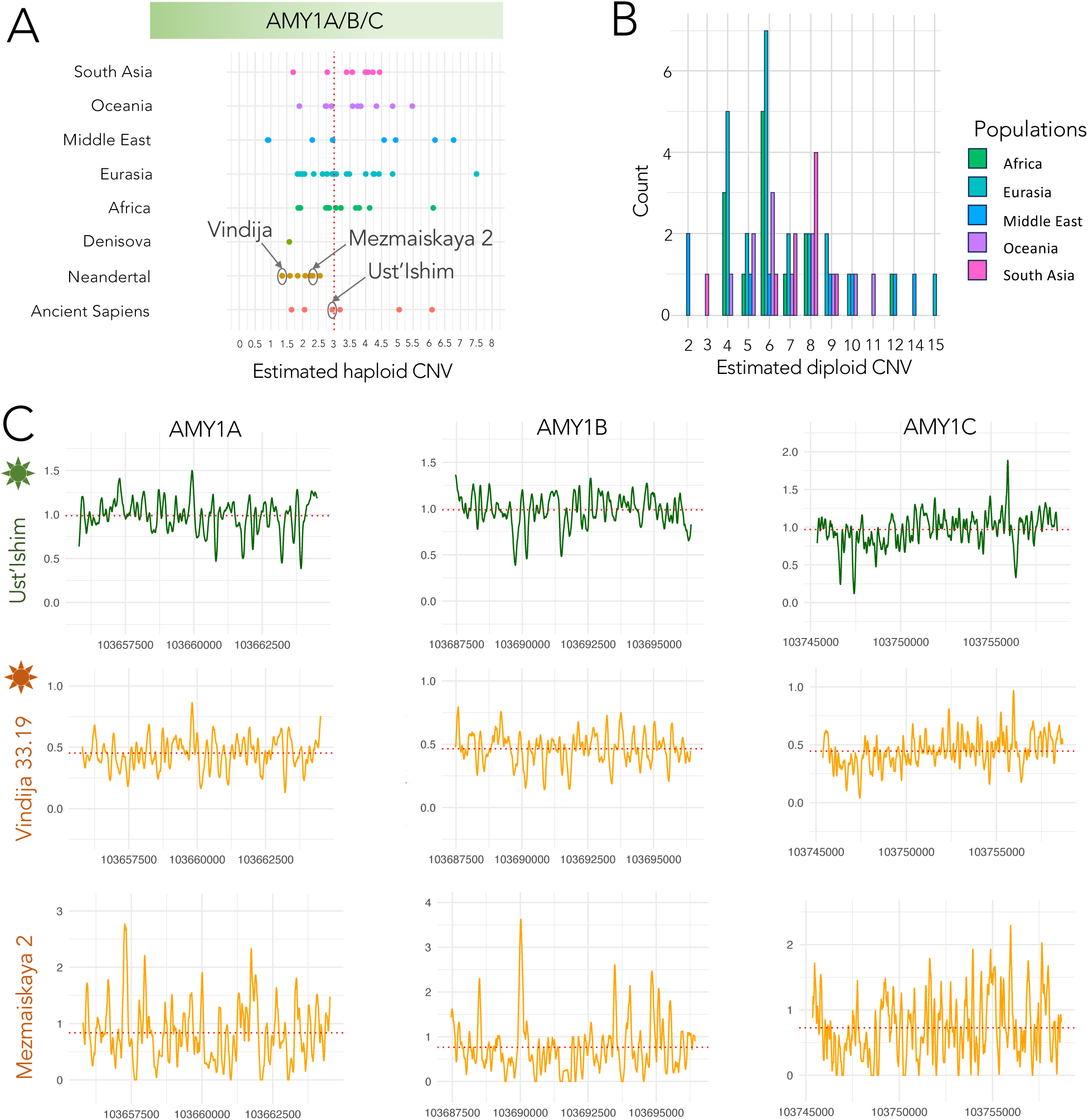
Read mapping analysis for the AMY1A/B/C gene cluster. **A.** Comparison of the estimated haploid CNV of the AMY1A/B/C gene cluster across all ancient and modern individuals in this study. See **Figure S1** for an extended analysis including genes AMY2A and AMY2B. **B.** Estimated diploid CNV for the modern populations considered in A. See also **Figure S1**. **C.** Read mapping for genes AMY1A (left column), AMY1B (center column) and AMY1C (right column) for Ust’Ishim (top row), Vindija 33.19 (middle row) and Mezmaiskaya 2 (bottom row). Stars for Ust’Ishim and Vindija 33.19 indicate high-coverage sequencing. For further analysis on all ancient genomes, see **Figure S8**.

After the first four steps identifying the genes with differential CNV, we used the notion of *CNV trend* to analyze them within more than one population. Formally, a gene has a positive/negative *CNV trend* in a population if its gene copy number is either greater/smaller than that of the modern reference genome for more than the half of the individuals in the population (5 of the 8 *Hn* genomes, 4 of the *Hs* 6 genomes). This criterion is aimed to highlight the tendency for a gene to have more or fewer CNVs than the modern reference genome GRCh38/hg38 within a population. The gene MST1, for instance, displays positive CNV trends for both *Hn* and *Hs*. The gene cluster AMY1A/B/C was selected because of its negative differential CNV in *Hn*, but in the *Hs* population there is no trend that is picked up by the CNV trend criterion. Also, the gene ZNG1E/F/C displays a positive trend in *Hn* and no specific trend for *Hs*. **Table 1**, **Table 2**, and **Table S6** show the aforementioned genes, their estimated copy number (rounded and not, respectively) for the 15 ancient genomes and the CNV trends for the two populations. The analysis on the Denisova genome is carried out for comparison.

### Selection of ancient genomes for genome-wide analysis

Among all publicly available human ancient genomes, we identified all genomes presenting a minimum number of features that allowed us to properly perform a CNV analysis (see **Table S1**). Specifically, we considered genomes that:

1. present a minimum of 1x coverage of the human reference sequence, with available alignment;
2. have been affected less than 5% by contamination from modern human sequences;
3. retained sequences aligned with repetitive regions (indicated by a MAPQ value of 0).

The three conditions allowed 15 archaic human genomes to be considered. They were retrieved from accession codes published in manuscripts before December 2020, to which we added the more recent genome of Peştera Muierii (July 2021); see **Table S1** for downloading information.

Reads from these 15 genomes were remapped to GRCh38/hg38. We studied 18,043 genes/gene clusters (see below) which are covered by the 15 genomes and are shared with the modern reference human genomes hg38.

It is worth mentioning that some very interesting genomes, although characterized by a sufficiently high coverage, were discarded for various reasons: the available alignment files for Anzick-1, Kostenki 14, Sunghir 1--4, and Sunghir 6 do not comprise repeated regions; LaBraña lacks data for chromosome 22; Den D2, Hohlenstein Stadel, Scladina, San Teodoro 3, San Teodoro 5, and Zlatý kůň have low coverage and/or high contamination.

### Selection of modern genomes for genome-wide analysis

To compare ancient and modern genomes, we selected 64 individuals from The International Genome Sample Resource (IGSR) database (40,41,100) of The 1000 Genomes Project (101), representing ethnic groups from Africa (13 individuals), Eurasia (22), the Middle East (8), South Asia (9), and Oceania (12) (refer to Figure 1 and **Table S3)**. The sequence reads from these genomes were previously aligned to the hg38 reference genome. CNV analysis for these modern genomes was performed using the same pipeline as for the ancient genomes (see **Supplementary Text 2**).

### Technical warning

The quality of sequencing data plays a crucial role in read depth analysis and in the copy number determination that can differ in precision (53,102,103). Indeed, human genome assembly might not fully map highly complex genomic regions raising potential doubts on the correct estimation of gene copy numbers in human populations. By choosing GRCh38 genome as the current human reference, we guarantee the latest accessible estimation to gene copy numbers in the reference genome.

## Acknowledgments

We thank Alissa Mittnik (Johannes Krause lab) for sharing her estimate of contamination of the Motala 12 nuclear genome.

## Funding

This work has been partially supported by:

- ANR grant STARCH4SAPIENS – grant ANR-20-CE03-0002 (AC, SC, LL);
- LabEx CALSIMLAB grant ANR-11-LABX-0037-01 - constituting part of the “Investissements d’Avenir” program grant ANR-11-IDEX-0004-02 (AC, RV);
- SUG grant M4081669.090 (NTU, Singapore) (LL);
- PHC Merlion, 2018, Singapore-France bilateral agreement (NTU, Singapore) (LL, SC).

## Author contributions

Conceptualization: AC, LL, SC

Methodology: AC, RV

Investigation: AC, LL, SC, RV, LP, NR

Visualization: AC, RV, LP, NR

Funding acquisition: AC, LL, SC

Supervision: AC

Writing – original draft: AC, LL

Writing – review & editing: AC, LL, SC

Writing – SOM: AC, LL, SC, RV, NR, LP

All the authors discussed the results and agree on the final version.

## Competing interests

Authors declare that they have no competing interests.

## Data and materials availability

All data are available in the main text or the supplementary materials.

## Supplementary Text 1

### S1. Presentation of “archaic humans”

**Neandertals (*Hn)*** are a human hunter-gatherer fossil population that lived in Eurasia between 300 kya and 40k ybp. They were totally replaced throughout their territory by us, *Homo sapiens*. Paleoanthropologists have shown that the evolution of this fossil population could be originated from *Homo heidelbergensis* in Europe, back at least from 450k ybp, through the identification and progressive accumulation over time of typical *Hn* morphological features (104).

In Africa H*omo heidelbergensis* gave rise to *Homo sapiens* (*Hs*) whose oldest representatives, from fossil remains, are identified around 300 Ka (105). Thus, while *Hn* was differentiating in Europe, *Hs* was differentiating in Africa. These data are consistent with the genetic ones (106). By analysing the genomes of *Hn* and of modern humans (*Hs*) it was estimated that the two ancestral populations separated between 220 000 and 470 000 years ago (107). This splitting time is highly relevant for the emergence of the *AMY1A/B/C* gene cluster (9) and it is relevant for the reasoning presented in this paper. According to some studies, this timeframe would seem to match that of the extinct species *Homo heidelbergensis*, which has been found in Africa, Europe, and possibly Asia (108). The fact that only present-day Europeans and Asians possess this *Hn* genetic capital indicates that interbreeding must have taken place just after the first *Hs* left Africa to conquer Eurasia, which the researchers estimate to occur in several waves starting approximately 100,000 years ago (47,109). This hybridation has been since confirmed by several analyses (the last being reported in (35)), which succeeded in sequencing the genomes of five *Hn*, including four late *Hn* from Western Europe - one from France, two from Belgium and one from Croatia - and an older one from the Russian Caucasus. They also show that the *Hn* genes of present-day Europeans are not close to those of late European *Hn*. Both this genetic distance and the enigmatic absence of *Hs* genes in late *Hn* can then be explained if one’s makes - and validate - two hypotheses: on the one hand, that interbreeding mainly induced gene flows in the direction of *Hn* → *Hs*; on the other hand, that the main hybridization event occurred outside Europe well before the arrival of our *Hs* ancestors on this continent (106,107,109).

The European *Hn*, for which extensive morphometric data are available, underwent four glacial events and thus progressively adapted to colder environments (13,110). It is therefore under particular climatic and environmental conditions, with alternating glacial and interglacial phases, that the evolution of *Hn* took place in Eurasia. The climate had important consequences for their adaptability, density and therefore diversity, as well as for their possibilities of expansion and contact with their neighbouring population (111). The *Hn* populations have experienced few bottlenecks, therefore they have never been large in size, structured in small interconnected groups (about 20 individuals) that never outnumbered 70,000 individuals at the time of their “golden age” (during the “Eemian” interglacial at MIS 5, around 120k ybp) (112). The small size of the population has to be correlated with the partial geographic isolation of *Hn* caused by European climatic fluctuations during Pleistocene.

It was around this time (circa 100k ybp) that the European *Hn*, well identified by singular morphological features, spread eastwards carrying the same lithic assemblages and technology as far east as the Altai mountains (113) where they encountered the Denisova (*Den)*. Genetic data has also pointed out low genetic diversity in *Hn*, with a demographic depression peak in the Altai, where a very consanguineous individual has been found (31), and a genetic continuity across Europe from 120k ybp until the disappearance of the population around 40k ybp (114). This low variability is also visible in the morphology of the *Hn*, which remained the same during the last 100k of their existence throughout their broad territory from the Atlantic to the Altai (104) (see SOM p.15 in (113)).

### The expansion of *Hs* and the demise of *Hn*

After the great dispersion of *Hs* from Africa into Eurasia about 60,000 years ago (1), the other archaic humans (EAHs), notably the *Hn* and *Den*, who populated this geographical region, disappeared. The reasons for the success of *Hs* are debated among specialists. The most commonly accepted hypothesis is that EAHs would have competed with *Hs* for food resources during the climatic changes that affected Eurasia (104,115).

The replacement of EAHs, *Hn* in particular, would have been favoured by *Hs*’ greater technical skills (116), their greater cognitive abilities (116–120) and lower social capacities and network (116,121,122).

However, all these hypotheses have been challenged (123,124) as also those suggesting that the disappearance of EAHs was brought about by violent confrontations between the two populations (125) and by the exposure to new infectious agents (126,127).

The great deal of attention paid to behaviourally and culturally driven reasoning somehow overwhealmed the reasoning on the biological complexity of physiological adaptation (110,128) and comparatively little attention was paid to the metabobolic and physiological processes ruling the bioenergetic requirements (8,129–131), the role of microbiomes (128,132) and how to transform different foods into nutrients (133,133). Our analysis add fresh data to the genomic architecture across EAHs suggesting a selective pressure on Neandertal, and possibly on *Den*, towards lipid-rich food to respond to the cold climatic regimes at the boreal latitudes. Conversely, our results support an adaptation to a diet rich in glycemic carbohydrates for *Hs*, evidenced by the old duplication of *AMY1A/B/C* gene cluster (9).

For our CNV analysis, 8 *Hn* genomes are available: 2 from Asia (in Altai: Altai Neandertal D5 and Chagyrskaya 8) and 6 from Europe (in Russia: Mezmaiskaya 1, Mezmaiskaya 2; in Croatia: Vindija 33.19; in Belgium: Spy 94a, Les Cottés Z4 1514, Goyet Q56.1).

**Denisovan, *Den*,** on the other hand, are also an extinct human population but their “discovery” in 2010 is mainly due to mitochondrial (mt) and nuclear (n) DNA studies (47,134–136) extracted from the collagen of bones so fragmentary that they did not allow paleoanthropologists to identify their morphological and anatomical characteristics. *Den* are considered an Asian sister group of *Hn* that have evolved in Asia from *Homo heidelbergensis* after the expansion from Africa and the split with the African ancestors. This African origin has been confirmed and reinforced by genetic studies (from 2010 up today), that have also contributed to date the time of divergence between the African lineage, that will evolve into *Hs,* and the Eurasians’ one that will give rise to *Hn* in Europe and to *Den* in Asia. The analysis of the nuclear genome suggests a recent common ancestor between European *Hn* (notably with Vindija, Vi 33-19) and *Den* dating between 440 to 390 ka ago (34) and thus, situated the *Den* as a sister group to *Hn* (30,34,137) and *Hs*.

The exact geographic repartition of *Den* is not yet clear but available data supports that the Altai is the western edge of their territory that could likely expand to North and South East Asia, until Papua and possibly the Sonda archipelago. *Den,* originally known only from the Denisova cave in the Siberian Altai, has been recognized even in the incomplete hemi-mandible from Baishiya cave (Tibet), on the basis of proteomic study (138) and from DNA extracted from the sediments (139), providing evidence for their occupation of the cave perhaps until 45 ka, when early dispersal of *Hs* was already roaming into Asia.

Probably, *Den* occupied a large part of continental and southern regions of Asia (93,140–143). Indeed, it is in modern Asian populations that the highest percentage of *Den* genes are found (144,145). *Hn* and *Den* have interacted in the Altai area and they occupied successively and perhaps even together Denisova Cave, as witnessed by the Den D11 individual (118.1-79.3k ybp) that had a *Den* father and an *Hn* mother (134,146). The characteristics of the Den D3 genome enable our CNV analysis.

### S1.1 The *Hs* peopling of Europe seen from paleoanthropological and genomic analyses

From archeological, paleoanthropological and genetic studies the European *Hs* peopling show that the ancestry of modern Europeans is quite complex (147). In western Eurasia five dispersals of *Hs* arrival can be identified:

1. An “Aurignacian dispersal” of *Hs*. This arrival is supported by the presence in Eastern and Central Europe of archeological evidence for “Aurignacian flaked industries “ resembling those of the Levant, e.g. Ksar Akil, Lebanon, dating about 43k ybp (148) (see also the recent debate in (149)). These cultural assemblages, that lasted in western Europe until around 31k ybp are always associated to *Hs*, although at the time of modern humans’ arrival *Hn* was still present in Europe (150) making it possible that hybridation have occurred (151). Although for this early phase of dispersal no genomes are available that belong to it with certainty, in this study we analyzed the Peștera Muierii genome. Other genomes from early *Hs* individuals like Kostenki 14, dating to 38k ybp (152,153), Bacho Kiro, dating 45-58k ybp (151) and Zlatý kůň, 45-43k ybp (154) could not be included in our analysis due to genome sequencing characteristics not allowing for CNV analysis (see **Methods**). For this earliest dispersal we analyzed the Ust-Ishim genome (> 45k ybb).
2. *Hs* belonging to the Gravettian assemblages. The Gravettian population seems to have formed a homogeneous biological group despite their chronological and geographical extension that covers all Europe over about 10.000 years. The oldest Gravettian occurrences are dating from around 31k ybp, when *Hn* had already disappeared, until 23k ybp, hence Peștera Muierii 1 and Mal’ta child genomes fall in this group. These fossil populations of hunter-gatherers developed a meta-culture famous for its flourishing art, as well as some emblematic archaeological sites including the Cro-Magnon shelter discovered in 1868 (Dordogne, France) that gave the name with which modern humans are aknowledged. Unfortunately, all the genomes available from this span time (notably Sungir, Kostenki I, Dolni Vestonice, Pavlov, Krems-Wachtberg) do not have the genome sequencing characteristics allowing for CNV analysis (see **Methods**).
3. A new arrival of an unidentified *Hs* population of hunter-gatherers around 14k ybp, probably coming from Anatolia (155,156). This new arrival expanded in Europe during the last Ice Age *Hs* experienced an important demographic bottleneck. Therefore, there is genetic evidence for discontinuities in the European peopling before and after last glacial maximum. For this span time no available *Hs* genomes are worth to match the criteria set for our CNV analysis.
4. Anatolian *Hs* farmers, who spread into Europe about 8,500 years ago, via the Mediterranean coast and the Danube valley (38). In our study, we analyzed the CNV of genomes from late Mesolithic *Hs*, namely Loschbour (Belgium) and Motala 12 (Sweeden) and from an early farmer from Stuttgart, all living in Europe after the great bottleneck that followed the last glacial event.
5. A nomad population from the steppes of southern Russia, who migrated to Europe and East and South Asia about 5,000 years ago (39,157) which is beyond the scope of the present analysis.

### S2 Presentation of the fifteen re-analyzed probands

In recent years, the availability of good quality nuclear DNA from various human fossils has increased the potential of reconstructing the hominins phylogeny and their interbreeding (158). Also, the availability of hundreds of thousands of complete human genomes from different modern populations allows the reconstruction of maps of single nucleotide polymorphisms and structural variants. Among all the ancient DNA genome sequences from *Hn* and *Den* publically available to date for our unsupervised analysis (this study was finalised in 2024), we selected only those with high-quality genomes (see **Methods**) namely one *Den* (D3) and eight *Hn (*Altai Neandertal D5, Chagyrskaya 8, Mezmaiskaya 1, Spy 94a, Vindija 33.19, Mezmaiskaya 2, Les Cottés Z4 1514, Goyet Q56.1). These probands are representative of the two archaic human (*Hn* and *Den*) that spanned over 50,000 years of the Late Pleistocene and approximately 8,000 km across Eurasia. We also analysed three *early Hs* genomes (Ust’Ishim, Peștera Muierii 1, Mal’ta), two late hunter-gatherers and one early farmer from Europe *Hs* genomes (Loschbour, Motala 12, Stuttgart). In all, 15 genomes were selected for the CNV study (**Table S1**).

### S2.1 Denisovan (*Den*)

#### Den D3 (Denisova cave, Altai Mountain, Central Siberia)

Den D3 (“pinky”), is a distal phalanx of the fifth finger belonging to an adolescent female individual. This is the first fossil attributed to a new fossil population on genetic ground (30,47,134,135). It was unearthed in layer 11.2 square D2 of the East Gallery at Denisova cave (51.40N, 84.68E) during 2008 field season. It has recently been dated to 69–48k ybp from optical dating of the associated sediments and to 76.2–51.6 kybp using a Bayesian modelling approach that combines chronometric (radiocarbon, uranium-series and optical ages), stratigraphic and genetic information to estimate ages for the hominin fossils at the site (159). Den D3 lived approximately at the same time as *Hn* Chagyrskaya 8 (32) or slightly before, thus sharing similar environmental conditions.

### S2.2 Neandertals (*Hn*)

**Altai Neandertal**, also known as **D5 (Denisova cave, Altai Mountain, Central Siberia)**

Denisova is the name of a cave which is located at a height of 700 m asl dominating a green valley crossed by the river Sibiryachikha. The cave is located at the foot of the Altai Mountains in Siberia, near the border between Russia, Mongolia and Kazakhstan. This is not the only cave or prehistoric site in the region as several *Hn* skeletal remains have been retrieved in the nearby Okladnikov cave and at Chagyrskaya cave, about 100 km away (32).

The Altai *Hn* occupied Denisova cave roughly 110 ka ago (159,160). The Altai Neandertal is represented by a proximal pedal phalanx of a female that furnished a 52-fold genome coverage (31). Several very incomplete fossils from layers 12.3 and 11.4 have also been assigned to *Hn* based on their mitochondrial (161) and nuclear DNA (31). Additional evidence for the presence of *Hn* in the cave comes from analyses of sedimentary DNA from at least 2 layers (11 and 14), which indicate several episodes of *Hn* occupation (146).

### Chagyrskaya 8 (Chagyrskaya cave, Altai Mountains, Central Siberia)

Chagyrskaya is a karst cave discovered in 2007 and it is located in southern Western Siberia, near the Charysh River in the foothills of the Altai Mountains. Since 2008, 74 *Hn* remains, along with fauna and a rich lithic industry, have been unearthed, providing a large and mostly well-preserved collection of morphologically diagnostic *Hn* remains from the Altai mountains. The faunal and pollen remains suggest a milder climate in the Altai foothills that was likely a refugium for *Hn* populations during the late Pleistocene (162–166).

Chagyrskaya 8 is a distal manual phalanx from a *Hn* individual whose genomes have been sequenced to 27-fold genomic coverage (32).

Although the age of Chagyrskaya 8 is constrained by a DNA-based estimate of ∼80 ybp (28) and an optical age for the associated deposits of 59–49 ybp, Chagyrskaya 8 may have lived at around the same time than Den D3 or up to several millennia later. The genome from Chagyrskaya 8 shows that this *Hn* is related to *Hn* from western Eurasia (notably with Vindija 33-19) more than to the Altai *Hn* who lived earlier in Denisova Cave (31,32,34,35). It is possible that distinct groups of Neandertals repeatedly migrated between Western Eurasia and Siberia (113), but it is also possible that Siberian *Hn* lived in relatively isolated populations of less than 60 individuals on the Eastern edge of the *Hn* territory. The long-distance movements from western Europe to Altai of *Hn* is further confirmed by the studies of lithic assemblages – more than 90,000 pieces – resembling to the so-called “Micoquian-like assemblages” observed in Central and Eastern Europe (113). These connections are also supported by a recent study on *Hn* blood system (104). These data make it possible to explain the great homogeneity that is observed in the anatomical characteristics of the *Hn*, which is found throughout their territory from west to east and over the course of time from the earliest to the most recent.

**Mezmaiskaya 1**, **Mezmaiskaya 2 (Mezmaiskaya cave, Krasnodar region, Caucasus, Southern Russia)** Mezmaiskaya Cave is located near the right bank of the Sukhoi Kurdzhips river, in the north-western foothills of the North Caucasus, southern Russia. Among several *Hn* remains retrieved in the cave, the genome extracted from a rib of a still breast-feeding neonate (Mez 1, who died about 2 weeks after birth) dating back to ∼70k ybp was the second *Hn* genome sequenced after the Feldhofer, retrieved in the eponym site, in the early 2000s (167,168). In 2016, a successful extraction from a skull fragment of another infant (Mez 2, 1-2 years old, so putatively still breast-feeding) dated to 44,6–42,96k ybp, revealed an optimal collagen preservation that permitted to sequence the genome (35).

The still ongoing excavation, by yielding human skeletal remains associated with middle Paleolithic assemblages of local facies called “Micoquian tradition” (169), has revealed the key role played by the Caucasus corridor not only for *Hn* but also as one of the gateways for *Hs* into the boreal latitudes.

### Vindija 33.19 (Vindija cave, Northern Croatia, Southern Central Europe)

The Vindija Cave, in the Dinarides Alps, discovered in 1974, is a highly recognized site for the richness in human skeletal remains of both *Hn* and a few *Hs.* Three fragmentary bones of *Hn* were analyzed for the first draft sequence of the Neanderthal genome project (109). In our analysis we consider Vindija 33.19, one of the five *Hn* osseous remains that were recently genotyped (34,35). It belongs to a *Hn* female that lived 48-65k ybp (170) and furnished a 30-fold genome (34). The comparison with the previous *Hn* genomes (109) allowed inferences on the similarity the Neandertal population from Vindija with late *Hn* from western Europe and from the Gibraltar area (114). However, the recently obtained Vi 33-19 genome was compared with those of Chagyrskaya 8 and of the other *Hn* individuals retrieved in that cave, allowing inferences on the western origins of these Altai *Hn*, even considering the technological skills observed in the lithic assemblages (32,113). Several individuals from Croatian site have been also deeply investigated for dental microwear texture analysis and the results suggest that some individuals among the Vindija *Hn* group where consuming soft food like meat at high level (171) and this data are fully in agreement with our results and with the isotopic analysis (172) although anisotropy on teeth wear may suggest some raw fibrous plant food, features that can be also compatible with the use of teeth as a third hand involved in fibres processing.

### Spy I/94a (cave in the Namur province, Belgium)

The site of Spy is well known since the end of 19th century and yielded *Hn* several individuals, highly fragmented. The first two skeletons were discovered in 1886 in Jemeppe-sur-Sambre, province of Namur, Belgium (173,174) and represent the second official discovery of Neandertal fossils ever reported, thirty years after the one in the Neander Valley in Germany (Feldhofer). Since 2010, the re-examination of all the human remains have permitted to identify at least 6 individuals, one juvenile and five adults.

The genome considered in our study, was obtained from Spy 94a upper right molar (M_3_) from a male individual which maxillary fragment including the M_3_ *in situ* was radiocarbon dated to 35,810 + 260, –224 years BP (173,175). The dental calculus analysis of Spy I lower right molar (M_1_) revealed some starch grains, most probably to be attributed to USOs (under storage organs such as water lily (15) which rhizomes are poor in proteins, (175)) and also to grasses. On the basis of these occurrences have been inferred that Neandertals from Spy were including plant food in their diet (15), which is in accordance with the tooth wear-traces observed by (176). Also, tooth wear from Spy I resemble those observed in different individuals from Vindija (but not from Vi 33.19, the specimen in our analysis), Krapina, Hortus and from Kulna 1 (these three latter individuals are not genotyped), and the wear is interpreted as coherent with some plant food consumption (175). By showing two copies of the AMY1 A/B/C gene cluster, our data on Spy I/94a lend poor support to the efficient digestion of starchy food.

However, at present, it is not possible to directly compare our genes CNV trends with the data from other *Hn*/Early *Hs* specimen yielding starch entrapped in dental calculus as reported in recent reviews (175,177). Furthermore, other studies caution about the attribution of plant remains retrieved in dental calculus to the exclusive connection with food consumption (177) and bring about insights on the use of teeth as the “third hand” therefore starch, fibres, tissues residues entrapped in the calculus might also be referred to alternative plants processing (e.g. for technological processing) or for medical purposes (178,179). This economic activity can also serve as an alternative interpretation for the tooth wear recognized on Spy 1(M^2^) previously interpreted as due to course dietary resources (175).

### Goyet Q56–1 (“third cavern” of Goyet cave, Belgium, Central Europe)

The so-called “third cavern” cave, excavated during the 19th century and then re-opened in the 1990s, belongs to a large karstic complex in the Mosan basin, Belgium. It yielded almost 100 human skeletal remains and a rich Middle and Upper Paleolithic industries difficult to disentangle due to the methodology applied by Dupont, one of the pioneers of the studies in Belgian Prehistory. Among the many remains attributed to *Hn*, for the CNV analysis we consider the Goyet Q56-1 genome, obtained from the fragment of the right femur (35) and dated to 40.5–45.5k ybp. The isotopic analysis has been carried out on several fragments of other *Hn* genotyped fragments (Q119-2, Q305-4, Q305-7, according to mtDNA) and the data closely resemble those from other Western and Central Europe hominins like Feldhofer, El Sidron and Vindija (180). The modest genetic variation shown by these genomes is supportive of the low genetic variability typically explained by the limited number of individuals of late *Hn* clades. It has to be mentioned that dental calculus analysis was carried out on Goyet VI *Hn* individual highlighting that this individual was adding some plant food on his prevailing carnivore diet (16). According to our data Goyet Q56-1 – together with Spy I/94a and Les Cottés Z4–1514 (see below) – shows the duplication of the *AMY1 A/B/C* gene cluster lending credence to the digestion of dietary carbohydrates for these individuals. However, carbon isotopic data suggest that other *Hn* individuals living at Goyet cave were heavily relying on mammoth, as it occurred for Spy and Scladina Neandertals (181), which is also a dietary condition supported by our data (high duplication of the genes involved in lipid and protein metabolism).

### Les Cottés Z4–1514 (Les Cottés cave, New Aquitaine, France, Western Europe)

The Cottés cave or Cottets cave is located in Saint-Pierre-de-Maillé, in France. The first excavations and discoveries of Mousterian and Aurignacian occupation levels date back to 1880. Since 2006, new excavations have been carried out, as this cave has the particularity of presenting a continuous archaeological sequence from the end of the Middle Paleolithic to the beginning of the Upper Paleolithic, thus including the Mousterian, the Chatelperronian and the first phases of the Aurignacian. Les Cottés Z4–1514, a *Hn* tooth dated to 43,740–42,720 ybp, has been recently genotyped (35) allowing the CNV analysis. As for the two mentioned Belgian *Hn* individuals, even for Le Cottés Z4–1514 it would be possible to suggest the access to a more mixed diet.

### S 2.3 Homo sapiens (*Hs*)

In the frame of this article, we analysed the CNV of an Early *Hs* Palaeolithic hunter-gatherer from Europe, Peştera de Muierii, and two from Asia, Ust’Ishim (north-western Siberia), at present the oldest *Hs* at the boreal latitudes who lived prior to the split between Western and Eastern Eurasian, and the Mal’ta child, that lived in eastern Siberia around 24k ybp.

### A-*Hs* Palaeolithic hunter-gatherers

#### Peștera Muierii 1 (“Women’s cave”, Romania, Eastern Europe)

Peștera Muierii, or the “Women’s Cave”, opens in a complex karst system in the Carpathian Mountains. The cave yielded numerous cave bear remains as well as a *Hs* human skull.

Together with the human remains found in 1952 – a skull and some post-cranial remains (Peștera Muierii 1 or PM1) a temporal bone (Peștera Muierii 2) that cannot be articulated with the skull, and an isolated fibular diaphysis (Peștera Muierii 3) – lithic artifacts attributed to both the Mousterian technology (usually referred to Neandertals) and to the Aurignacian (associated to early modern humans) were retrieved. The *Hs* remains are dated to about 34 ybp (182), thus, they are somewhat younger than those from the nearby cave of Peștera cu Oase, dated to about 40k ybp (183).

The PM1 skull displays modern human features, including a high forehead, a small maxilla and small eyebrow arches. The large cranial vault and intact facial bones indicate a woman with ‘robust features’.

The mitogenome of the skull (182) obtained from two of its teeth suggest that the female from Peștera Muierii was part of the first population of *Hs* after its expansion into Eurasia.

The genome was recently analysed (36) and shows some level of admixture (∼3.1%) and also a quite high level of genetic diversity, a common trait within the earliest European. The human remains of Peștera Muierii represent a group that was a side branch to the ancestor of modern-day Europeans that underwent a great bottleneck during and after the most recent Ice Age.

### Ust’-Ishim (cave on the bank of the Irtysh river, Central Asia, Northern Siberia)

Ust’-Ishim Man is the name given to the fossil of a *Hs* femur, fortuitously discovered in 2008 on the bank of the Irtysh River by a Russian sculptor near Ust’-Ishim, in the Omsk province of north-western Siberia. This fossil is notable for the preservation of its DNA, which allowed the complete sequencing of its genome, thus becoming the earliest – > 45k ybp – modern human genome sequenced to date (33,184).

The examination genome shows that Ust’-Ishim individual is closely related to the 24 ybp Mal’ta boy (see below) from central Siberia, but also to the much later individual from La Braña (Spain) who lived about 8,000 ybp. This genetic relationship show that Ust’-Ishim femur belong to a population that predated the separation between western Eurasians (European population) and the eastern Eurasians (Asians populations) (185). Some authors consider Ust’-Ishim likely belonged to a population that predated this split or represented another migration to the Asian continent that would not have left descendants among the present human populations (151). The same authors observed that Neanderthal DNA in Ust’-Ishim appears in longer sequences, indicating that interbreeding between modern humans and *Hn* took place shortly before the life time of this fossil. Calibration of the genetic clock, following the sequencing of the Ust’-Ishim genome, permits to estimate the date of interbreeding between *Hs* and *Hn* at between 52,000 and 58,000 ybp. This age corresponds to the estimated age of the last exit of *Hs* from Africa (148). No hybridisation between Denisovans and Ust’-Ishim has been demonstrated, although Denisovans are thought to have lived at the same time in East Asia as suggested by the DNA retrieved in the sediments of Denisova (146,151).

### Mal’ta (open air site, Eastern Siberia, Central Asia, Russia)

The four-year-old **Mal’ta** boy lived along the Bolshaya Belaya River near Irkutsk, in central Siberia. This individual is dated 24,300 ybp and belonged to a population settled between the Lake Baikal and the Yanisey River under very rigid climatic conditions with few plants and animals to live on. Nonetheless, they survived in a large community – much larger than the Neandertal’s ones – and built hypogeum construction made of mammoths’ bones and reindeer antler likely covered with leather, and they were capable of fine carving amulets using the ivory and the antler of that animals (186). The Mal’ta eastern Paleo-Siberian group became genetically isolated from its original population being different from the other populations of the eastern Asia (187). The Mal’ta population is ancestral to Native Americans that received around the 40% of their ancestry (37) and to the present-day Kets and the Selkups, known as highly carnivores (188). Our data clearly show that this same adaptation is evident in Neandertal – Spy I/94a and D5 Altai Neandertal – and also Denisovan – D3. Mal’ta represents the closest link to Motala 12, since the Swedish late hunter-gatherer shows about 22% of ancient North Eurasian genetic capital (157).

### B. *Hs* from Europe prior the Neolithic: Mesolithic (Motala 12, Loschbour) and early farmer (Stuttgart)

#### Motala 12 (necropolis – Kanaljorden, Sweden, Northern Europe)

Motala 12 is a maxilla belonging to a male individual from the Scandinavian site of Kanaljorden, excavated in the town of Motala between 2009-2013. In this site, attributed to late Mesolithic hunter-gatherers, several individuals - an infant and 10 adult burials – were retrieved on the shore of a small lake. Motala 12 maxilla was directly dated to 7,212 ± 109 BP (SOM p. 4 in (38)). The sequenced genome (38,39) informs that this fossil shares with its cohort large teeth and thick dark hairs and pigmented skin, as it occurred in present days East-Asians (157) and Native Americans. It must be noted that our results finely match with previously available ones for the late northern hunter-gatherers regarding the efficient digestion of dietary carbohydrates and the perception of bitter taste (37).

### Loschbour LB 1-3 (rock shelter, Luxembourg, Central Europe)

The “Loschbour man” (also Loschbur man) is a skeleton discovered in 1935 in rock shelter in <u>Mullerthal</u> (Luxembourg) by Nicolas Thill (SOM p. 2 in (38)). The skeleton now belongs to the collections of the <u>National</u> <u>Museum of Natural History</u> in <u>Luxembourg City</u>. Loschbour man, directly dated in 1998 (AMS) to 7,205 ± 50 ybp (SOM p. 2 in (38)), was a late Mesolithic <u>hunter-gatherer</u> aged 34-47 years old, circa 1.6 m tall who weighed between 58 and 62 kg (189). The Loschbourg population was supplanted by new populations more likely used to herd rather than hunting. According to DNA analysis made from a tooth (LB1-3) by (38), Loschbour man was “a light skinned (white) individual” (90%), with brown or black hair (98%), and likely blue eyes (56%). Loschbour’s late hunter-gatherer was effective in digesting starchy food, displaying multiple copies of *AMY1A/B/C*, as the later agro-pastoralist populations (19) that lived in the same area. Also, this individual shows the PAV haplotype for taste receptor *TAS2R38* as Motala 12 individual. Loschbour shares with ancient North Siberians the duplication of those genes involved in the amino acid catabolic process, with some genes also involved in the urea cycle and in tyrosine and phenylalanine catabolism (188).

### Stuttgart (LBK 1) (Viesenhäuser Hof, Bavaria, Germany)

A tooth – M_2_ listed as LBK1– was genotyped within one of the largest studies dedicated to the sequencing of the Holocene ancestral population for present-day Europeans (38). This is an interesting site where both local population and migrant from Anatolia was buried. A female – 25-to 35 years old – affected by primary hyperparathyroidism, lived all her life in the surrounding as the strontium isotope analysis suggests (190) and was buried in this Early Neolithic site as showed by the associated Linear Band Keramik artifacts dating back to 5100 to 4800 ybp (191).

## Supplementary Text 2

### Methods

#### Detailed steps of the unsupervised procedure identifying genes with differential copy number involved in digestion

The following seven steps of the pipeline will be described in detail:

1. Selection of all available ancient genomes suitable for estimating haploid gene copy number.
2. Mapping of the selected ancient genomes on the modern human reference genome GRCh38/hg38.
3. Clustering of recent paralogs and estimation of haploid gene copy numbers for the genomes identified in 1.
4. Identification of genes with population-dependent variation in copy number.
5. Selection of a subset of genes based on tissue-specific expression and functional relevance.
6. Comparison with CNV data of modern human populations.
7. Gene Ontology (GO) and literature-based analysis for functional characterisation.

### Step 1. Genome selection for genome-wide analysis

From publicly available human ancient genomes, we identified those that met specific criteria necessary for accurate CNV analysis (see **Table S1**). Specifically, we included genomes that:

1. had a minimum of 1x coverage of the human reference sequence GRCh37/hg19, with available alignment;
2. exhibited less than 5% by contamination from modern human sequences;
3. retained sequences aligned with repetitive regions (indicated by a MAPQ value of 0).

The three conditions allowed 15 archaic human genomes to be considered. It is worth mentioning that some very interesting genomes, although characterized by a sufficiently high coverage, were discarded for various reasons: the available alignment files for Anzick-1, Kostenki 14, Sunghir 1--4, and Sunghir 6 do not comprise repeated regions; LaBraña lacks data for chromosome 22; Den D2, Hohlenstein Stadel, Scladina, San Teodoro 3, San Teodoro 5, and Zlatý kůň have low coverage and/or high contamination.

Reads from these 15 genomes were remapped to the GRCh38/hg38 (42), and CNV estimation was directly based on hg38. Starting with 19,988 human genes, we organized them into 18,043 genes/gene clusters (see below), which are covered by the 15 genomes and shared with the modern reference human genomes hg38.

### Data download and general settings

Reads for the ancient genomes were downloaded either from ENA database or cdna.eva.mpg.de or NCBI with accession IDs in **Table S1**. For genomes available in the ENA browser, all reads in fastq files have been mapped into the GRCh38/hg38 refrence genome. Vindija 33.19 genome is an exception since it displays more than 30B reads and we decided to unmap the BAM files for hg19 available at cdna.eva.mpg.de, and re-map approximately 1.7B reads on hg38. For the genome of Chagyrskaya 8, not available on ENA, we unmapped the BAM files for hg19 available at cdna.eva.mpg.de, and re-mapped approximately 1.2B reads on hg38. All the 79M reads of the Mal’ta genome which were available on NCBI, have been mapped on hg38. Each genome was processed independently up to the final CNV analysis (see below), with the following settings:

- genome = “GenomeID”
- fr_ln = “Fragment length used for sequencing (might differ from genome to genome)”
- autosome_sum_length = “Sum of the lengths of all autosomes in the reference genome (GRC37/hg38)”

### Scripts for the different steps of CNV analysis

We developed four main scripts for data preparation towards CNV analysis:

- **read2CNV.slurm**: handles *de novo* mapping, data preparation, and CNV estimation for individual genomes (Figure 2AB);
- **CNV_Analyses.slurm**: merges single genome estimations and clusters recent paralogs (Figure 2C);
- **genes_with_CNV_at_pop_lvl_extracter.R**: selects genes with differential copy number at the population level (Figure 2C);
- **HPA_Info_Extracter.R**: performs gene expression analysis using data from the Human Protein Atlas (Figure 2D).

All command lines and programs will be available with the publication. Note that read2CNV.slurm is designed for processing single genomes, while CNV_Analyses.slurm is designed to combine the results from individual genomes.

### Step 2. *De novo* mapping, data preparation and CNV estimates

This step is performed using the slurm file **read2CNV.slurm**. It takes as input the reads of a single ancient genome, maps them to the GRCh38/hg38 reference genome and, following the filtering process detailed below, returns the copy number estimate for each gene.

### De novo mapping

The reads in .fastq format from each ancient genome were mapped to the human reference GRC37/hgr38 as follows. The output is then processed with a filter on PCR duplicates, on the percentage of identity, and on read length. All parameters were set in accordance with the guidelines given in (192) and (193), and taken up by (53) specifically for the study of ancient genomes.

### a - Alignment to the reference genome

Reads were mapped to the reference through bwa aln (v. 0.7.17-r1188) (194) with parameters -l 16500, -n 0.01, -o 2, and -t 48. Note that it has been shown that bwa mem works worse when mapping reads from ancient genomes (193). Indexed files from hg38 were retrieved from Zenodo database (https://zenodo.org/records/5146236/files/bwa.tar.gz?download=1) and were stored in a single directory named Data/References/hg38/bwa_2021/hg38, while fasta sequences were obtained from https://hgdownload2.soe.ucsc.edu/goldenPath/hg38/bigZips/hg38.fa.gz.

### b - Format conversions

Mapped reads were converted to SAM format through bwa samse with default parameters. Aligned reads were then filtered to exclude unmapped reads and converted to BAM format using Samtools, samtools view -F 4 -h -Su (v. 1.6) (195,196). The resulting BAM file was sorted using Samtools sort and saved to Analyses/mapping/$genome/${genome}.bam.

### Filter out of PCR duplicates

To remove PCR duplicates, the BAM file was processed using Picard’s MarkDuplicates tool with REMOVE_DUPLICATES and REMOVE_SEQUENCING_DUPLICATES set to *true* (http://broadinstitute.github.io/picard/). This process generated a new BAM file without PCR duplicates and a metrics file recording the details of the duplicates removed.

### Filter reads with percentage of identity < than 90% and shorter than 30bp

In order to remove reads with a percentage of identity lower than 90% and shorter than 30bp, the BAM file was first converted to BED through bedtools bamtobed (v2.30.0) (197). Read lengths and percentage of mismatches were then retrieved through custom bash scripts. Finally, reads were filtered out with Picard’s FilterSamReads and the output was saved in Analyses/mapping/$genome/${genome}_nopcr_90perc_30bp.bam. The 90% threshold was chosen based on standard practices in read mapping for both ancient and modern genomes, while the 30bp threshold was selected to capture a larger number of reads. The validity of these parameters was confirmed using the Mezmaskaia 1 genome.

### Data preparation

D*ata preparation* is carried out in several steps, creating distinct directories and subdirectories:

1. The BAM file is separated in 24 separate files, one for each chromosome, with numbers from 1 to 22 or letter X/Y. From now on, all the analyses of this section will be performed on each chromosome.
2. The GC-bias for each chromosome is computed and each alignment is corrected based on the results.
3. The per-base depth of coverage for each chromosome is computed.
4. The average depth of coverage for each chromosome is computed and the results are concatenated in a single file.
5. The average depth of coverage is calculated excluding the blacklisted regions from step 3.
6. A .txt file containing the average depth of coverage of the genome, taking into account all autosomes, is generated.
7. A gzipped file containing the depth of coverage of each base of protein coding genes defined in the .bed reference is obtained. This will be the main input for the CNV estimation.

Next-generation sequencing (NGS) technologies can introduce, to varying degrees, uneven coverage of reads resulting from GC bias (198). As our analysis of copy number variation is based on aligned reads and depth of coverage of specific regions, to correct the datasets for such biases is of utmost importance. We therefore used deepTools2 (v. 3.5.4) (199) to compute and correct GC-bias using Benjamini’s method (200). More precisely, we used the utility computeGCBias with parameters –effectiveGenomeSize chr_size and -bl blacklist.bed to compute the GC-bias, where chr_size is the ungapped length of placed scaffolds of the chromosome under consideration in GRCh38/hg38 reference genome (lengths are available in column “Placed scaffolds” of section “Ungapped lengths” at https://www.ncbi.nlm.nih.gov/grc/human/data) and blacklist.bed refers to the ENCODE DAC Blacklisted Regions (201,202) (i.e. a comprehensive set of regions in the human genome that have anomalous, unstructured, high signal/read counts in NGS experiments). Finally, alignment files were corrected for GC-bias using the utility correctGCBias with parameter --effectiveGenomeSize chr_size. Given a genome G, the depth of coverage (average or per-base) was computed using mosdepth (v. 0.3.3) (203) with default parameters on the GC-bias corrected alignment files. More precisely, given a chromosome chri where *i*∈*{1,…,22}*, its average depth of coverage $C^i$ was computed as

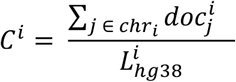

where 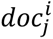 is the depth of coverage computed by mosdepth at position *j* of chromosome *i,* and 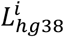 is the ungapped length of chromosome *i* in the GRCh38/hg38 reference genome. As analyzed datasets involve male and female individuals, we decided to restrict the copy number variation analysis to autosomes.

A set of protein coding genes G for GRCh38/hg38 was retrieved from GENCODE (204) and genes overlapping any of the ENCODE DAC Blacklisted Regions (201,202) were excluded from the analysis, regardless of the length of the overlap.

The coverage 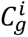 of a gene *g* annotated on chromosome *i* is computed as

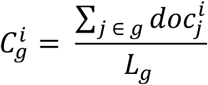

where the depth of coverage 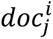 is computed exclusively at positions *j* of the gene *g* of length *L_g_*. Hence, we estimate the *haploid* copy number of a gene *g* as 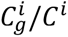.

### Step 3. CNV estimation

For each genome, CNVs were estimated with a custom python script (computeGeneCopy.py) which is called in the main slurm file. The script uses two intermediate input files:

- chrom_doc=“Analyses/Data_preparation/$genome/avgdepth/nobl/$genome.hg38.nobl.avgdepth.t xt”, which provides the average depth of coverage, calculated by excluding blacklisted regions, with each line corresponding to a chromosome.
- gene_doc=“Analyses/Data_preparation/$genome/$genome.fixed.hg38.genes.protein_coding.cov erage.gz”, which contains the depth of coverage for each base of protein coding genes.

The output, Analyses/CNV_estimates/$CNV_estimates_dir/$genome.nobl.cnv.tsv, contains the CNV estimates for each gene in the genome.

### Merging single genome estimates and clustering of recent paralogs

This step is performed using the slurm file **CNV_Analyses.slurm**, which takes as input the gene CNV estimates from each ancient genome ($genome.nobl.cnv.tsv). The process generates two output tables:

- The first table, AllGenomes.type.nobl.cnv.matrix.ALL.tsv, groups genes into clusters of recent paralogs, with each cluster represented as a single line, replacing individual paralog entries. Each column corresponds to an ancient genome, and an additional column provides CNV estimates for genes/clusters of genes in the human reference genome hg38. A total of 18,043 genes/clusters of genes are included.
- The second table, AllGenomes.type.nobl.cnv.matrix.tsv, selects only those genes/clusters of genes that show a different number of copies in at least one genome when compared to hg38. This results in a subset of 2,217 genes/clusters of genes.

The CNVs estimates for each genome were unified in a single file through bash commands. Then, human paralog genes, defined as duplicated copies at the *Homo sapiens* level in ENSEMBL database (column “Ancestral Taxonomy” in ENSEMBL), were clustered together if they had over 80% sequence identity. For exemple, the three copies of the AMY1 gene appeared at the *Homo sapiens* level, remain highly conserved and are merged into a cluster AMY1A/B/C. After clustering, the estimated copy numbers of the “clustered genes” is defined by adding together the copy number estimation of each of the paralogs. Note that, retrieval of human paralogs from the ENSEMBL database was done with a custom R scripts, using the package BiomaRt (205,206).

### Definition of population-dependent CNV

This step is performed using the R script **genes_with_CNV_at_pop_lvl_extracter.R**. It takes the input file AllGenomes.type.nobl.cnv.matrix.tsv and generates the output file AllGenomes.type.nobl.cnv.matrix_ with_CNV_pop_lvl.tsv, which includes two additional columns, “gain” and “loss”, reflecting differences in CNV between *Hn* and/or *Hs* populations. Genes without such differences are discarded, resulting in a final table of 781 genes.

To define population-dependent CNVs, we focused on genes showing an increase or decrease in copy number in at least 5 Neandertals (>50% of the population, where pop=8) and/or in at least 4 ancient *Sapiens* (>50% of the population, where pop=6), compared to the reference human genome hg38. Since only one *Denisova* specimen (Den *3*) was available, it was treated as a point of comparison rather than a population.

### Step 5. Analysis of tissue-specific expression and functional relevance

This step is performed using the R script **HPA_Info_Extracter.R**. It uses two downloadable datasets from the Human Protein Atlas (HPA) (44,207–209) as input: https://www.proteinatlas.org/download/rna_tissue_consensus.tsv.zip for RNA levels, and https://www.proteinatlas.org/download/normal_tissue.tsv.zip for protein expression. Two output files are generated. The first file (all_genes_3organs_protein.tsv) lists the three organs with the highest protein expression for each human gene. The second file (all_genes_3organs_rna.tsv) lists the three organs with the highest RNA expression for each human gene.

All human genes were analyzed using data from HPA. For each gene, the three organs with the highest RNA levels (given as a continous value) and the three organs with the highest protein expression levels (three values are given, low, medium and high) were identified. In cases where multiple organs showed equal expression levels, additional organs were included. The subset of 781 genes with differential CNV at the population level (see above) was combined with the RNA/protein expression analysis. For each gene, we considered the list of organs with highest RNA/protein expression and selected only those associated to at least one digestive organ—the gastrointestinal tract, pancreas, proximal digestive tract, liver and gallbladder. These 489 selected genes were classified as “digestion-related genes”.

### Filtering for strictly diet-related and accessory diet-related genes using HPA

Each gene was manually reviewed using the HPA database to assess whether it was expressed in a wide range of organs beyond digestive ones. Additionally, “protein function” data from UniProt, accessed through HPA, or “gene description” from Ensembl (when UniProt data was unavailable in HPA) was used to exclude genes with overly generic functions associated with multiple organs (Figure 2E). This manual screening resulted in a final set of 55 genes/clusters of genes.

### Step 6. Filtering by comparison with modern human CNVs

CNV estimates for the subset of 55 genes in ancient genomes were compared with those from modern populations (Figure 2F). This step aimed to ensure that the analysis captured adaptive variations specific to the two ancient populations, excluding genes with common variation in modern populations. An example is provided in the main text for the *PGA3/4/5* gene, and **Figure S3** shows all genes filtered out by this process. This step refined the final set to 50 genes.

### Step 7. GO-terms analysis and diet-related literature

All genes chosen based on the Tissue expression and passing the modern CNV estimation filter have been analysed with a GO-term analysis. The 50 selected genes were given as an input to pantherdb.org (210) to compute a first functional classification. Subsequently, the same has been done with DAVID online server (45,211) and the resulting GO-terms were grouped in slim categories with QuickGO online server (212). Subsequentely, each gene/cluster of genes was analysed further based on literature search to decide how to classify them in six functional categories: dietary carbohydrate metabolism, lipid metabolism, liver lipid/brown fat metabolism, immunity/antibiotic action in the digestive tract, environmental related pathways, obesity/diabetes. See Figure 2GH.

### Estimation of the “absence” of a gene

We estimated that some genes (*UGT2B28, ACOT1, CD24, LYPD8, PECAM1, TCAF2, TCAF2C*, and *TMEM86B*) are not present in some ancient population or individual. For instance, the *ACOT1* gene is not present in *Hn,* Den D3 and Loschbour genomes. In this respect, notice that in the 8 *Hn* genomes, *ACOT1* coverage is estimated to be at most <0.42 (**Table S5**), which can be explained by the presence of the paralogous gene *ACOT2*, duplicated at the Homininae level, and of many other copies which preceeded the *ACOT2* duplication, such as *ACOT4* with 70% sequence identity from *ACOT1. ACOT1* and *ACOT2* are known to have the same function, *ACOT1* acting in the cytosol and *ACOT2* in mitochondria (213). *ACOT2* codes for a protein of 483 amino acids that contains a 62 amino acid leader sequence at the N-terminal that targets the protein to mitochondria. The *ACOT1* protein is shorter than *ACOT2*, with its 421 amino acids showing 98.81% sequence identity to *ACOT2*, differing by only 5 amino acids. Read mapping to *ACOT2* and *ACOT4* is frequently estimated below 1, which supports the absence of the *ACOT1* gene copy in the genome and suggests incorrect read mapping to *ACOT1*. See (214) for a general description of type 1 and 2 *ACOT* genes. For the other genes: *UGT2B28* has a paralog *UGT2B11* of 94.71% sequence identity, duplicated at the Hominoidea level; *TCAF2* has a paralog *TCAF2C* at 99.58% sequence identity, duplicated at the Apes and Old World Monkey/Catarrhini level. *CD24, LYPD8, PECAM1* and *TMEM86A* do not have recent paralogs within *Homo sapiens* nor sufficiently high sequence identity, leaving their absence unexplained. For example, *TMEM86B* and *TMEM86A* originated from a duplication event at the *Vertebrata* level and exhibit 36.28% and 34.17% sequence identity, respectively, when *TMEM86B* is used as the query and *TMEM86A* as the target. This low sequence identity between these paralogs in the reference human genome is insufficient to support the possibility of incorrect read mapping.

### Construction of the multidimensional space defined by CNV

#### Individuals plotted in the CNV multidimensional space

We represented individuals in a multidimensional space by using gene CNVs as real-valued features.

#### Principal Components Analysis (PCA)

PCA is a well-known computational method used in exploratory data analysis and for making predictive models (50). It is commonly used for dimensionality reduction when the number of features (or dimensions) in a given dataset is too high. We used PCA to project each data point onto only the first few principal components to obtain lower-dimensional data while preserving as much of the variation as possible. The first principal component can be defined as a direction that maximizes the variance of the projected data. The 2nd principal component can be taken as a direction orthogonal to the first principal component that maximizes the variance of the projected data, and so on. Thus, a new dimensional system, where each dimension relies on a combination of the initial dimensions of the data, is generated.

CNVs of the final set of genes were studied from a statistical point of view in two subsets: strictly diet-related (23 genes) and accessories to diet (35 genes, different from the 23 genes above). In both cases, a PCA has been performed to reduce the 23 (or 35) dimensions of the high-dimensional space where *Hn* and *Hs* individuals are represented by their gene CNVs. In Figure 4, the position of the reference genome GRGh38/hg38 was plotted for comparison. PCA was performed with the program R (215) with prcomp function and scale set to *TRUE*. The contribution of each variable to the first three components has been computed through the function *fviz_contrib* from the Factoextra package (216), and a 2D plot of the two most representative PCs was otained using the plotly package (v4.9.3) (217).

The dotted red line is the expected average contribution. A variable with a higher contribution than this is considered important, and was highlighted in Figure 4CD. For the visualization, we used the function *plot_ly* of the R package plotly. With this function the main dimensions of the PCA to be represented cin the n-dimensional space can be chosen. We chose to represent the first three principal components.

### The fossils, geographical regions and time period

The fossils/genomes (**Table S2**) of the 15 individuals have been associated to their geographical region and time period (**Figure S5** and **Figure S6**). For this, we defined six different regions of the Asian European belt, from the most eastern to the most western: Central Asia, Eastern Europe, Central Europe, Northern Europe, Southern Europe, Western Europe. These geographic areas were identified according to (218,219), modelling eco-geographical and morphological data on the Neandertalian territory and variability of mtDNA. To take ecological diversity into account, we added Central Asia and Central Europe to the geographic area already defined in these two publications. Indeed, since the latter studies, new evidence for the great *Hn* extension in Central Asia has been collected (from Altai Neandertal and Chagyrskaya caves). Also, we reduced the “Northern Europe” area by splitting it into two regions and adding an eastern one, named “Central Europe”, where the climate was different. This geographical division is used here for *Hn* and *Hs*. We considered seven time periods: 120k-90k ybp, 90k-50k ybp, 50k-45k ybp, 45k-40k ybp, 40k-35k ybp, 30k-20k ybp, and 10k-5k ybp.

## Supplementary Text 3

### A description of the 19 strictly diet-related genes

In the following section, refer to **Table 1** for the list of diet-related genes identified in our study, and to Figure 3 for a direct CNV comparison between ancient individuals and those from modern human populations.

### Duplication of three gene clusters and one gene adapted to carbohydrate digestion

***AMY1*** is highly duplicated on the human chromosome 1 and we refer to it as a gene cluster, ***AMY1A/B/C***. *AMY1A/B/C* is expressed in salivary glands (Figure 2E) and protein expression of *AMY1C* has been also observed in the pancreas (together with the pancreatic amylase gene *AMY2*). Amylases **efficiently break down starch into maltose** in the oral cavity (7,9). The CNV of *AMY1* has an observed range of 2–18 copies per person (5,8–10) and an average of 6 copies per person (25). *AMY1A/B/C* duplication has been estimated to have emerged after the divergence between human-specific lineages and archaic hominins either approximately 550-590k ybp (5,9) or around 400-275k ybp (9,10,19,107), a reconstruction that agrees with recent analyses of genomes from Cameroon (220). Across the 15 ancient genomes studied here, our data confirm that *AMY1* is highly duplicated in *Hs,* whereas two copies are present in most *Hn* and *Den* (**Table 1AB** and Figure 3). More precisely, the trend in *Hs* since Ust’Ishim dated at 45k ybp, including present-day humans, is of copy numbers greater than 3. This particularly high copy number is additional evidence that *AMY1A/B/C* was duplicated after the ancestral lineages split. Interestingly, the late Palaeolithic hunter-gatherer from Motala 12 (Sweden) has two copies of the cluster, although Mal’ta, the easternmost Upper Palaeolithic modern human who lived 24k ybp in severe climatic conditions in Central Asia, has only two copies of the *AMY1* gene. On the other hand, a late *Hn* in our sample, Les Cottés ZA-1514 has 3 copies.

***SULT1A3*** and ***SULT1A4***, collectively called *SULT1A3/4*, encode identical protein products (221). Previous genomic studies revealed the duplication of these genes during the evolutionary process (222,223). From our analysis, all the EAHs have only one copy of *SULT1A3/4* whereas all the *Hs*, from the oldest to present-day humans, have 2 or 3 copies of this gene cluster. *SULT1A3/4* is a **sulfotransferase enzyme** catalyzing the sulfate conjugation of many hormones, neurotransmitters, drugs, and xenobiotic compounds, and so its function depends on its tissue location (224). Highly expressed in the gastrointestinal tract of humans and closely-related primates, *SULT1A3/4* indirectly **influences carbohydrate metabolism** (225) through dopamine sulfonation, a reaction involved in the regulation and biotransformation of catecholamines (226). The gene cluster may therefore detoxify potentially lethal dietary monoamines in the intestine and eliminate toxic compounds from the body (227). A diet that includes a large intake of starch, nuts and seeds, is rich in flavonoids, which boost the levels of serotonin and dopamine, two key neurotransmitters regulating many physiological processes under hormonal control.

***SLX1A/B*** encodes proteins that play a crucial role in **maintaining genome stability**, DNA repair, homologous recombination, and telomere maintenance (228). These proteins form the catalytic subunit of the *SLX1-SLX4* structure-specific endonuclease complex, which is essential for resolving DNA secondary structures that arise during repair and recombination processes. They exhibit read-through transcription with the adjacent carbohydrates-related *SULT1A3* gene. Note that the CNV pattern observed in *SLX1A/B* is the same as for *SULT1A3/4*, possibly suggesting a co-evolution between gene clusters.

***UGT2B28*** encodes a protein that belongs to the uridine diphosphoglucuronosyltransferase family. This enzyme facilitates the **transfer of glucuronic acid** from uridine diphosphoglucuronic acid to various substrates, including steroid hormones and lipid-soluble drugs, through the process of glucuronidation, an intermediate step in steroid metabolism. Dietary intake can influence this metabolic pathway, as certain foods, particularly those **rich in flavonoids**, can affect glucuronidation activity. Consuming a diet high in fruits, vegetables, and other flavonoid-rich foods may enhance the body’s ability to metabolize and eliminate various substances, potentially impacting the activity of *UGT2B28*. **Flavonoids have been found to interfere with the digestion, absorption, and metabolism of carbohydrates** (229). Also, significant amounts of flavonoids can be found in leafy vegetables, onions, apples, berries, cherries, soybeans, and citrus fruits (230), tea and wine.

### Gene copy number for adaptation to fatty food of animal origin

A large number of genes directly or indirectly related to high intake of **animal fat** present a differentiated CNV in EAHs (**Table 1A**). A few genes are differentiated either in *Hn* (*LPA, ACOT1, ZNG1E/F/C*) or in *Hs* (*ACOX3, HAND1*), but the vast majority have a higher CNV for both *Hn* and *Hs* than the modern reference genome.

***LPA, CLPS*** and ***CLPSL1*** all **encode enzymes involved in lipid hydrolysis and oxidation of long-chain FA** (i.e., arachidonic acid metabolism). More FA are consumed in diets incorporating more animal food and are critical for brain development and metabolism and for inflammatory responses (91). *CLPS*, *CLPSL1* and *LPA* genes are **involved in lipid digestion and lipid transport** (231). Colipases (*CLPS* and its paralog *CLPSL1*, Colipase Like 1) are cofactors needed by pancreatic lipase for efficient dietary lipid hydrolysis as they allow the lipase to anchor itself to the lipid-water interface and avoid being washed off by bile salts (232). Lipases (*LPA*) are mainly expressed in the liver and perform essential roles in digestion, transport and processing of dietary lipids. There is huge inter-individual variation in *LPA* expression levels and a high heritability of the traits, and while the physiological function of *LPA* is not fully elucidated, it is acknowledged to be linked to the risk of cardiovascular disease (233). All *Hn*, almost all *Hs* and *Den* individuals display a positive CNV trend for *CLPS* and *CLPSL1*. Most *Hn* individuals and the early *Hs*, Peştera Muierii 1 and Malta, have multiple copies of *LPA*. The oldest *Hs*, Ust’Ishim, along with later *Hs*, Loschbour, Motala 12 and Stuttgart, exhibit the same pattern as the modern reference genome—a single copy. Notably, most modern individuals in our study show duplication of *LPA*.

***ACOT*** genes are involved in **lipid metabolism** as they encode acyl-CoA thioesterase that regulates the cellular balance between free FA and acyl-CoA, feeding into key pathways for energy expenditure and neuronal function (49). In contrast with *Hs*, none of the *Hn* individuals has the *ACOT1* gene whereas copies of all other type I *ACOT* genes (214), whether peroxisomal or mitochondrial, are present in *Hn*, and most *Hs*. *ACOT2* is 98.6% identical (213) to *ACOT1* and is present as one copy in all *Hs*, *Den* and *Hn* individuals. The *ACOT2* protein localizes to the mitochondrial matrix and shares the same function as *ACOT1* by hydrolysing long-chain fatty acyl-CoA into free fatty acid (FA) and CoASH. It has been demonstrated that *ACOT1* regulates fasting hepatic FA metabolism by balancing oxidative flux and capacity (84). Hence the absence of *ACOT1* could have been lethal for EAHs in periods when FA intake was scarce, such as during the colder stages of marine isotope stage (MIS) 3 (chiefly Heinrich Event 4) when a strong reduction in herds of large herbivores deprived EAHs of their basic food (234,235). Conversely, *Hs* would have had broader access to different kinds of food. An exception is the individual from Loschbour who lacked this gene.

***RFPL4A/L1*** produces a protein **metal-binding zinc**, an essential mineral, and the ***ZNG1E/F/C*** cluster, crucial for **zinc transfers**, exhibits a notable trend of increased CNV in *Hn* and certain *Hs*. Notably, the richest sources of zinc are animal products and oysters (71) compared to plant foods. However, pinenuts, seeds such as hemp, sesame, or chickpeas and lentils among legumes are also rich in zinc and were available in the ecological realm of both *Hn* and *Hs*.

***ACOX3*** (Acyl-Coenzyme A oxidase 3, also known as pristanoyl-CoA oxidase) possibly facilitated **strong intake of animal fat**. Namely, the expression of *ACOX3* has been documented to increase in response to a high-fat diet (236) and to be related to fat metabolism (237). Both *Hs* and a large number of *Hn* were equipped with multiple copies of this gene.

***HAND1***, especially duplicated in *Hs* and in the latest *Hn* (Mezmaiskaya 2, Les Cottés et Goyet), has been shown to inhibit the conversion of cholesterol to steroids in trophoblasts, playing a crucial role in the development and physiological function of the human placenta (96). Further analysis of this gene could provide valuable insights into the relationship between **diet and reproduction** in the evolution of the Homo genus.

***PDX1*** is a transcription factor active in the **pancreas,** where the protein produced by this gene plays as transcriptional activator of several genes, including insulin, somatostatin, glucokinase, islet amyloid polypeptide, and glucose transporter type 2. It is essential for pancreatic β-cell development and adult β-cell function (238). It has been documented to **mediate lipid metabolism** gene networks in intestinal cells (239). In high-fat diets, **saturated fatty acids** entrap *PDX1* in stress granules and impede islet beta cell function (240). An increased production of *PDX1* due to the higher number of copies of the gene could facilitate the activity of *PDX1* in enhancing the efficiency of this enzyme activity in *Den*, *Hn* and *Hs*.

***WFS1*** encodes for a protein called wolframin that is thought to **regulate the amount of calcium** in cells. It is expressed in all the endoplasmic reticulum of all the cells but shows its highest levels in brain, pancreas, heart, and insulinoma beta-cell lines tissues. Due to the varied and fundamental functions played by calcium in cellular regulation, it plays a fundamental role in cell-to-cell communication, muscles contraction, triggering the myosin and actin filaments to slide into each other, and protein processing. *WFS1*’s association with diet has been elucidated in (72), where it was demonstrated that heterozygous *WFS1* mice, possessing only one functional gene copy, exhibited metabolic impairments when subjected to a high-fat diet. The presence of multiple copies could facilitate individuals following high-fat regimes such as *Hn*, *Hs*, and *Den*.

***H2AC11*** is a member of the histone deacetylase family and it is involved in the biological functions of almost every system of the human body and its duplication may play a role in tumorigenesis (241). **Deficiency in *H2AC11* prevents the onset of obesity and metabolic syndrome induced by a high-fat diet** (242). Surprisingly, we observe a duplication in all *Hs* and *Den*, while we see a heterogeneous pattern with regard to *Hn*, suggesting that the role of this gene in diet needs to be further investigated.

***F7*** (coagulation factor *F7*) encodes for an enzyme of the serine protease class. It is part of a group of proteins produced in the liver and it is vitamin K-dependent. They are involved in the **coagulation system**, a series of chemical reactions that form blood clots. *F7* is the most common of the hemophilia-like bleeding disorders. **Quick time for coagulation**, induced by the duplication of the gene, may have provided a leap to both *Hn* and *Hs*, who might have inherited these traits while breeding with *Hn* (75).

Under possible harming conditions that *Hn*, *Den* and *Hs* might have encountered, the capacity of quickly healing the wound so that being less exposed to infection, ending with a higher probability to survive and thrive. Yet, under far less dangerous settings, hypercoagulation might have turned into a higher risk for stroke, embolism and troublesome pregnancies (243). *F7* has also been documented to be **linked to fat intake and cholesterol levels** in mice (98). The impact of diet on *F7* activity is reported in (99).

### Adaptation to cold stress metabolism related to eating food of animal origin

Cold conditions are expected to be better tolerated by human populations that react better to cold stress. We found that both *Hn* and *Hs* populations show a positive CNV trend for a gene involved in this metabolic process – ***MST1*** - compared to the modern human genome. We noted that the trend was due to the positive differential CNV of this gene in the Holocene *Hs* individuals, while the three early *Hs*, Ust’Ishim, Peştera Muierii 1 and Mal’ta, belonging to the earliest migration wave out of Africa about 60ka, have single copies of each gene. One more gene shows higher CNV in *Hs*, ***H1.2***, and one lower, ***CD24***.

***MST1*** (Macrophage stimulating 1) gene is known to be important in **liver lipid metabolism**, limiting liver damage that might be induced by a high-fat diet or during fasting (244). This gene also plays a role in **cellular responses to hypoxia**, **oxygen supply** and **thermogenic metabolism**. Other genes, *EPAS1* (endothelial PAS domain containing protein 1) and *EGLN1* (Egl-9 family hypoxia inducible factor 2) have previously been highlighted as being involved in the physiological response to oxygen depletion in ancient populations (69). Indeed, tolerance of lower oxygen availability was observed in modern Andean and Tibetan people who have hemoglobin with adapted oxygen affinities (245,246). When the *EPAS1* gene was identified in *Den* DNA it was proposed that the capacity of Tibetan people to thrive in low-oxygen or high-altitude conditions was possibly inherited through introgression of Den-like DNA (69). *MST1* is duplicated in *Den*, all *Hn* and Mesolithic *Hs* individuals.

***H1.2*, highly expressed in adipocytes,** regulates thermogenic genes in white adipose tissue (WAT) and affects energy expenditure. Its deletion promotes browning of WAT (brown adipose tissue - BAT) and **improves cold tolerance**, while its overexpression has the opposite effect (66). In cold environments, it has been observed a higher expression of *H1.2*, sensing thermogenic stimuli in WAT and BAT (66). Notably, we observe an increase in the copy number of this gene in the most part of *Hs* while it remains at a single copy in half of *Hn*, indicating a plausible adaptation to different environments.

***CD24*** gene loss, observed in the older *Hn* (Altai Neandertal D5, Chagyrskaia 8) and Vindija 33.19, and most *Hs*, has been linked to a **reduction of white adypocyte tissue** (67).

### A description of the 31 accessory to diet genes

High fat diets are known to be linked to several diseases including obesity, diabetes, fatty liver, inflammatory bowel disease and colon cancer (247). The set of 31 accessory genes is linked to these diseases. In the following section, refer to **Table 2** for the list of accessory genes identified in our study, and to **Figure S2** for a direct CNV comparison between ancient individuals and those from modern human populations.

### Inflammation/Immune response

***BOLA2/2B*** is involved in the immune response in the gut and may be associated with an increased risk of digestive disorders such as **inflammatory** bowel disease. It is known to be involved also in the maturation of cytosolic iron-sulfur proteins. Its duplication modifies iron homeostasis and its deletion is associated with anemia as it is active in the iron regulation. It is conjectured that its emergence is unique to *H. sapiens* and dated back to 282 kyr (24) and its expansion in modern populations, has a potential adaptive role in protecting against iron deficiency crucial during embryonic development (97). No *Hn* nor *Den* show duplication of this gene, which is duplicated in all *Hs*, including modern individuals and the reference human genome. Notably, copy number variation is highly diverse within modern human populations.

***ORM1*** encodes a key acute phase plasma protein, classified as an acute-phase reactant due to its increase during acute inflammation, and it may be involved in aspects of **immunosuppression**. ***ORM2*** (ancient paralogue of *ORM1*) encodes an acute-phase plasma protein with a role in **immunosuppression during acute inflammation**. *ORM1* and *ORM2* **regulate lipid homeostasis** and protein quality control (248).

***TPSAB1/B2*** encodes alpha-tryptase, the major neutral protease in mast cells, playing a role in **innate immunity**. Mast cell tryptases have proposed roles in allergic inflammation and host defense against infection (249). Duplication of this gene can cause severe gastrointestinal disorders and stomach disorders (250). Hereditary alpha tryptasemia syndrome (HαT) is an autosomal dominant genetic disorder caused by an increased number of copies of the *TPSAB1* gene. This condition is characterized by elevated basal serum tryptase levels. The syndrome presents with symptoms that can affect multiple organ systems (250). Nearly all *Hn*, *Hs*, and *Den* have a single copy of this gene, in contrast to the modern human reference genome and all modern human populations, which dispay two copies.

***GRTP1*** is associated with poor prognosis in stomach and colon cancer when expressed at low levels (251). Most expressed in liver and intestine, poorly studied and identified as participating in the multigene signature for **non-alcoholic fatty liver disease** (NAFLD) (252). It is duplicated in *Den*, *Hn* (with the exception of Spy 94a), and all *Hs*.

***CLPTM1L*** (Cleft lip and palate transmembrane protein 1-like protein) overexpression can induce apoptosis, and polymorphisms in this gene increase susceptibility to various cancers, including lung, pancreatic, and breast cancers. It participates in the **biosynthesis of glycosylphosphatidylinositol and lipid transport** (253). Most expressed in liver and pancreas. Nearly all *Hn* individuals show duplication of this gene.

***ZNF431*** is a member of the Krueppel C2H2-type zinc-finger family of transcription factors, the largest transcription factor family in the mammalian genome. It may negatively regulate transcription of target genes and mutations may be linked to non-syndromic facial clefting. It has been observed that homozygous mutations in the *ZNF431* gene, inherited in an autosomal recessive manner, result in reduced STAT3 levels, producing a phenotype that closely resembles autosomal dominant hyper IgE syndrome (254). Hyper IgE syndromes (HIES) are rare forms of **primary immunodeficiencies** (PI) characterized by recurrent eczema, skin abscesses, lung infections, eosinophilia (high numbers of eosinophils in the blood), and high serum levels of immunoglobulin E (IgE). *Den*, most *Hn,* and *Hs* show duplications.

***C4A/B*** encodes complement factors essential for the binding of the CR1 receptor with the preformed and nascent immune complexes (255,256), hence with roles in **immune complex solubilization and clearance**. Deficiency is linked to systemic lupus erythematosus. *Den*, most *Hn,* and *Hs* exhibit a high copy number (at least three copies), whereas all modern human populations consistently show two copies.

***N4BP3*** enhances **antiviral innate immune** signaling by promoting polyubiquitination of MAVS. It plays a crucial role in the RIG-I-like receptor (RLR)-mediated innate immune response by targeting MAVS, shedding light on the mechanisms underlying innate antiviral responses (257). Most *Hn* and *Hs* exhibit duplication of this gene.

***LYPD8* prevents invasion of Gram-negative bacteria** in the colon epithelium by binding to bacterial flagella (intestinal homeostasis) (258,259). *Hs* and the older *Hn* are depleated of this gene.

***STAB1*** plays a role in host defense mechanisms against infections and regulates **anti-inflammatory responses**, potentially influencing atherosclerosis (260,261). Most *Hn* and *Hs* exhibit duplication of this gene.

***PECAM1*** (platelet endothelial cell adhesion molecule-1) is a 130-kDa transmembrane glycoprotein expressed on blood and vascular cells. It acts as a negative regulator of **nonalcoholic fatty liver disease** progression, nonalcoholic steatohepatitis (NASH) (262). This study shows that **genetic deficiency** of *PECAM1* potentiates the development and progression of NASH. The NASH disease spectrum ranges from **accumulation of fat** in hepatocytes (steatosis) to the presence of an **inflammatory infiltrate and fibrosis**, and ultimately to progressive fibrosis and cirrhosis. *Hs* and the older *Hn* are depleated of this gene.

***UQCRFS1*** is involved in mitochondrial respiratory chain complex assembly and function, with deficiencies linked to mitochondrial disorders (263). A diet-dependent proteome-wide differences in adipose tissues, highlighted down regulation of oxidative phosphorylation genes such as *UQCRFS1* in response to **dietary cholesterol** (264). Multiple copies may benefit individuals with dietary cholesterol and are observed in *Den*, as well as in nearly all *Hn* and *Hs*.

***IFNL2/3*** are cytokines with **antiviral, antitumor, and immunomodulatory activities** that play a crucial role in antiviral host defense, particularly in epithelial tissues (265). They act as ligands for the heterodimeric receptor composed of *IL10RB* and *IFNLR1*, activating the JAK/STAT signaling pathway to induce IFN-stimulated genes (ISGs) and mediate the antiviral state. Their action is primarily restricted to epithelial cells due to the cell-specific expression of the *IFNLR1* receptor and they significantly contribute to antiviral defense in the intestinal epithelium and up-regulate MHC class I antigen expression. All modern populations dispay duplications of this gene, while all ancient individuals exhibit only one copy.

### Environment and nutrition

***TCAF2*** and ***TCAF2C*** are *TRPM8* channel associated factors. They enable transmembrane transporter binding activity and are involved in the regulation of anion channel activity, cell migration, and protein targeting to membranes, with a localization in cell junctions and the plasma membrane. *TCAF2* and ***TCAF2C*** are ancestrally related within *Homo sapiens*, but their sequence similarity does not meet our strict threshold. *TCAF2* is longer than *TCAF2C*, with the two genes showing ENSEMBL Target_id of 99.58 and a Query_id of 77.15. This suggests that *Hn* likely has only 1 copy of the two genes, although it is unclear which of the two, while *Hs* possesses both copies. Research on the **deletion** of the **cold thermoreceptor** TRPM8, which is related to *TCAF2* and *TCAF2C*, shows **increased heat loss and food intake, leading to reduced body temperature and obesity** in mice (266). It has been established that *TCAF* duplications originated ∼1.7 million years ago but diversified only in *Hs* by recurrent structural mutations. Multiple evidence supported the hypothesis that *TCAF* diversification among hominins was likely due to positive selection, in **response to cold or dietary adaptations** (68). Both *TCAF2* and *TCAF2C* are absent in *Hn* and *Den*, yet present in *Hs*. Interestingly, modern human populations align with ancient individuals regarding *TCAF2*. For *TCAF2C,* it is noteworthy that ethnic groups from South Asia similarly lack this gene, as seen in *Den* and *Hn*.

***TMEM86B*** encodes an enzyme that **degrades lysoplasmalogen**, a group of lipids, impacting cell membrane dynamics. Plasmalogens are found in various **foods** such as scallops, squid, mussels, octopus but also **big game cervids** (267–270). Decrease in plasmalogen is related to Alzheimer and Parkinson diseases. All ancient individuals lack this gene, which is present in all modern populations.

***ATP4B*** is essential for **gastric acid secretion**, encoding the beta subunit of the gastric H+, K+-ATPase proton pump, which is vital for exchanging H+ and K+ ions across the plasma membrane (271). All ancient individuals exhibit a duplication of this gene.

***HGFAC***, a member of the peptidase S1 protein family, is **nutritionally regulated** and plays a critical role in **maintaining carbohydrate and lipid homeostasis** by activating hepatocyte growth factor (HGF) (272). All ancient individuals exhibit a duplication of this gene.

***SLC9A3*** is crucial for pH regulation and signal transduction, encoding an **epithelial Na/H exchanger** implicated in congenital secretory sodium diarrhea, bladder fibrosis, and dysfunction due to systemic homeostasis deregulation (273). Most ancient individuals exhibit a duplication of this gene.

### Obesity/Diabete

***FGFRL1*** has a negative effect on cell proliferation and shows significant methylation level differences that **correlate with obesity** status (274). All ancient individuals dispay duplications of this gene, with the exception of Spy 94a.

***DRD5***, a dopamine receptor gene, has been **linked to obesity**, with studies showing altered dopamine levels and receptor expression in adipocytes of obese individuals (275). All ancient individuals dispay duplications of this gene, with the exception of Goyet Q56.1.

***NKX6-1***, a homeobox gene, is essential for the development and function of pancreatic beta cells, and its inactivation causes beta cell dysfunction and hypoinsulinaemia, affecting insulin synthesis and secretion (276). Many ancient individuals display duplications of this gene; however, the oldest *Hn*, Altai Neandertal and Chagyrskaya 8, and Den D3 do not exhibit duplications.

***HAND2***, a basic helix-loop-helix transcription factor, plays a crucial role in cardiac and limb development and has been identified as an **obesity-linked** adipocyte transcription factor (277). It is duplicated in most *Hs*.

***H4C16*** (H4 clustered Histone 5 and Histone 16) is involved in central **carbon metabolism regulation**, and its acetylation status influences **diet-induced obesity** in mice (278). Most *Hs* exhibit a very high duplication number, with at least three copies.

***IDUA*** encodes an enzyme essential for the lysosomal degradation of glycosaminoglycans, with mutations leading to mucopolysaccharidosis type I, and has been shown to **prevent diet-induced obesity** in mice (279). All ancient individuals display duplications of this gene.

***ATP5ME***, part of mitochondrial ATP synthase, catalyzes ATP synthesis during oxidative phosphorylation and has been linked to attenuating type 1 GDM-induced fetal myocardial injury (280). Note that dysfunction of **mitochondria of adipose tissues** results in detrimental effects **on adipocyte differentiation**, lipid metabolism, insulin sensitivity, oxidative capacity, and thermogenesis, which consequently lead to metabolic diseases (281). All ancient individuals, except *Den*, display duplications of this gene.

***SEMA3B*** is part of a gene family known to **regulate obesity** by influencing adipogenesis (282). Most *Hn* display duplications of this gene.

***IFRD2*** deficiency, along with *IFRD1*, results in severely reduced adipose tissue and resistance to **high-fat diet-induced obesity** in mice (283). Most *Hs* display duplications of this gene.

***IER3*** (***IEX-1***) deficiency induces the browning of white adipose tissue and provides **resistance to diet-induced obesity**, highlighting its role in expanding adipocyte precursor cells (284,285). Most *Hs* display duplications of this gene.

***PLXND1*** determines body fat distribution by regulating the collagen microenvironment in visceral adipose tissue, playing a significant role in **obesity and fat distribution** (286). Most *Hs* display duplications of this gene.

## Supplementary Tables

**Table S1.** Download information for the 15 ancient human genomes and the reference genome analyzed in this study. From top to bottom, genomes are listed according to dating from the oldest to the most recent within each population: Denisovan (grey), Neandertal (orange), *Homo sapiens* (purple).

| Genome | Identifier | Repository site | Coverage | Contamination | Reference |
| --- | --- | --- | --- | --- | --- |
| Den D3 |  | http://cdna.eva.mpg.de/denisova/ | 31.59x | <0.5% | Meyer et al. 2012 |
| Altai Neandertal D5 | ENA PRJEB1265 | https://www.ebi.ac.uk/ena/browser/home | 52x | 1% | Prüfer et al. 2014 |
| Chagyrskaya 8 |  | http://ftp.eva.mpg.de/neandertal/Chagyrskaya/VCF/ | 28.36x | 0.7% | Mafessoni et al. 2020 |
| Mezmaiskaya 1 | ENA PRJEB1757 | http://cdna.eva.mpg.de/neandertal/Mezmaiskaya1/Pruefer_et al 2017 bam/ | 2.04 | <1% | Prüfer et al. 2017 |
| Spy 94a | ENA PRJEB21883 | https://www.ebi.ac.uk/ena/browser/home | 1.08 | 1.75% | Hajdinjak et al. 2018 |
| Vindija 33.19 | ENA PRJEB21882 | https://www.ebi.ac.uk/ena/browser/home | 31.32 | <1% | Prüfer et al. 2017 |
| Mezmaiskaya 2 | ENA PRJEB21881 | https://www.ebi.ac.uk/ena/browser/home | 1.85 | 0.83% | Hajdinjak et al. 2018 |
| Les Cottés Z4-1514 | ENA PRJEB21875 | https://www.ebi.ac.uk/ena/browser/home | 2.94 | 0.18% | Hajdinjak et al. 2018 |
| Goyet Q56.1 | ENA PRJEB21870 | https://www.ebi.ac.uk/ena/browser/home | 2.30 | 0.89% | Hajdinjak et al. 2018 |
| Ust'-Ishim | ENA PRJEB6622 | https://www.ebi.ac.uk/ena/browser/home | 42.72 | <0.13% | Fu et al. 2014 |
| Peștera Muierii 1 | ENA PRJEB33172 | https://www.ebi.ac.uk/ena/browser/home | 17.00 | 1.5% | Svensson et al. 2021 |
| Mal'ta | NCBI SRA SRP029640 | https://www.ncbi.nlm.nih.gov/sra/SRX346836[accn] | 1.21 | 1.6-2% | Raghavan et al. 2014 |
| Loschbour, Motala 12, Stuttgart | ENA PRJEB6272 | https://www.ebi.ac.uk/ena/browser/home | 22.13, 2.46, 19.28 | 0.4% (Loschbour, Stuttgart)<br>0.35% (Motala 12) | Lazaridis et al. 2014 |
| Human reference | GRCH38/hg38 | https://zenodo.org/records/5146236 |  |  | Schneider et al. 2017 |
| Human Blacklisted regions | GRCH38/hg38 | https://www.encodeproject.org/files/ENCFF356LFX/ |  |  | Amemiya et al. 2019 |

| Genome | Identifier | Total Amount of Used Reads | Total Amount of Mapped Reads after Filtering | Percentage of Reads that Mapped after Filtering |
| --- | --- | --- | --- | --- |
| Den 3 | ENA (PRJEB3092) | 4,789,230,696 | 1,329,869,335 | 27.77% |
| Altai Neandertal D5 | ENA (PRJEB1265) | 2,273,303,350 | 1,464,996,457 | 64.44% |
| Chagyrskaya 8 | http://cdna.eva.mpg.de/neandertal/Chagyrskaya/ | 1,208,110,335 | 984,904,699 | 81.52% |
| Mezmaiskaya 1 | ENA (PRJEB21195) | 1,401,739,418 | 131,887,968 | 9.41% |
| Spy 94 a | ENA (PRJEB21883) | 72,809,832 | 66,872,902 | 91.85% |
| Vindija 33.19 | http://cdna.eva.mpg.de/neandertal/Vindija/bam/Pruefer_etal_2017/Vindija33.19/ | 1,754,339,206 | 1,353,844,718 | 77.17% |
| Mezmaiskaya 2 | ENA (SAMEA104168735, SAMEA104168734, SAMEA104168736) | 111,777,820 | 103,837,327 | 92.90% |
| Les Cottés Z4-1514 | ENA (PRJEB21875) | 189,224,733 | 166,455,607 | 87.97% |
| Goyet Q56.1 | ENA (PRJEB21870) | 116,417,514 | 109,663,410 | 94.20% |
| Ust'-Ishim | ENA (PRJEB6622) | 2,039,353,632 | 1,572,569,967 | 77.11% |
| Peștera Muierii 1 | ENA (PRJEB33172) | 611,030,108 | 498,336,516 | 81.56% |
| Mal'ta | NCBI (SRA SRP029640) | 79,765,126 | 40,065,643 | 50.23% |
| Loschbour | ENA (PRJEB6272) | 862,379,535 | 693,299,437 | 80.39% |
| Motala 12 | ENA (PRJEB6272) | 94,818,771 | 94,463,217 | 99.63% |
| Stuttgart | ENA (PRJEB6272) | 791,983,938 | 643,277,473 | 81.22% |

**Table S2.** Description of the fossils of the 15 individuals whose genomes are analyzed in this study. From top to bottom, genomes are listed according to dating from the oldest to the most recent within each population: Denisovan (grey), Neandertal (orange), *Homo sapiens* (purple).

| Fossil remains | Fossil details<br>*anatomy*<br>sex | Geographical location | Cave and layer | Time period (ybp - calibrated age) | Relevant publications for the genome published |
| --- | --- | --- | --- | --- | --- |
| Denisova D3 | *Distal phalanx of the fifth finger.*<br>Female | Central Asia: Central Siberia, Altai Moutain | Denisova Cave Layer 11.2, East Gallery | 76200-51600 | Meyer et al. Science 2012<br>Reich et al. Nature 2010<br>Douka et al. Nature 2019 |
| Altai Neandertal D5 | *Phalanx foot toe*<br>Female | Central Asia: Central Siberia, Altai Moutain | Denisova Cave Layer 11.4 | 120000 | Prüfer et al. Nature 2014 |
| Chagyrskaya 8 | *Distal manual phalanx*<br>Female | Central Asia: Central Siberia, Charysh River, foothills of the Altai Moutain. | Chagyrskaya Cave | 80000-60000 | Mafessoni et al. PNAS 2020 |
| Mezmaiskaya 1 | *Almost complete skeleton*<br>Still breast-feeding Neonate | Eastern Europe: Southern Russia, foothills of the North Caucasus | Mezmaiskaya Cave | 70000-60000 | Prüfer et al. Science 2017 |
| Spy 94a | *Upper right molar associated to a maxillary fragment*<br>Male | Western Europe: Belgium, Namur province | Cave | 49800-42300 | Haidinjak et al. Nature 2018<br>Devièse et al. PNAS 2021 |
| Vindija 33.19 | *Fragments of long bones*<br>Female | Southern Europe: Croatia, Dinaric Alps | Vindija Cave | 47400-44300 | Prüfer et al. Science 2017 |
| Mezmaiskaya 2 | *Skull fragment*<br>Male? | Eastern Europe: Foothills of the North Caucasus | Mezmaiskaya Cave | 44.600-42.960 | Haidinjak et al. Nature 2018 |
| Les Cottés Z4-1514 | *Tooth*<br>NA | Western Europe: France, Saint-Pierre-de-Maillé | “Les Cottés” cave | 43200-42500 | Haidinjak et al. Nature 2018 |
| Goyet Q56-1 | *Right Femur*<br>NA | Western Europe: Belgium, Namur province | “third cavern” of Goyet cave | 42800-42100 | Haidinjak et al. Nature 2018 |
| Ust'-ishim | *Femora*<br>Male | Central Asia: North-Western Siberia, Omsk province | Cave on the bank of the Irtysh River | 45930-42900 | Fu et al. Nature 2014 |
| Peștera Muierii 1 | *Temporal bone*<br>Female | Eastern Europe: Romania, Carpatian mountains | “Women’s Cave” | 35257 +/-259 | Svensson et al. Current Biology 2021 |
| Malt'a 1 | *Skeleton burial*<br>Male | Central Asia: Central - Eastern Siberia, Bolshaya Belaya river | Open air site | 25600 | Raghavan et al. Nature 2014<br>Fiedel & Kuzmin, 2007 |
| Loschbour | *Skeleton*<br>Male | Central Europe: Luxembourg, Mullerthal | Rock shelter | 8200-7900 | Lazaridis et al. Nature 2014 |
| Motala 12 | *Maxilla (burial)*<br>Male | Northern Europe: Sweden, Motala | Necropolis - Kanaljorden | 8000-7600 | Lazaridis et al. Nature 2014 |
| Stuttgart | *Tooth (burial)* | Central Europe: South-Western, Germany | Stuttgart-Mühlhausen | 7000 | Lazaridis et al. Nature 2014 |

**Table S3.**
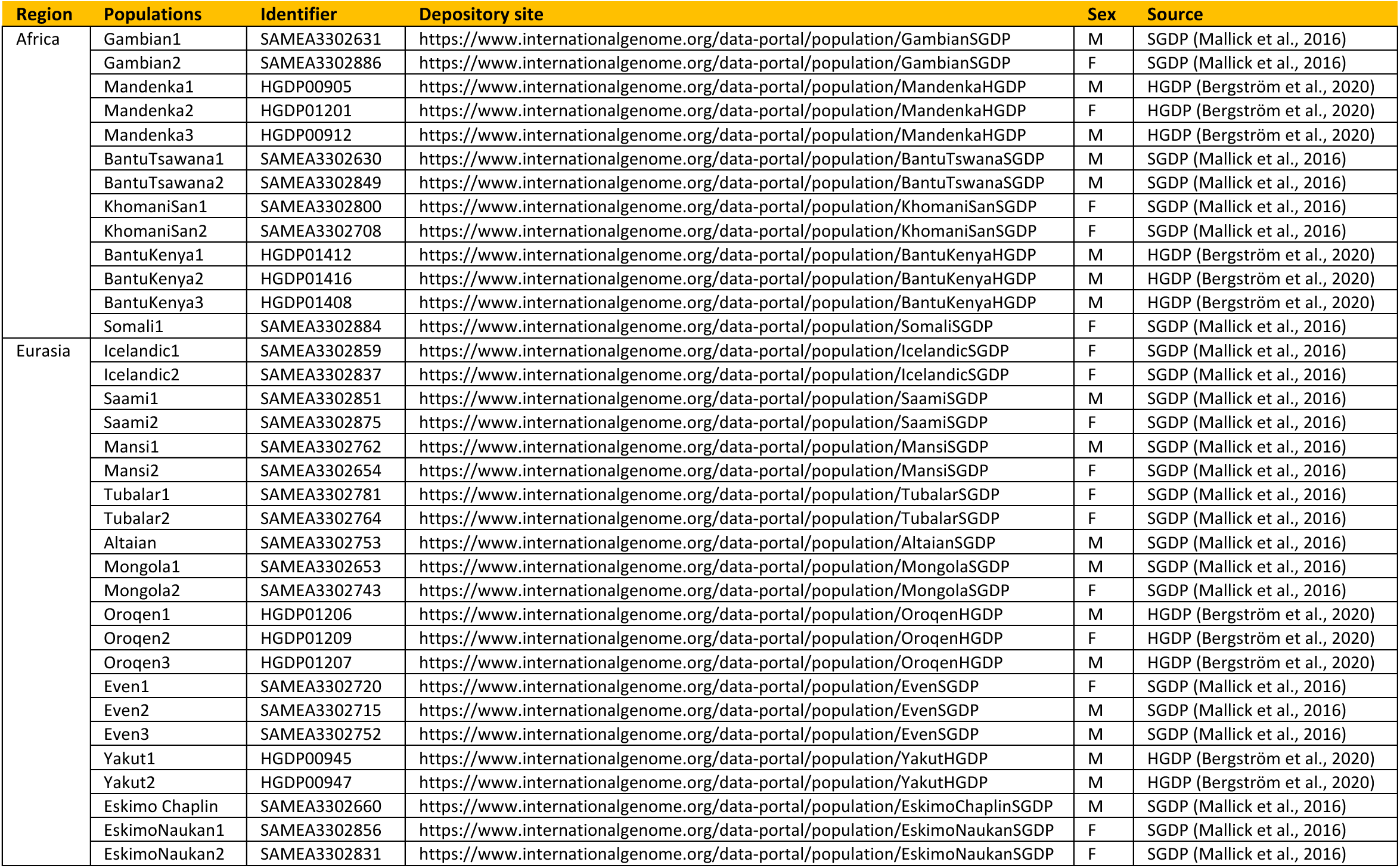
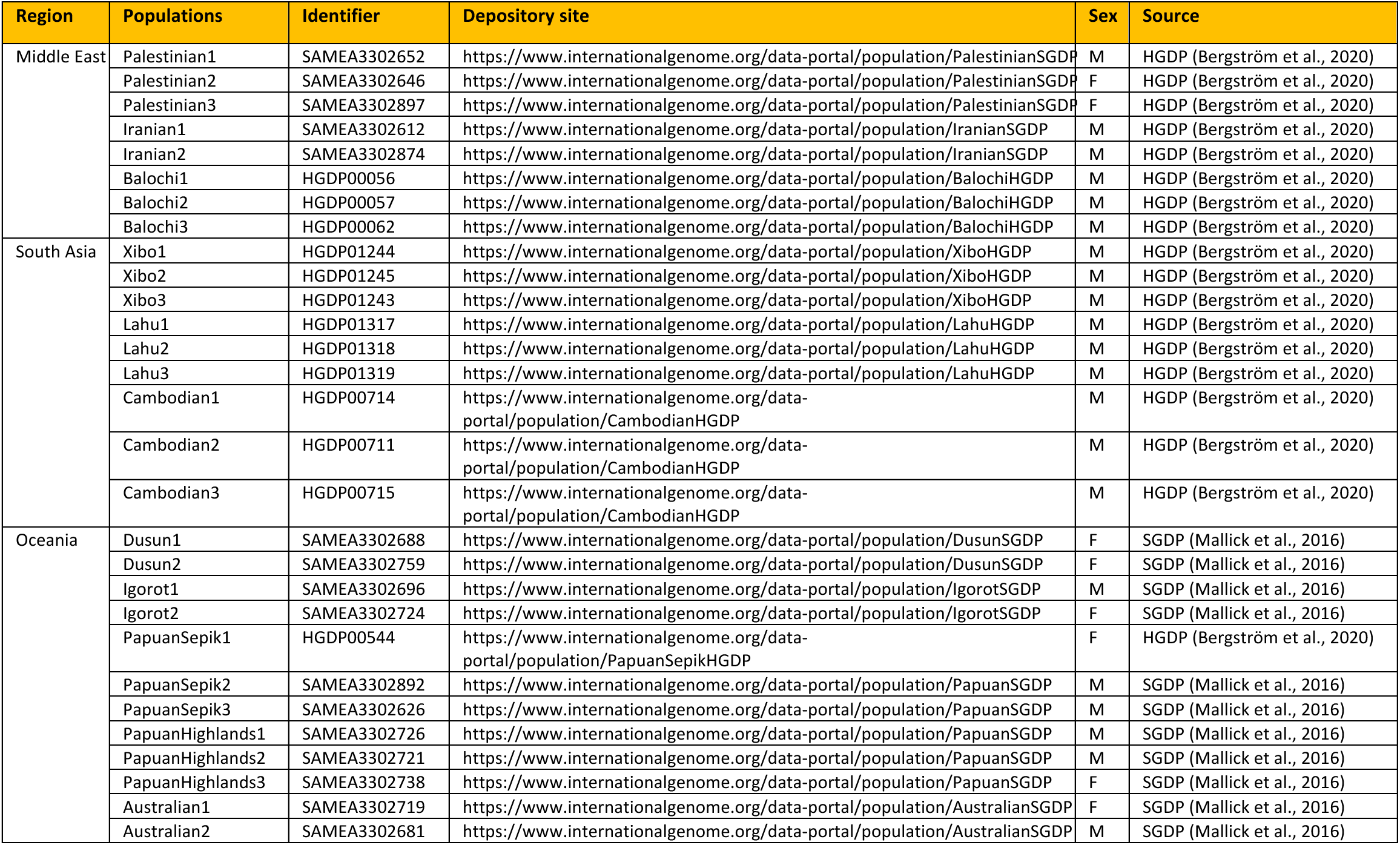
Dowloading information for the 64 modern genomes, organised in 5 populations (Africa, Eurasia, iddle East, South Asia and Oceania), in this study.

**Table S4.**
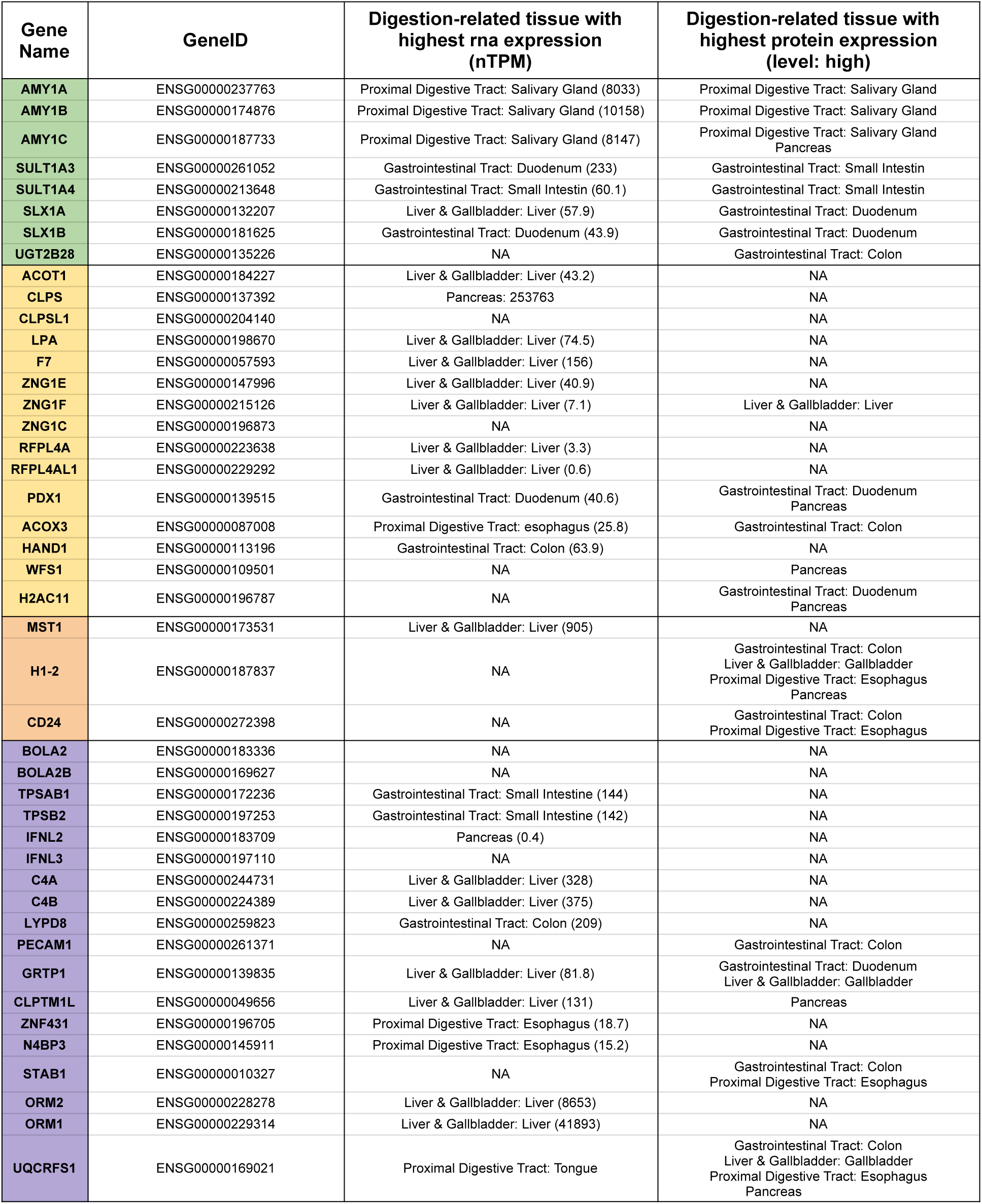
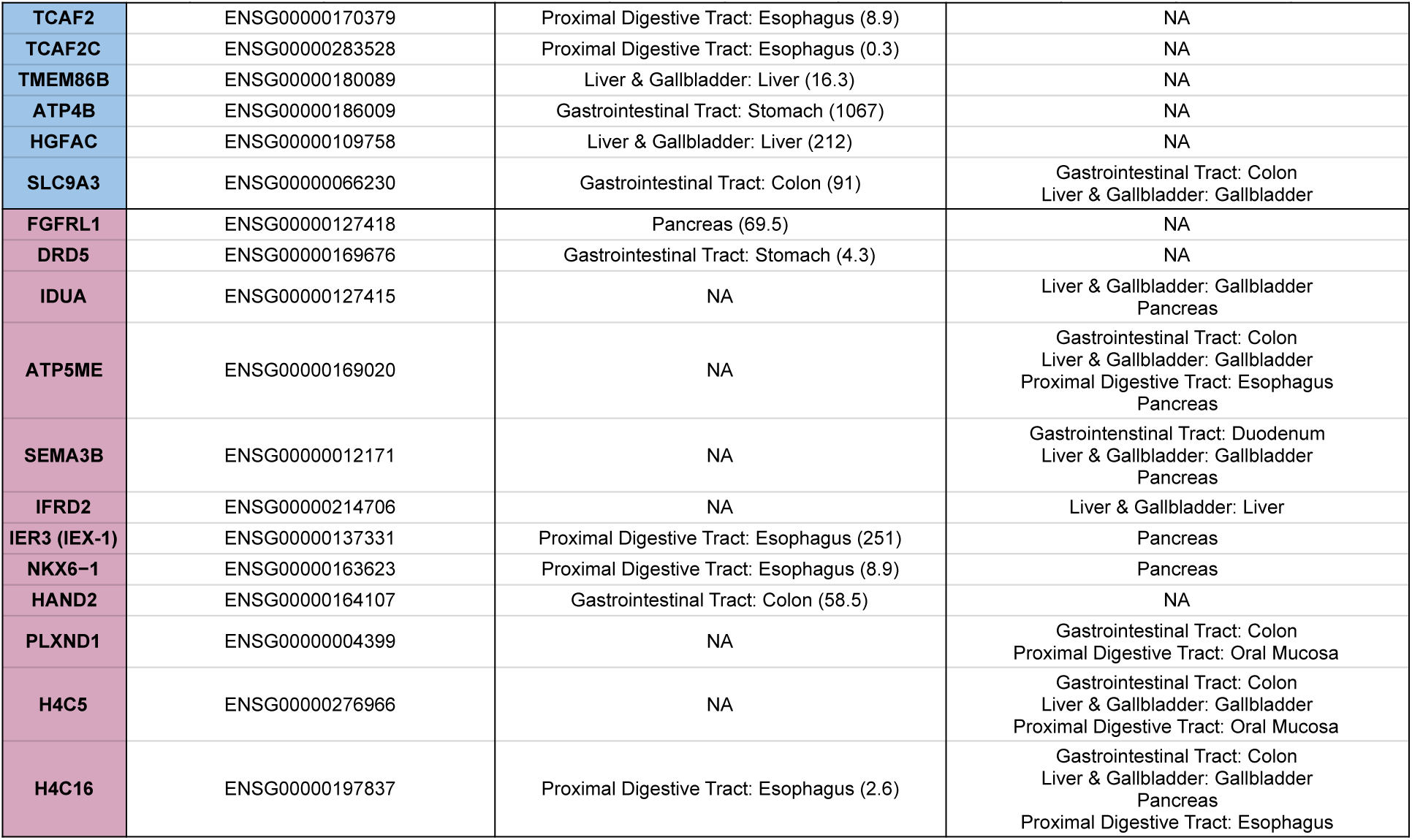
Primary and secondary tissues expressing the 50 genes. The genes listed in **Table 1** and **Table 2** are expressed across various tissues. For gene clusters, each gene is considered individually, totalling 62 genes in this table. We specify the tissues where each gene or gene copy in a cluster shows strong expression, with RNA and protein expression data in columns three and four, respectively. This information is sourced from the Human Protein Atlas Database (www.proteinatlas.org/), which provides RNA and protein expression data alongside additional functional details. Ensembl gene ID are listed in the second column, and genes are color coded according to **Tables 1** and **2**.

**Table S5.**
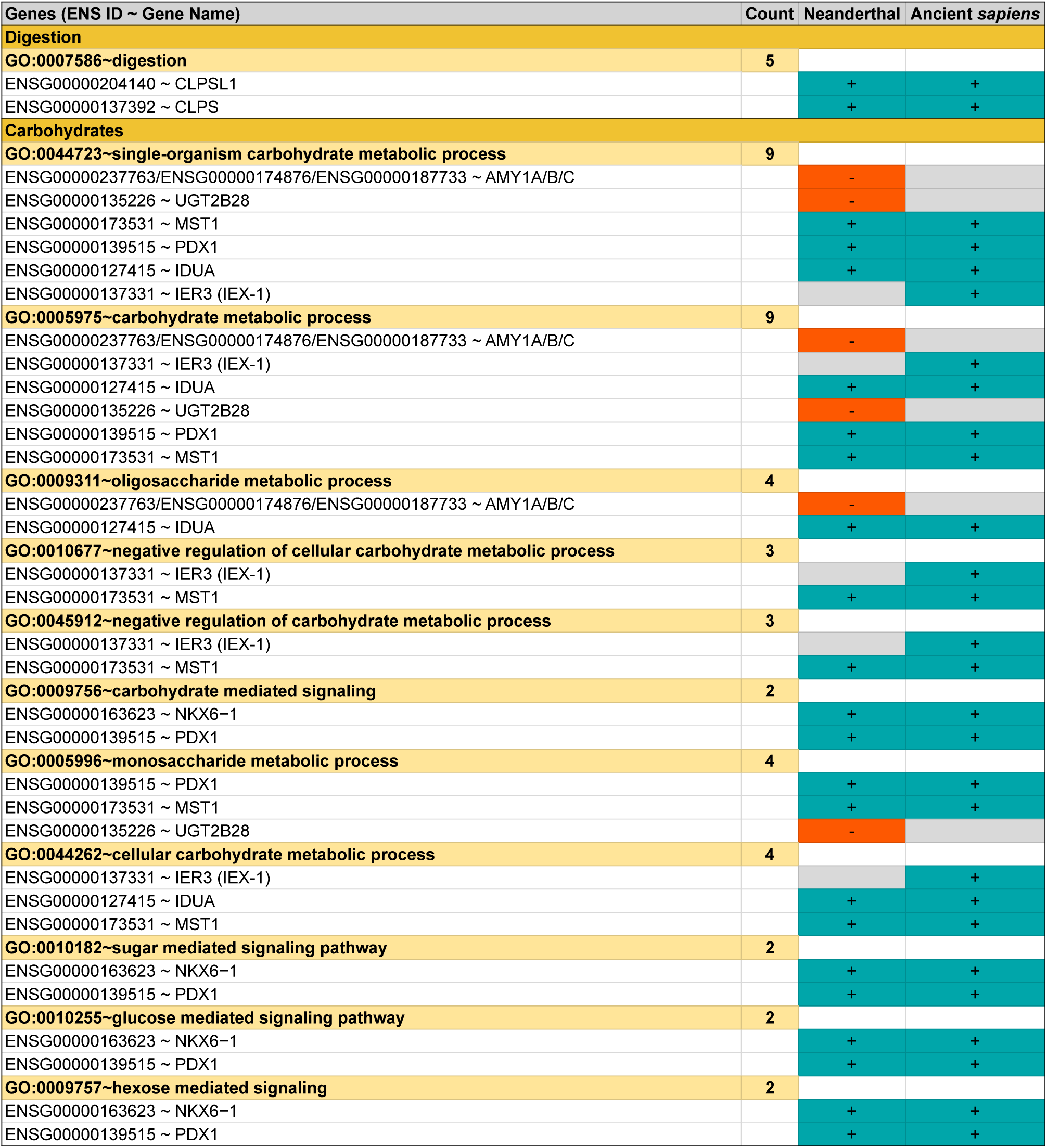
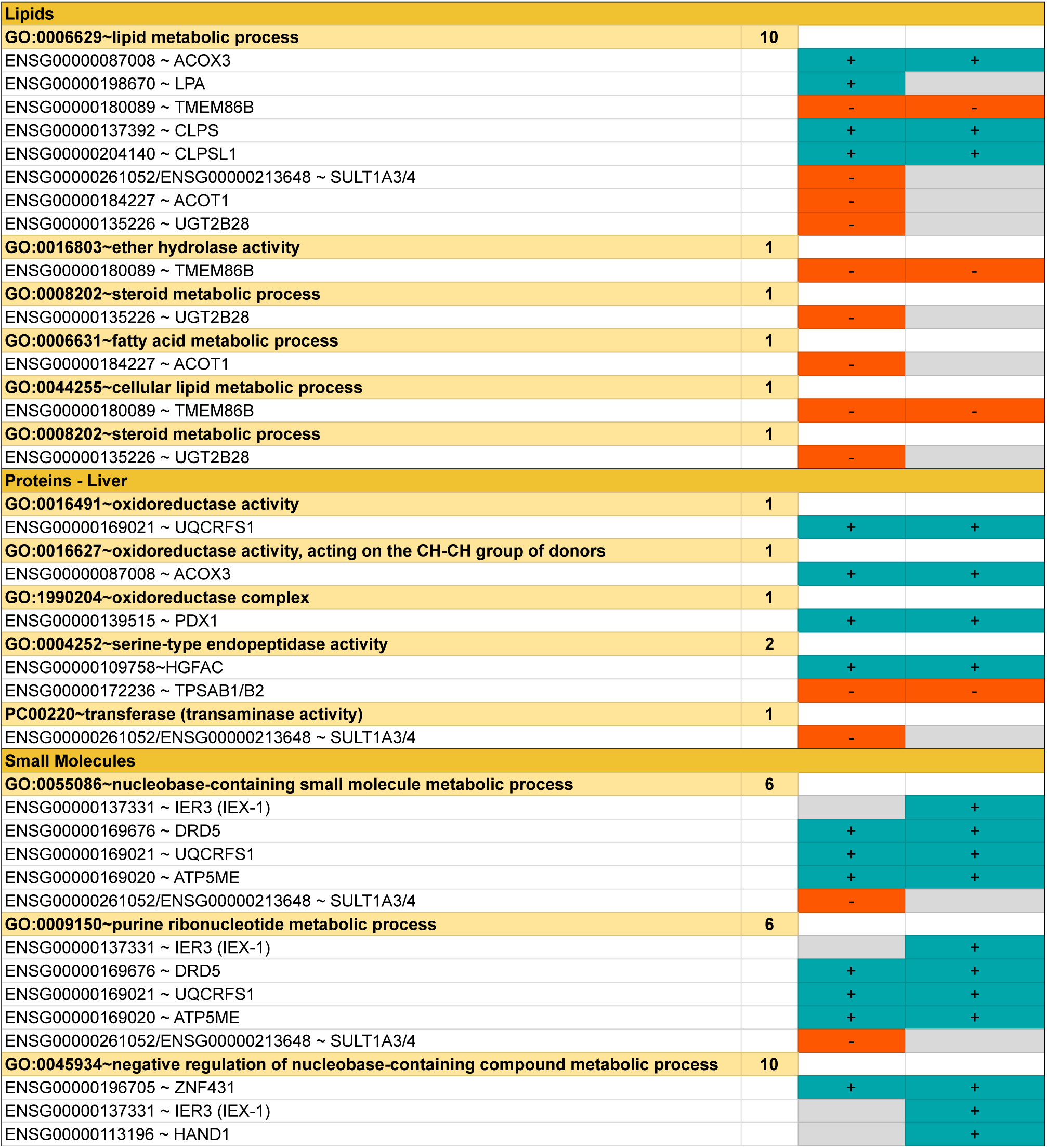
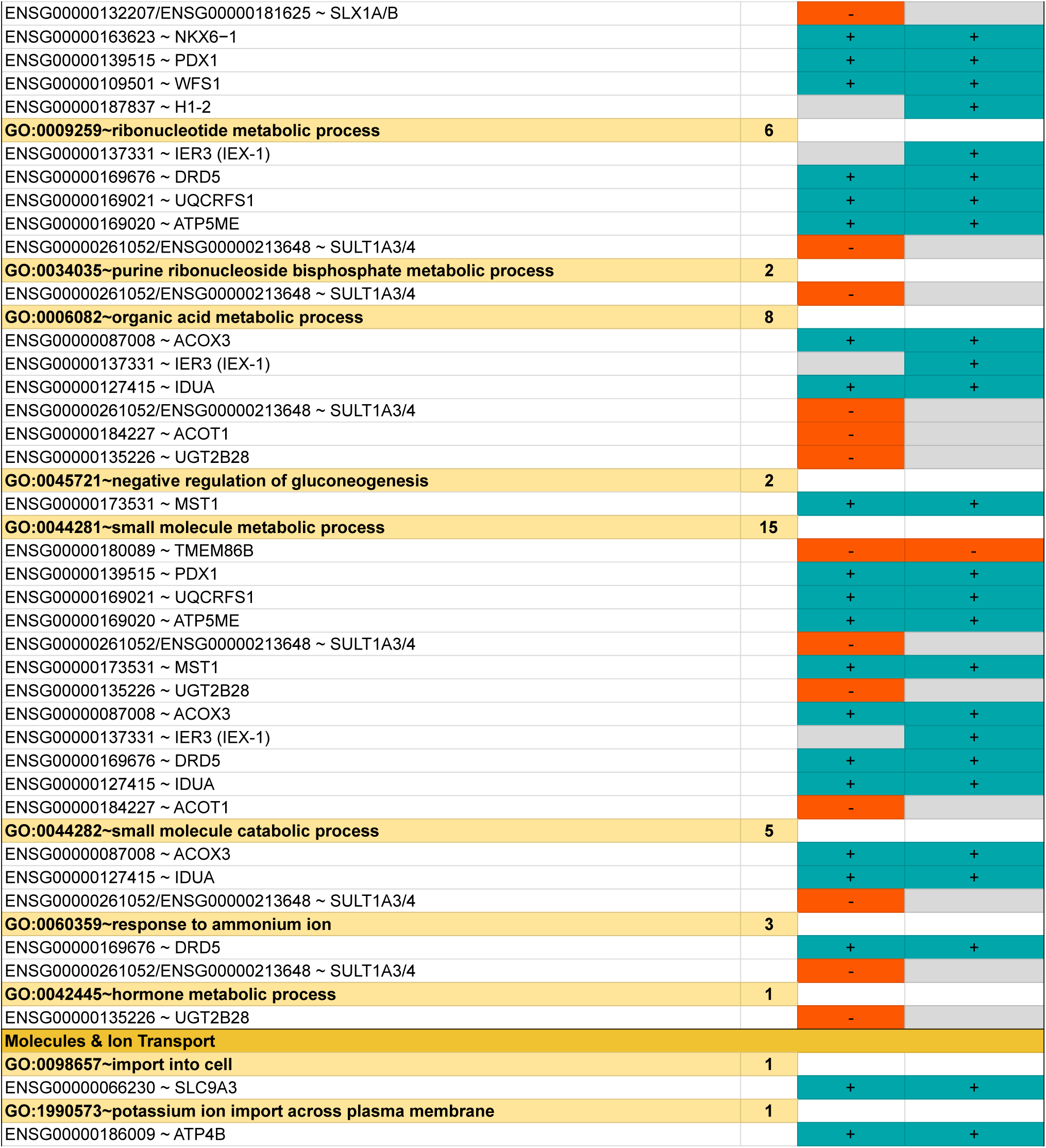
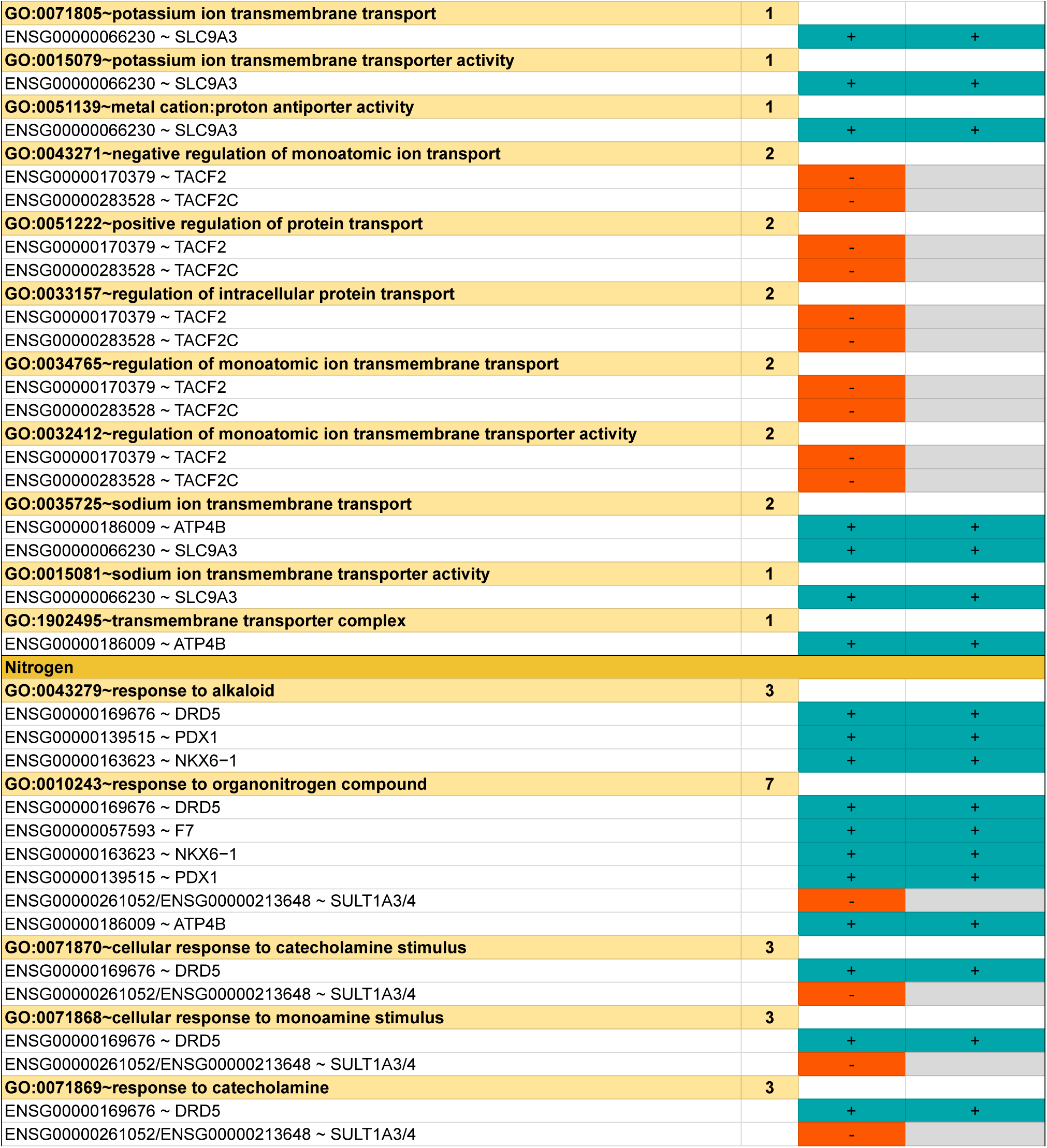
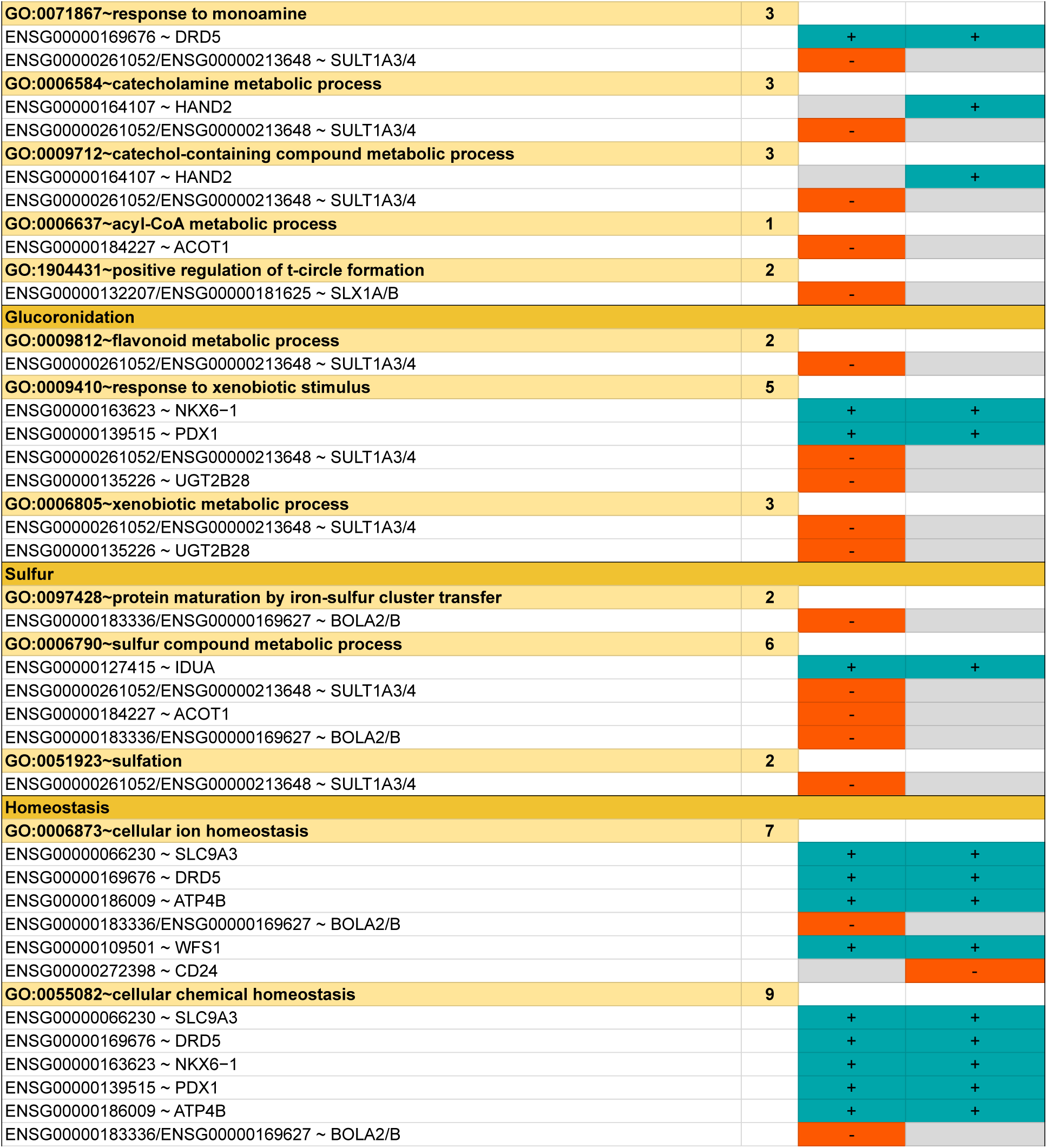
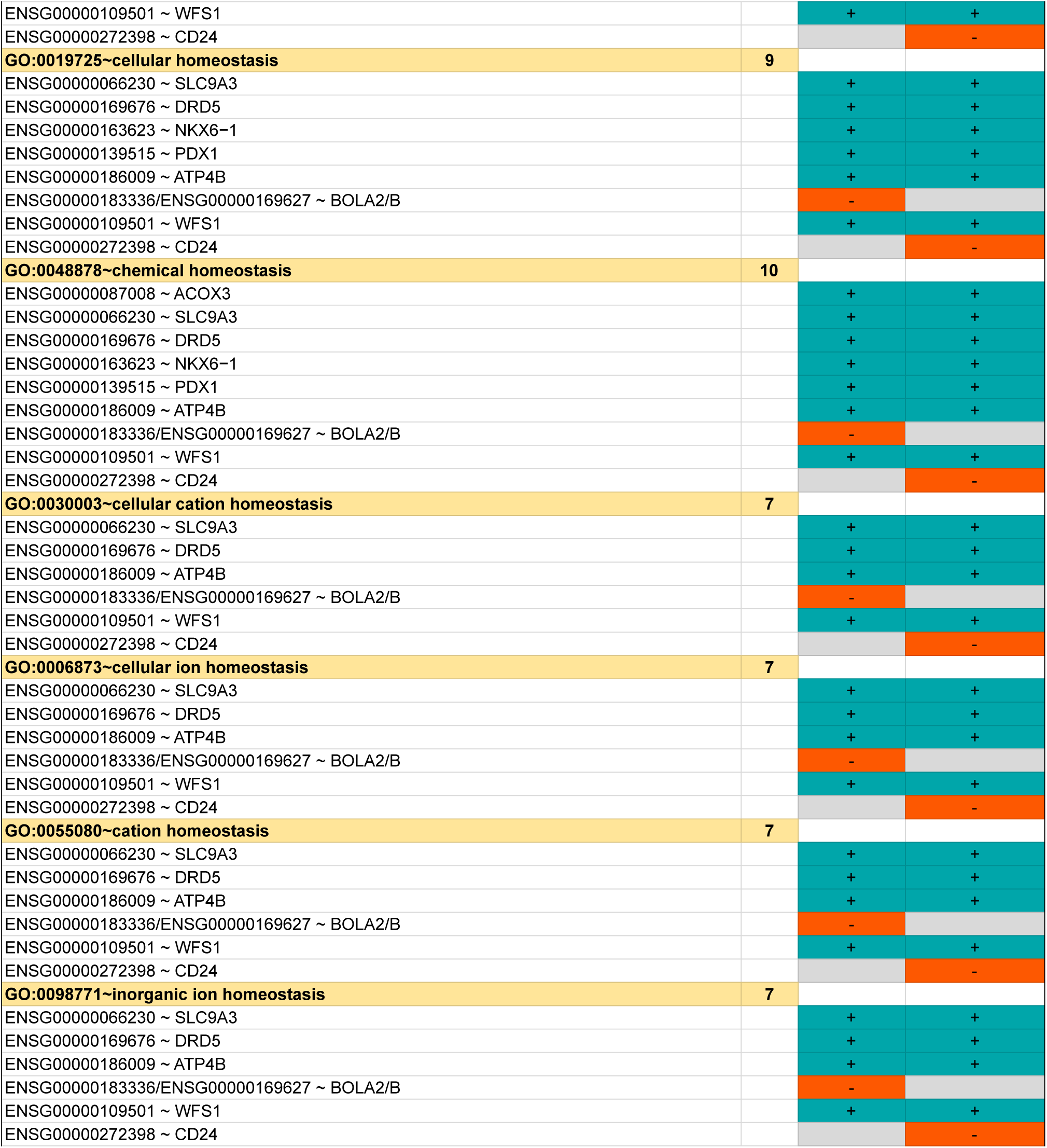
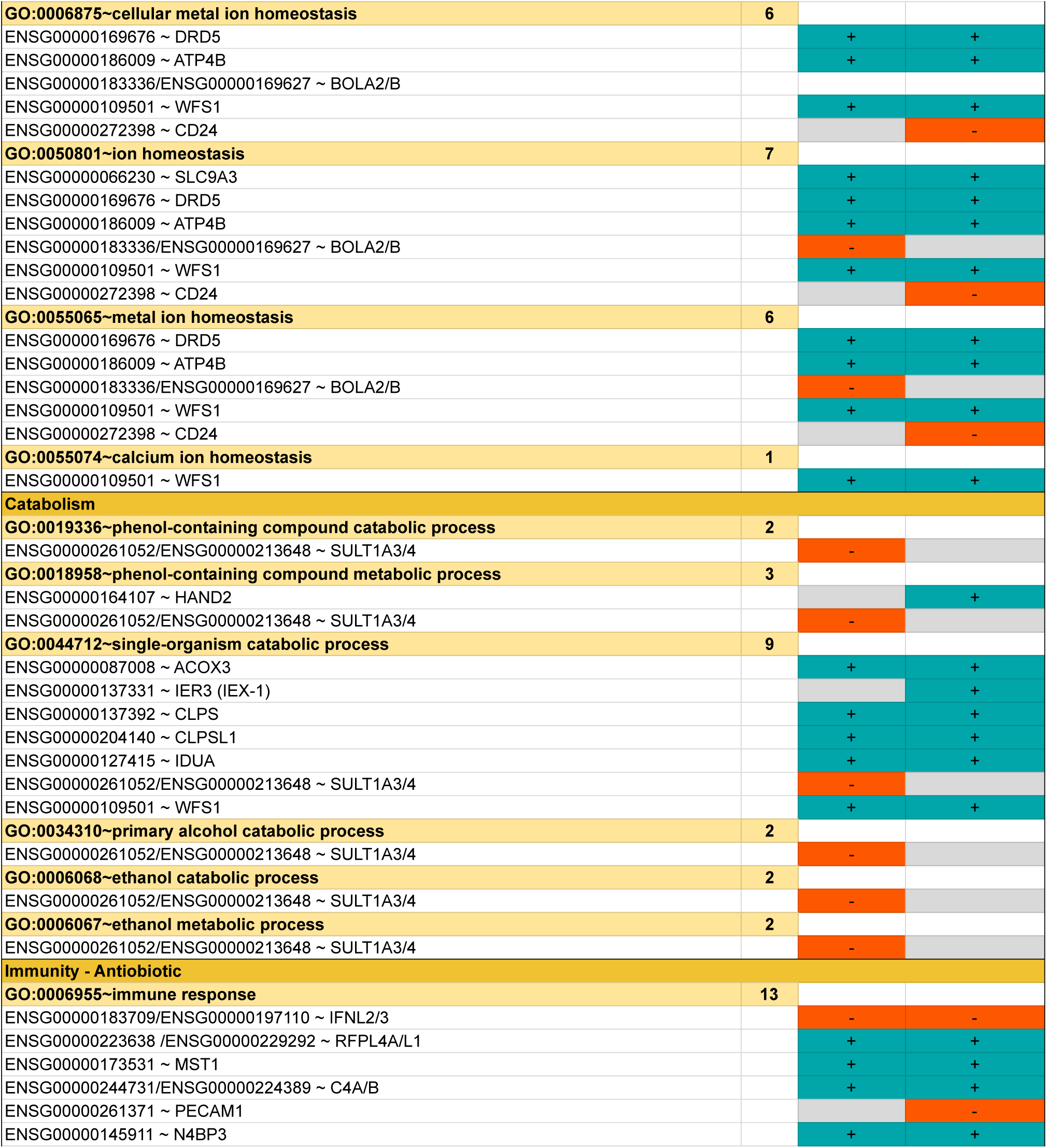
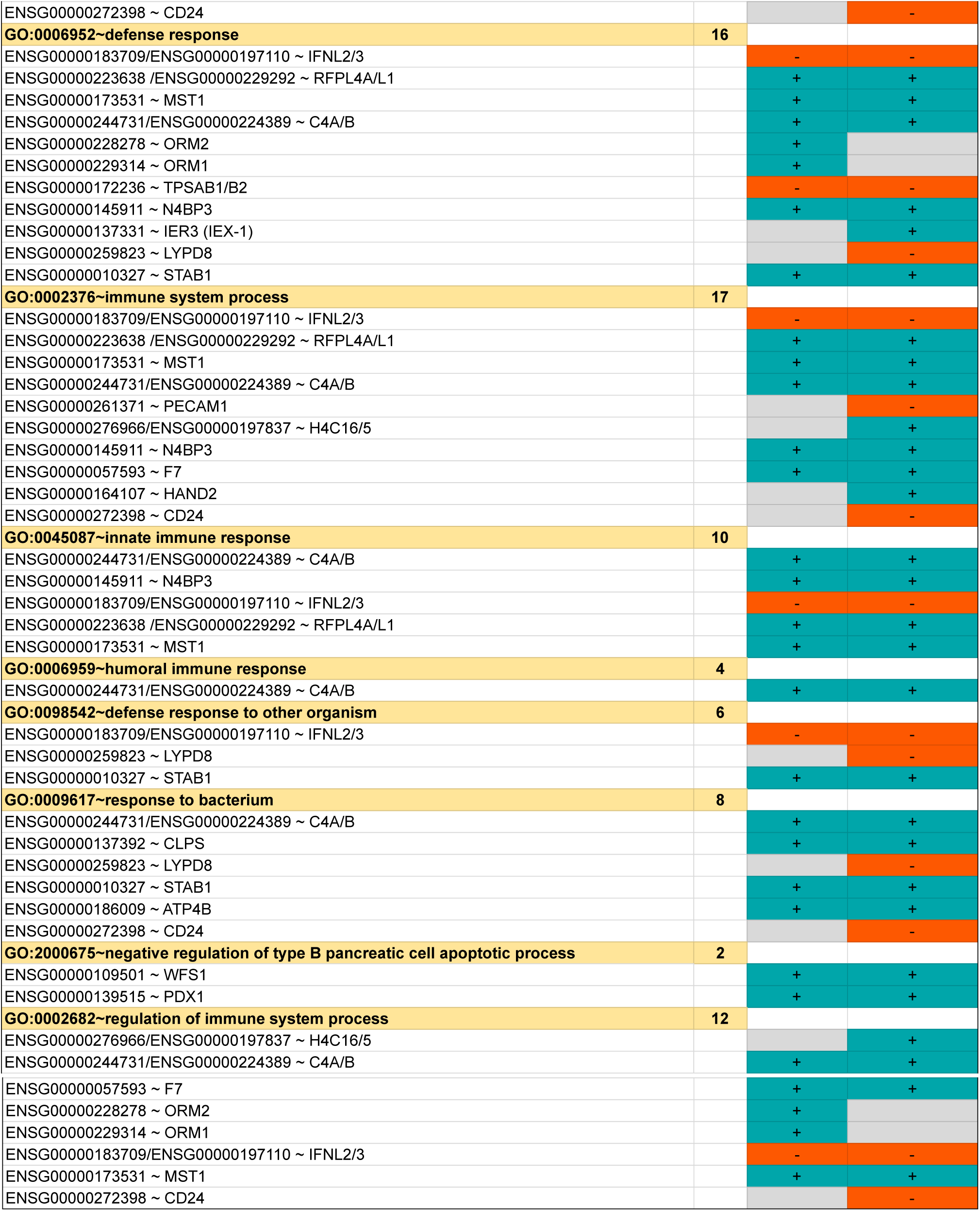
Gene Ontology terms associated with the 50 genes presenting differential CNV in archaic populations compared to the reference human genome. The first column report the GO term and the GO description for the genes presenting a gain (third column) or loss (fourth column) of copy number in *Hn* and *Hs* populations compared to the reference human genome. The genes ZNG1E/F/C, H2AC11, GRTP1, CLPTM1L, FGFRL1, SEMA3B, IFRD2, and PLXND1 are currently missing from any documented digestion-related GO term and are not listed in the table.

**Table S6.**
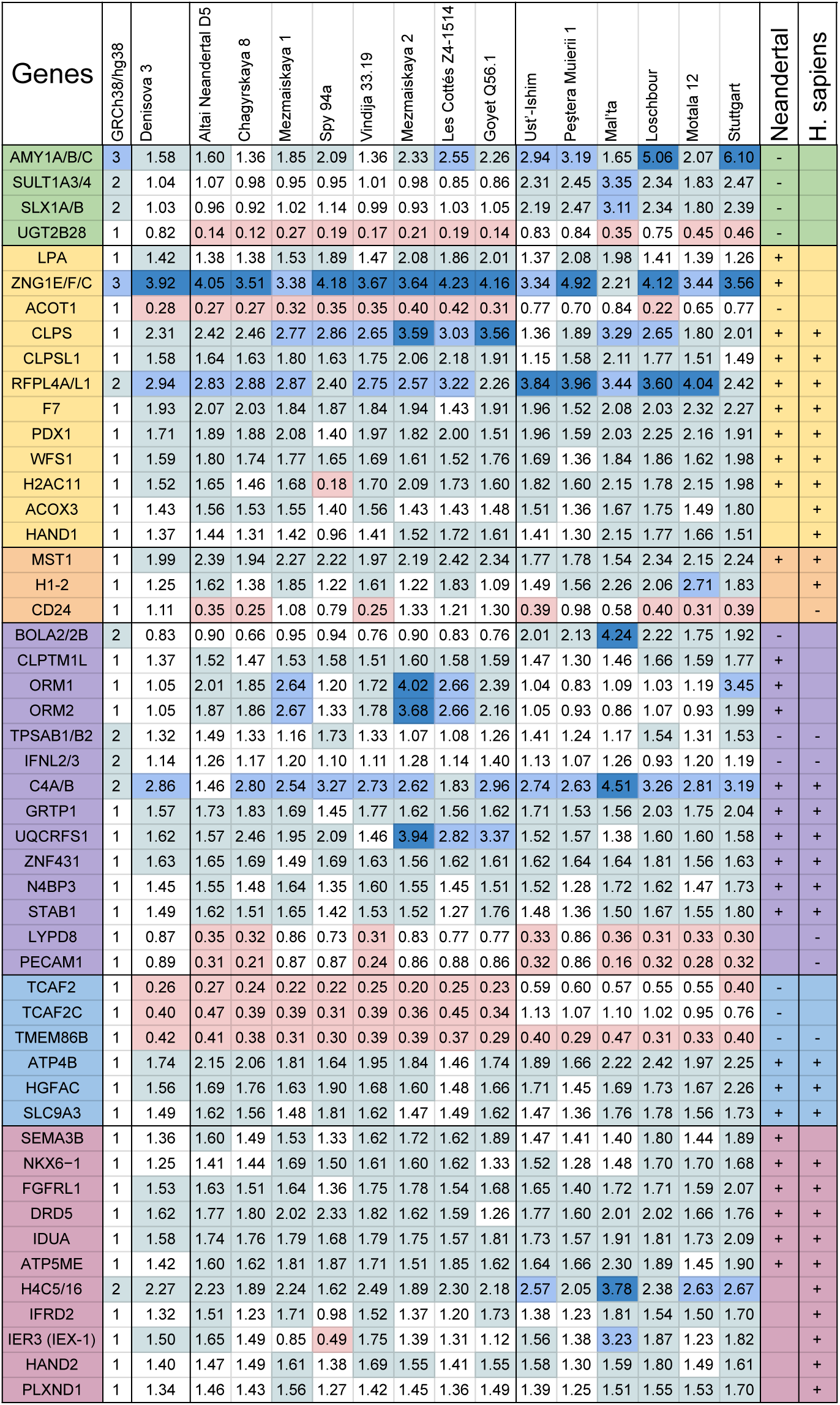
Estimated haploid copy number for genes related to digestion in ancient genomes and human population CNV trends (Complement to. **Table 1 and Table 2).** From left to right, the exact estimated gene copy number is reported for all genes and all genomes from *Den*, *Hn* and *Hs* populations. Genomes within a population are ordered from the oldest to the youngest. Blue shades correspond to the number of gene copies after mapping sequencing reads on the reference genome of modern humans GRGh38/hg38. Colors as in **Tables 1** and **2**.

**Table S7.**
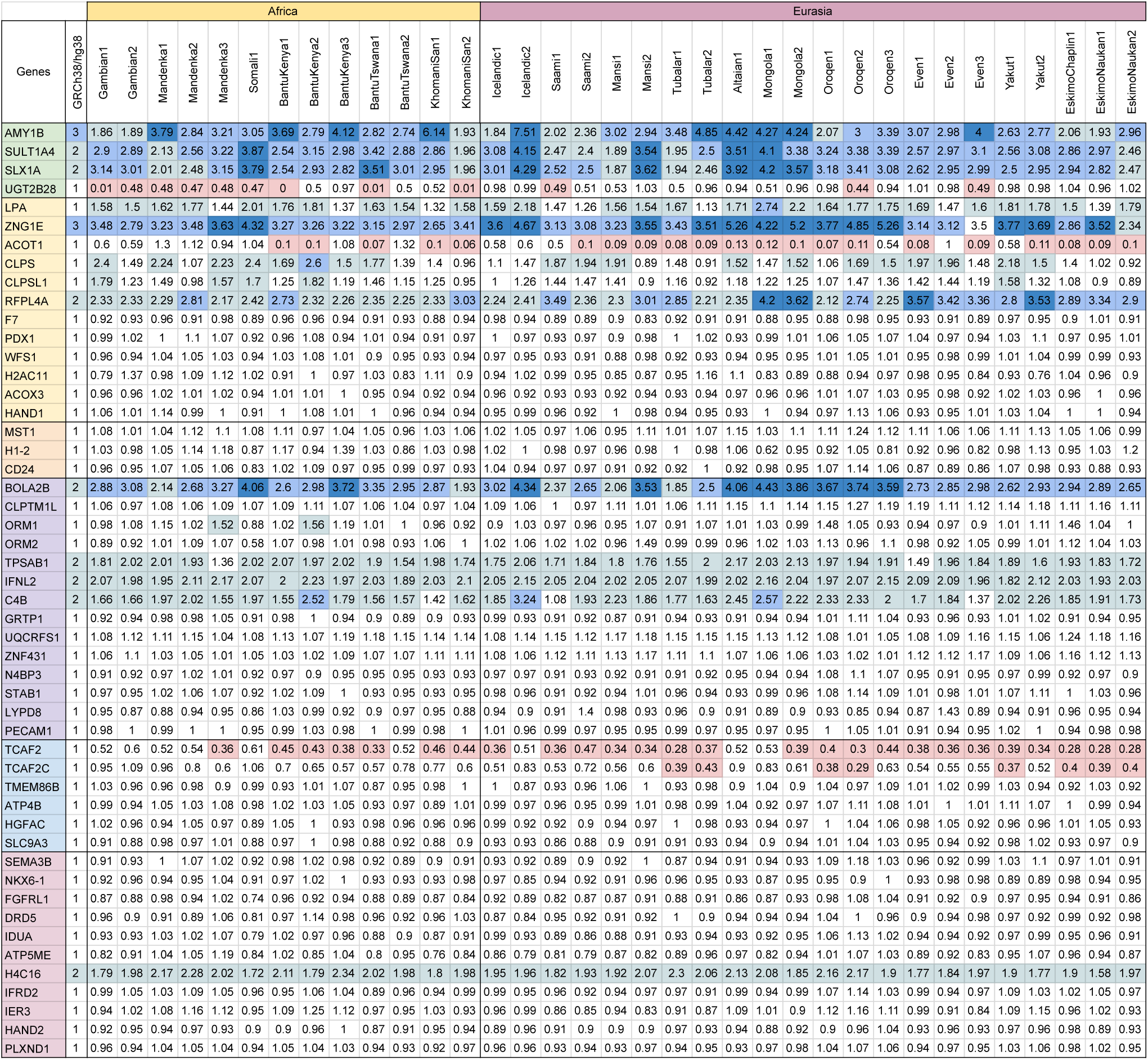
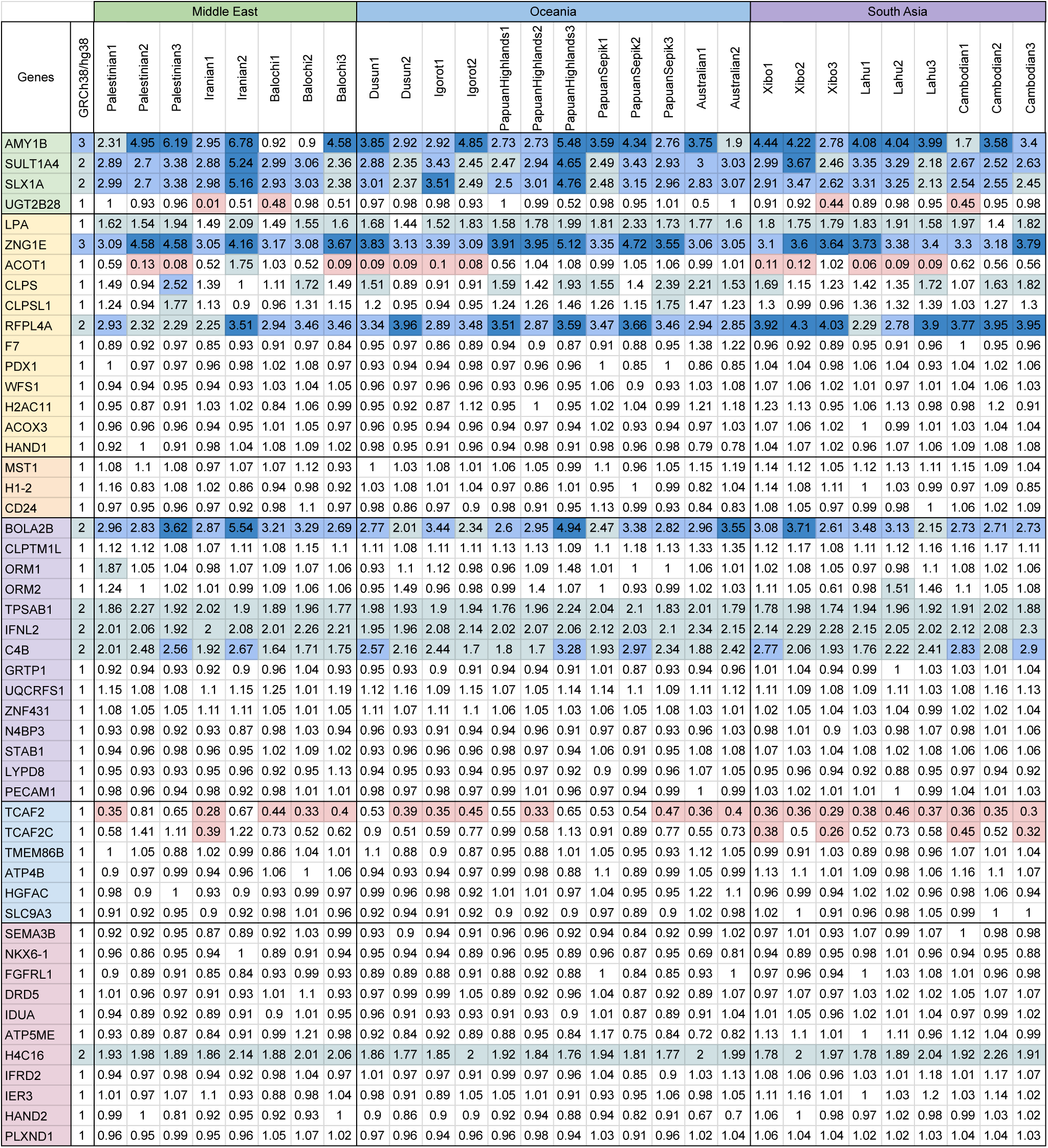
Estimated haploid copy number for genes related to digestion in modern genomes (Complement to. **Table 1 and Table 2).** From left to right, the exact estimated gene copy number is reported for all genes and all genomes from ethnic groups in Africa and Eurasia. Genomes within a population are ordered by ethnic group. Blue shades correspond to the number of gene copies after mapping sequencing reads on the reference genome of modern humans GRGh38/hg38. Colors as in **Tables 1** and **2**.

## Supplementary Figures

**Figure S1.**
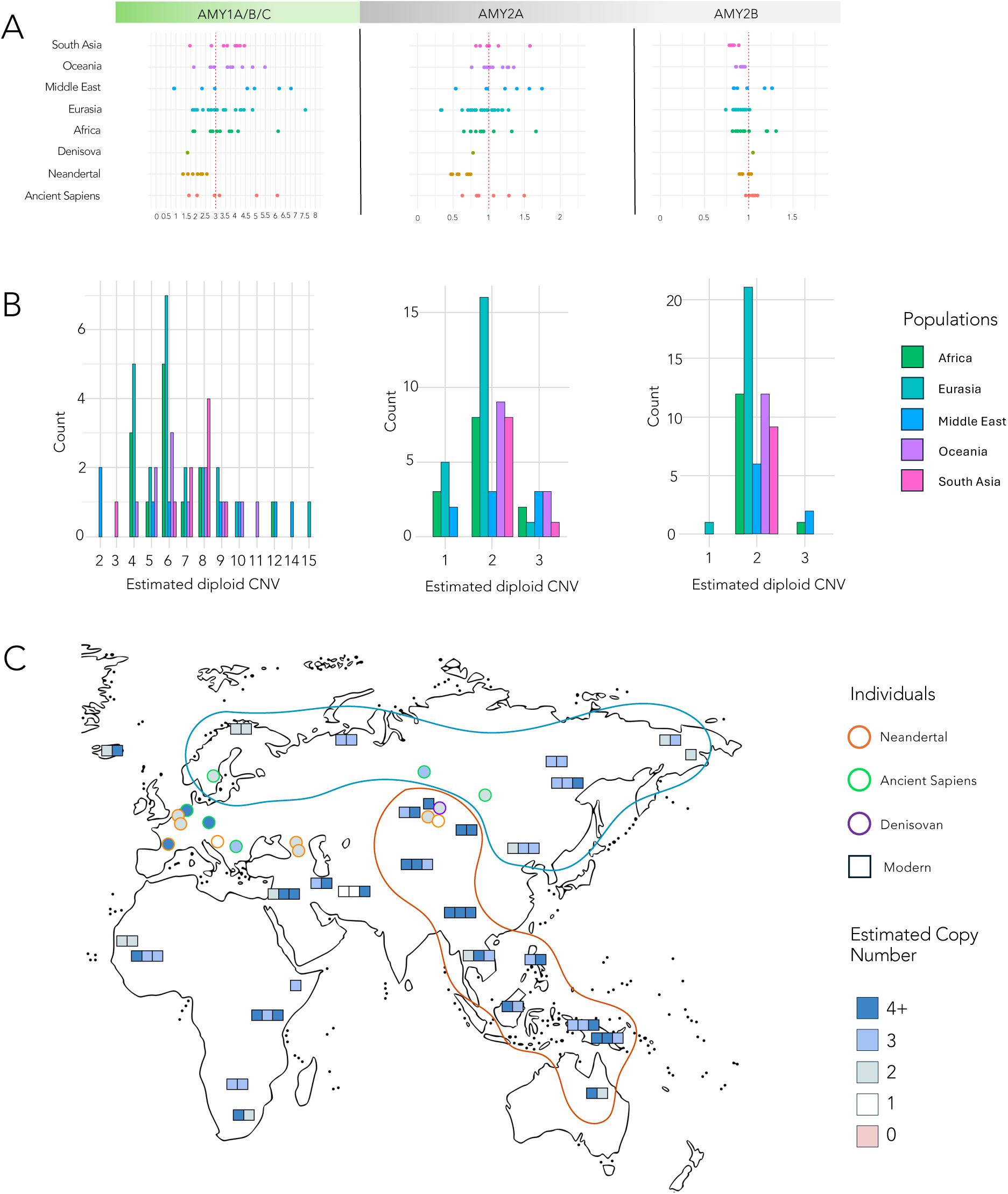
CNV estimations of the *AMY1A/B/C* gene cluster, and the *AMY2A* and *AMY2B* genes in ancient and modern populations. **A.** Haploid estimation of CNV for the *AMY1A/B/C* gene cluster, along with the *AMY2A* and *AMY2B* genes. Note that the CNV for the *AMY2A* and *AMY2B* genes is estimated to be 1. (Complement to Figure 3.) **B.** Distribution of diploid copy number for the 64 modern individuals analyzed in this study. Diploid CNV was calculated using 2x read estimates for each gene rounded to the nearest integer. As expected, pair diploid CNVs show higher numbers of individuals, as remarked in (Usher *et al.* 2015) **C.** Visualisation of *AMY1A/B/C* haploid CNVs (colors as shown in **Tables 1** and **2**) for both modern (square) and ancient (circle) individuals. Different populations are identified by colored borders: *Hn* in orange, *Hs* in green, *Den* in purple, and modern individuals in black. Most modern genomes from the Eurasian belt region (blue shape) exhibit low CNVs, similar to ancient individuals, while modern genomes from the East Asia, South Asia, and Oceania corridor (red shape) show higher CNVs. The CNV for each individual within the ethnic groups considered in this study is represented by a colored square (compare to Figure 1B).

**Figure S2.**
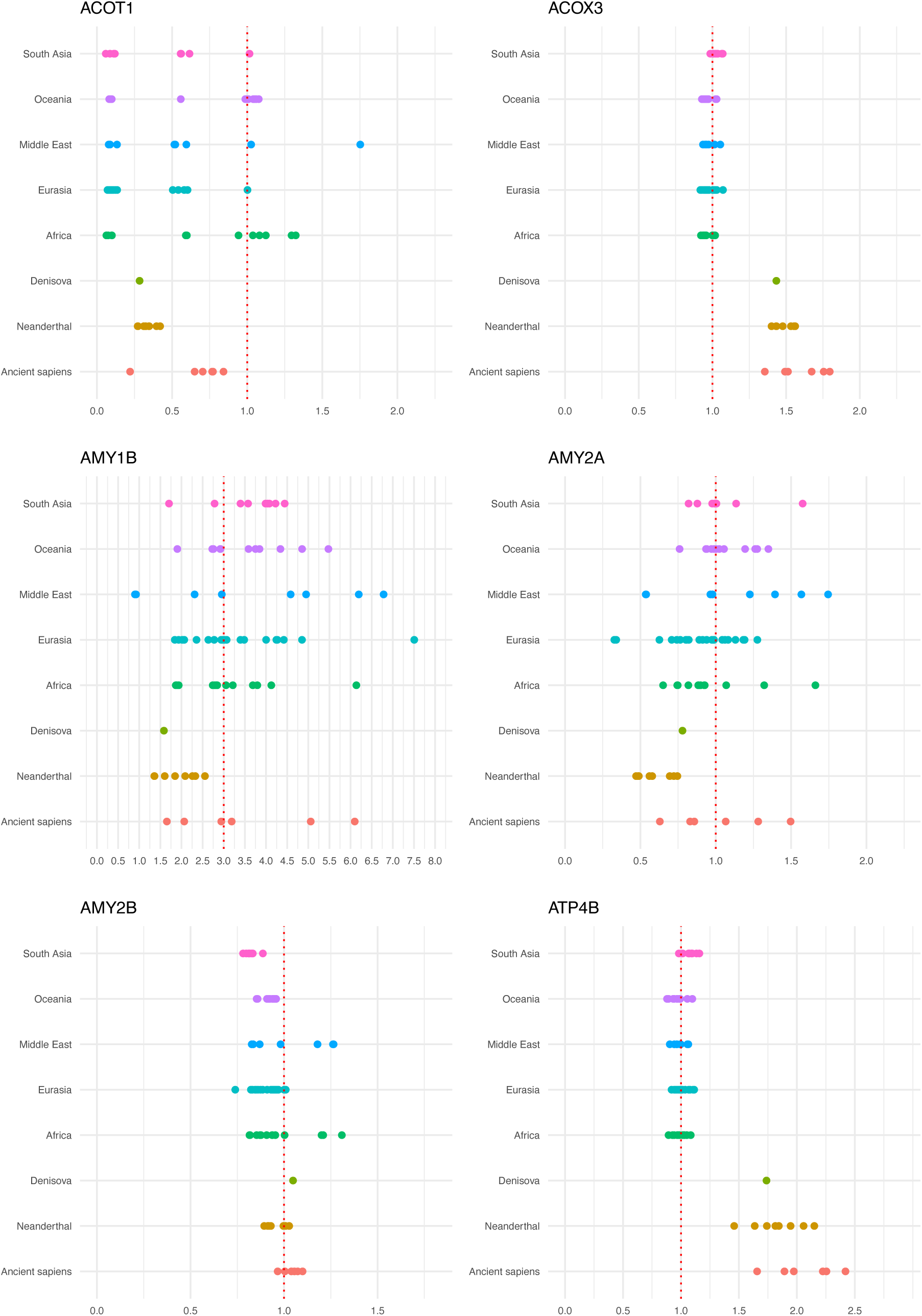
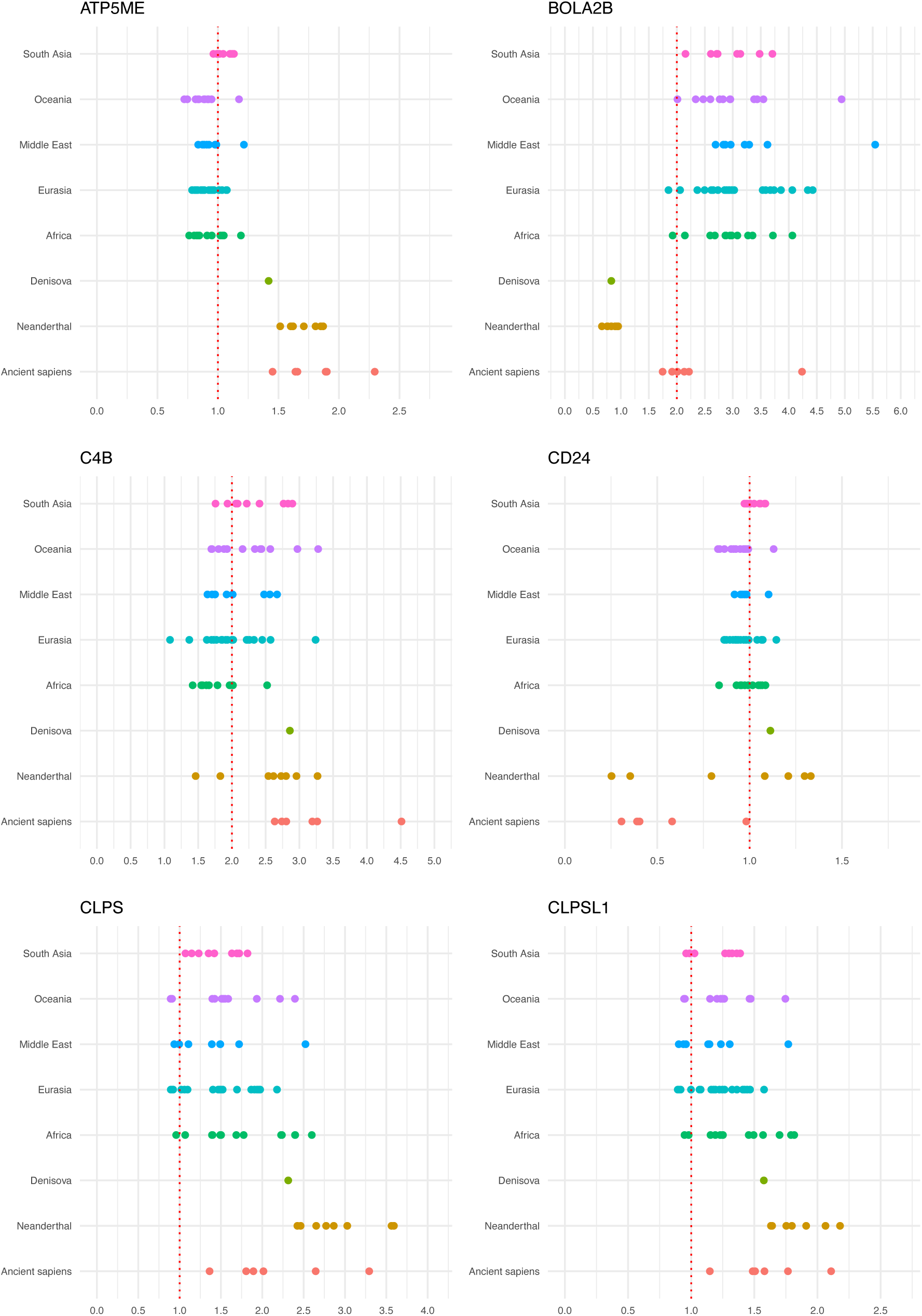
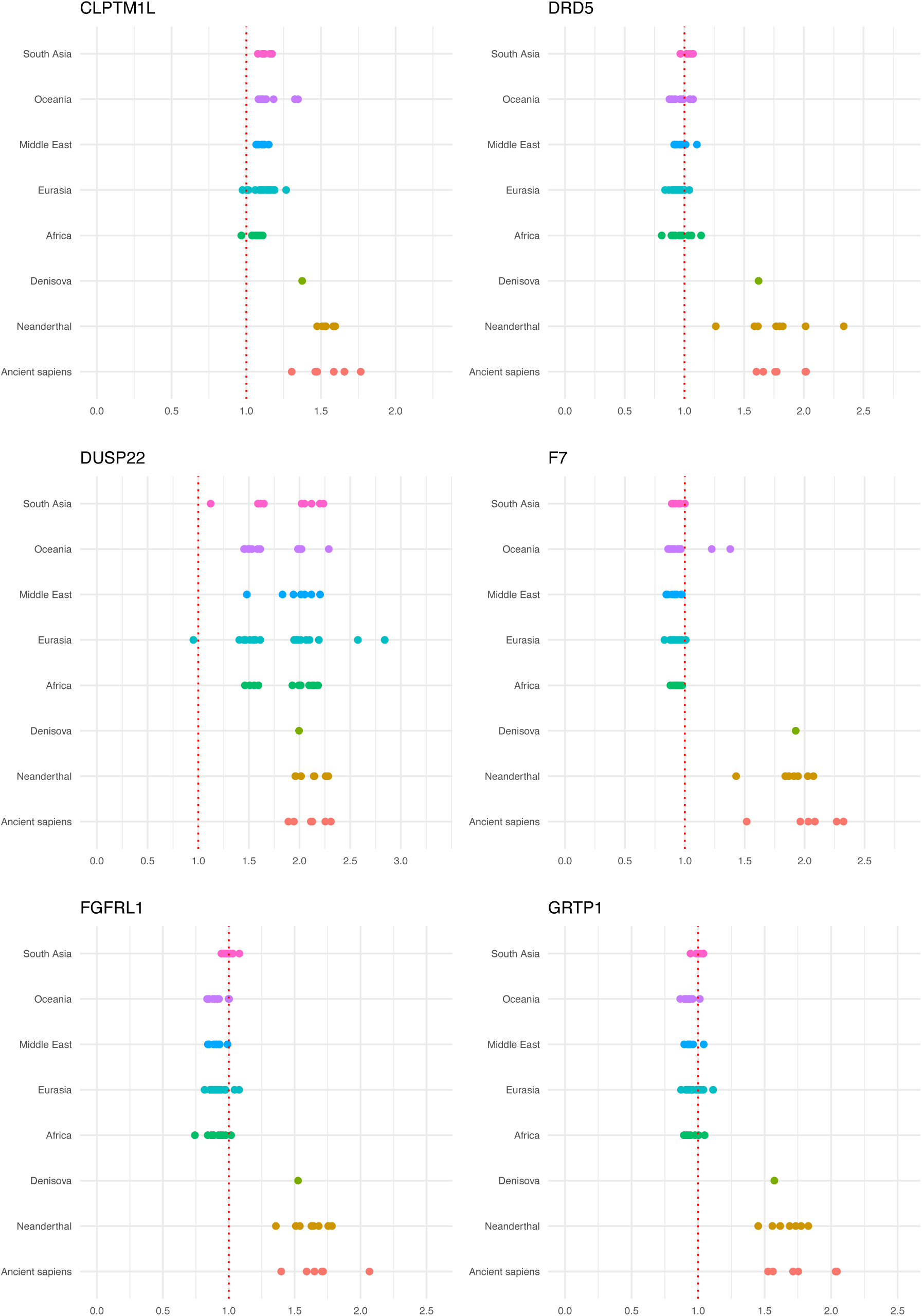
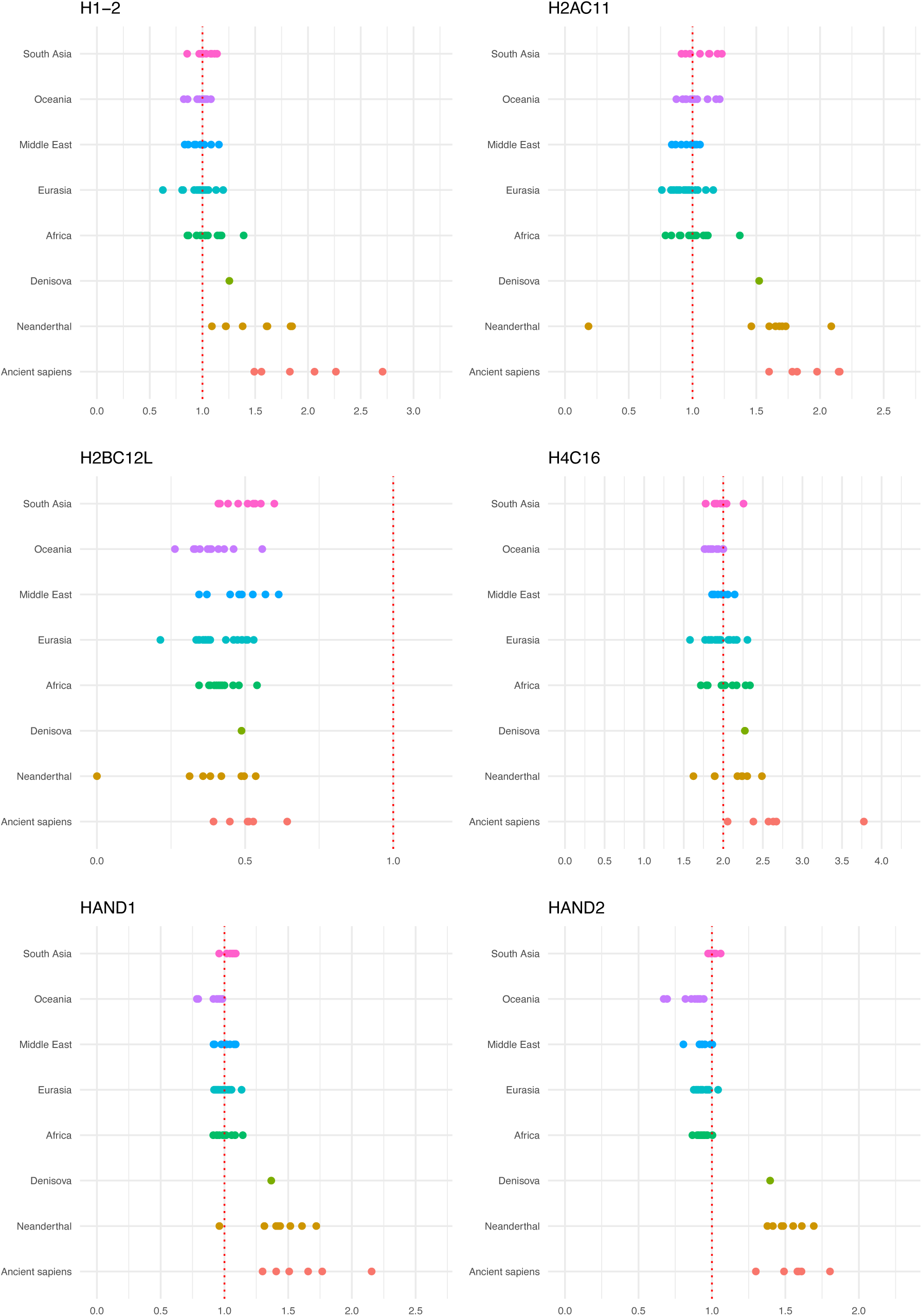
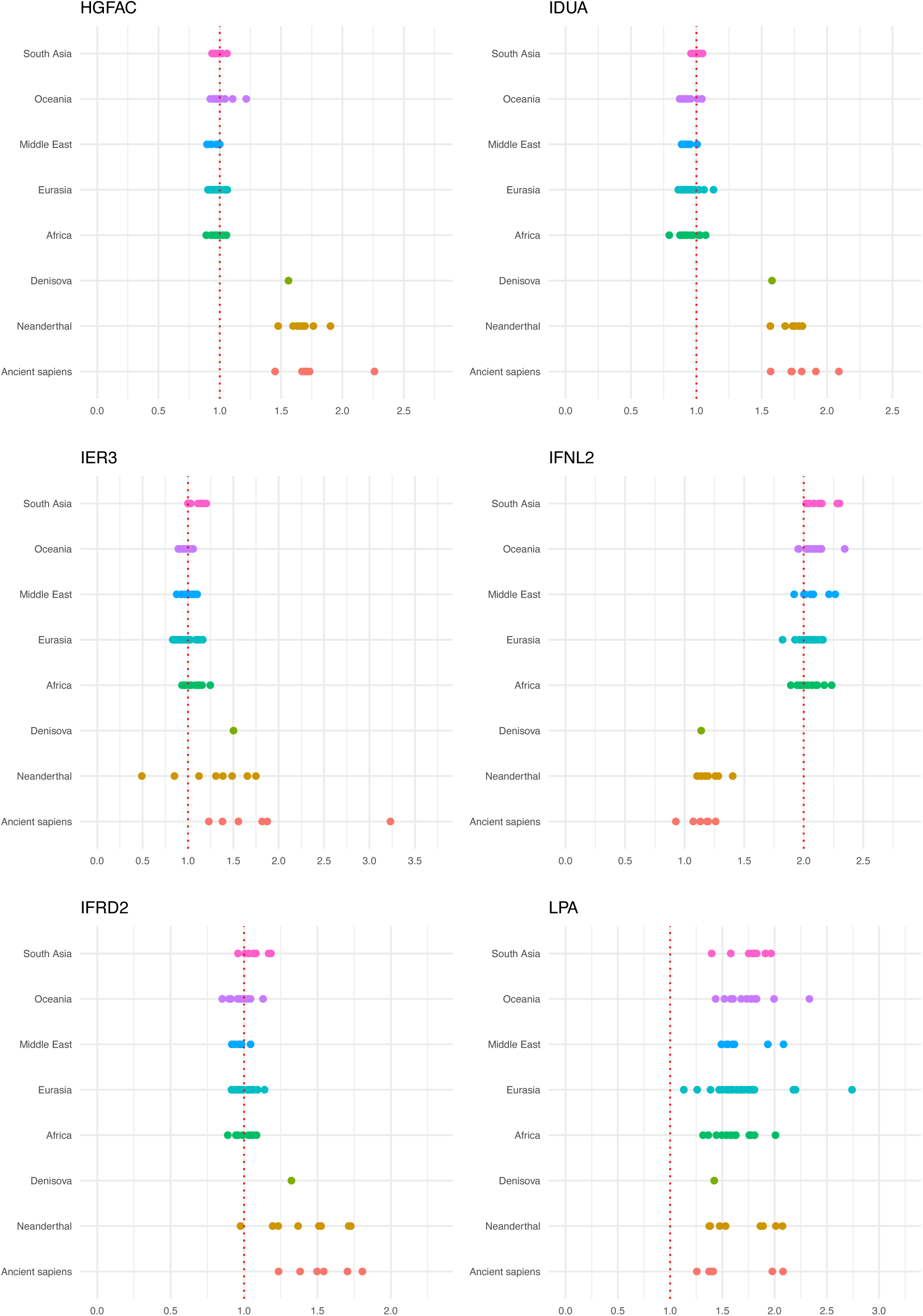
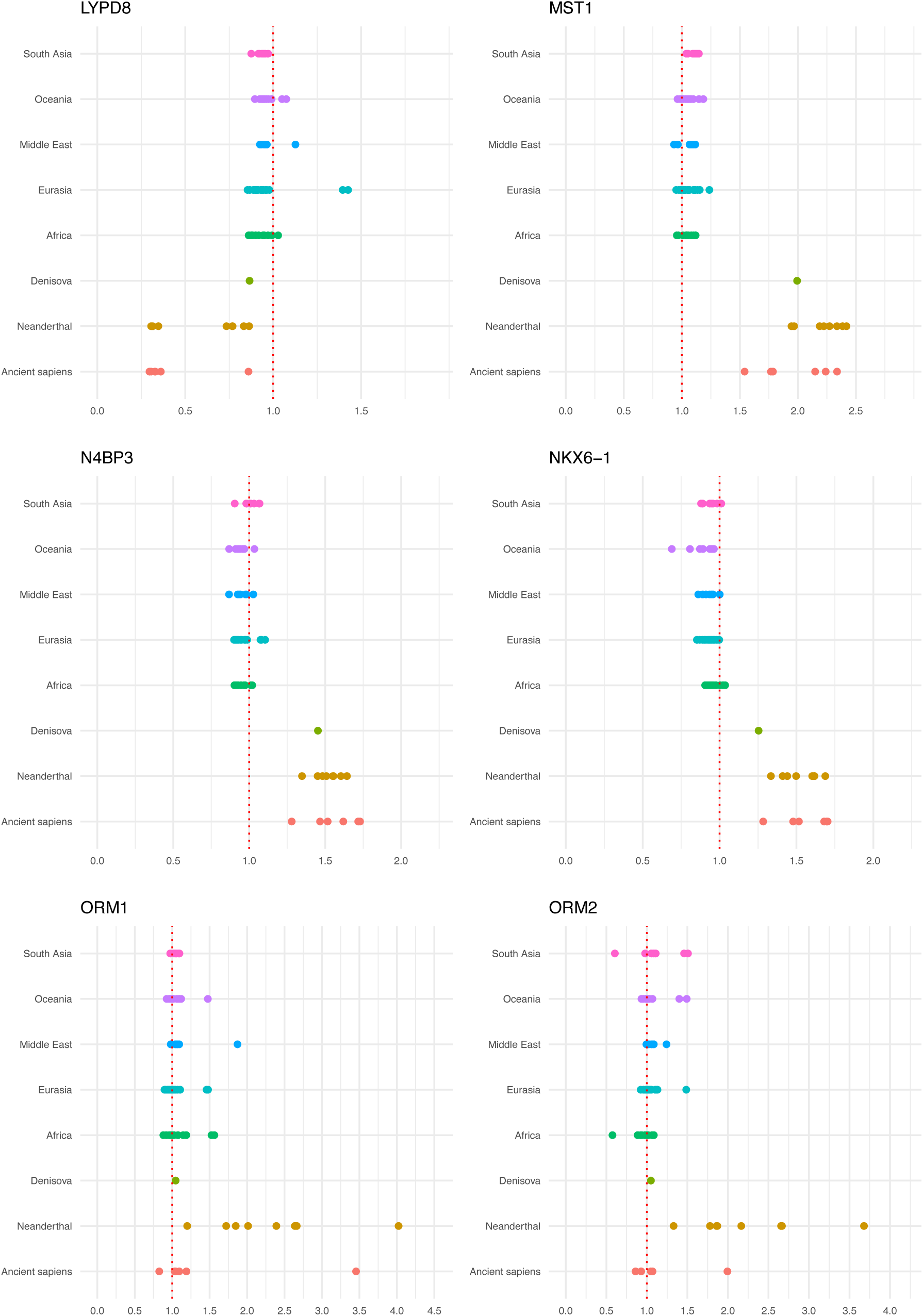
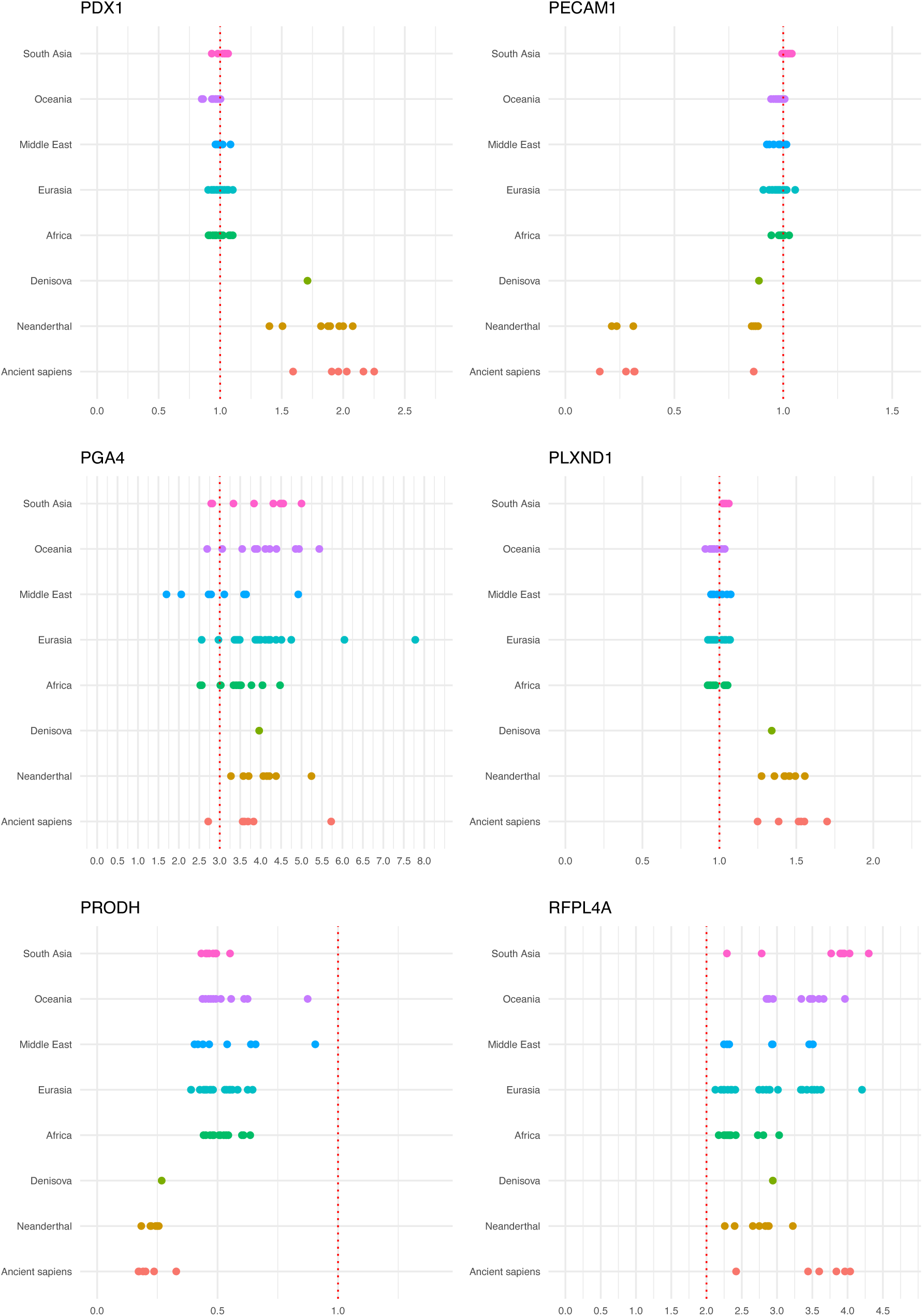
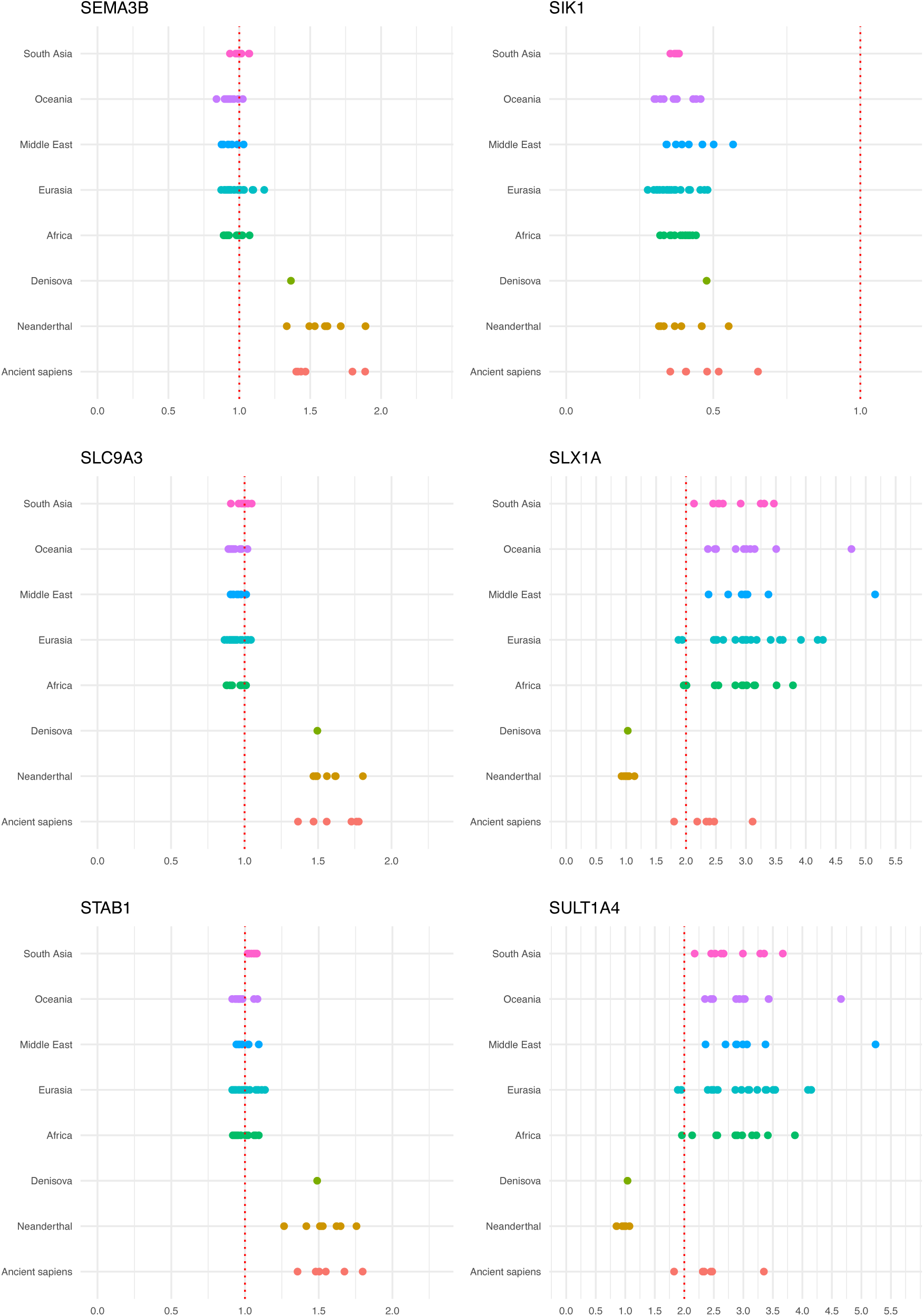
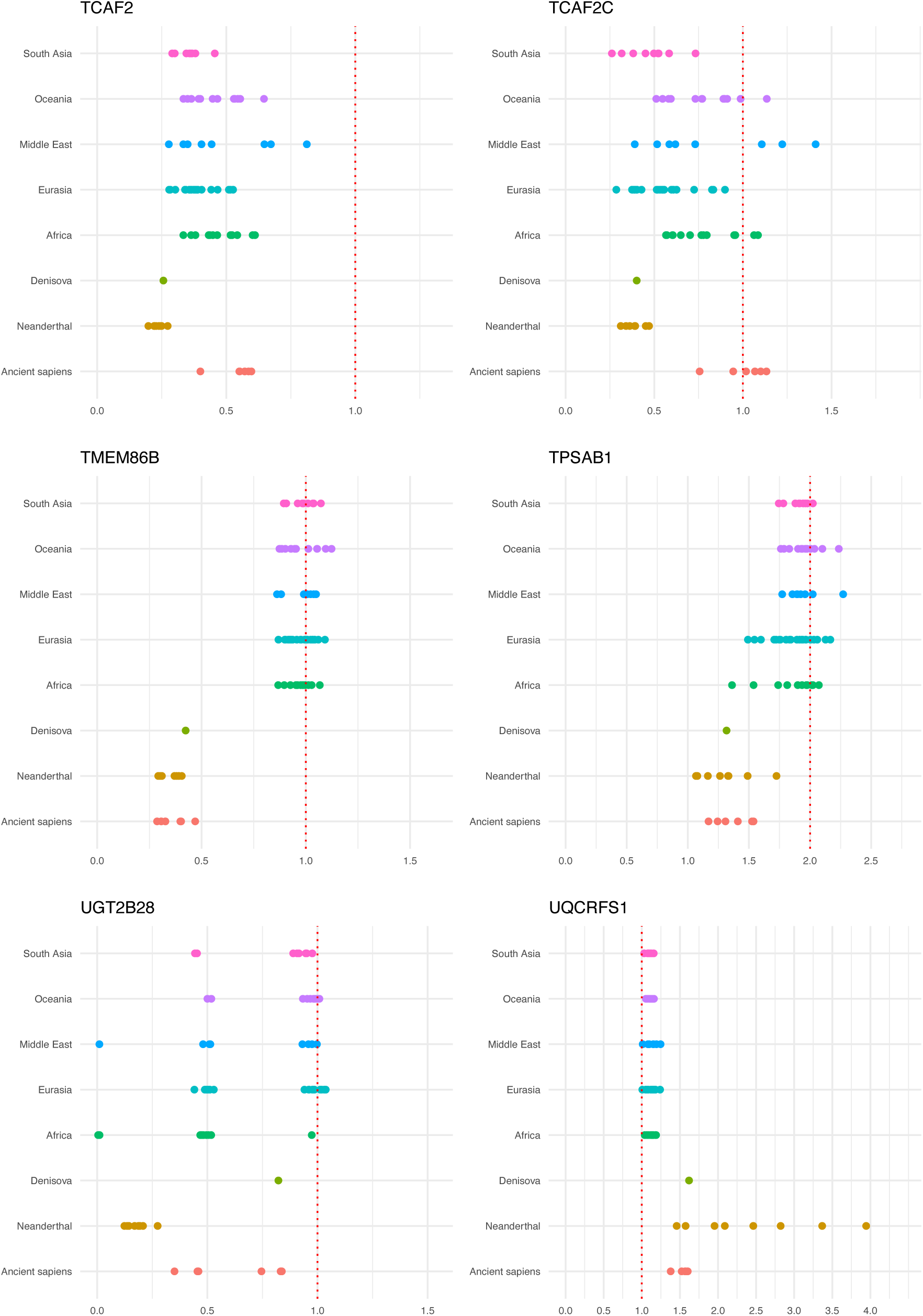
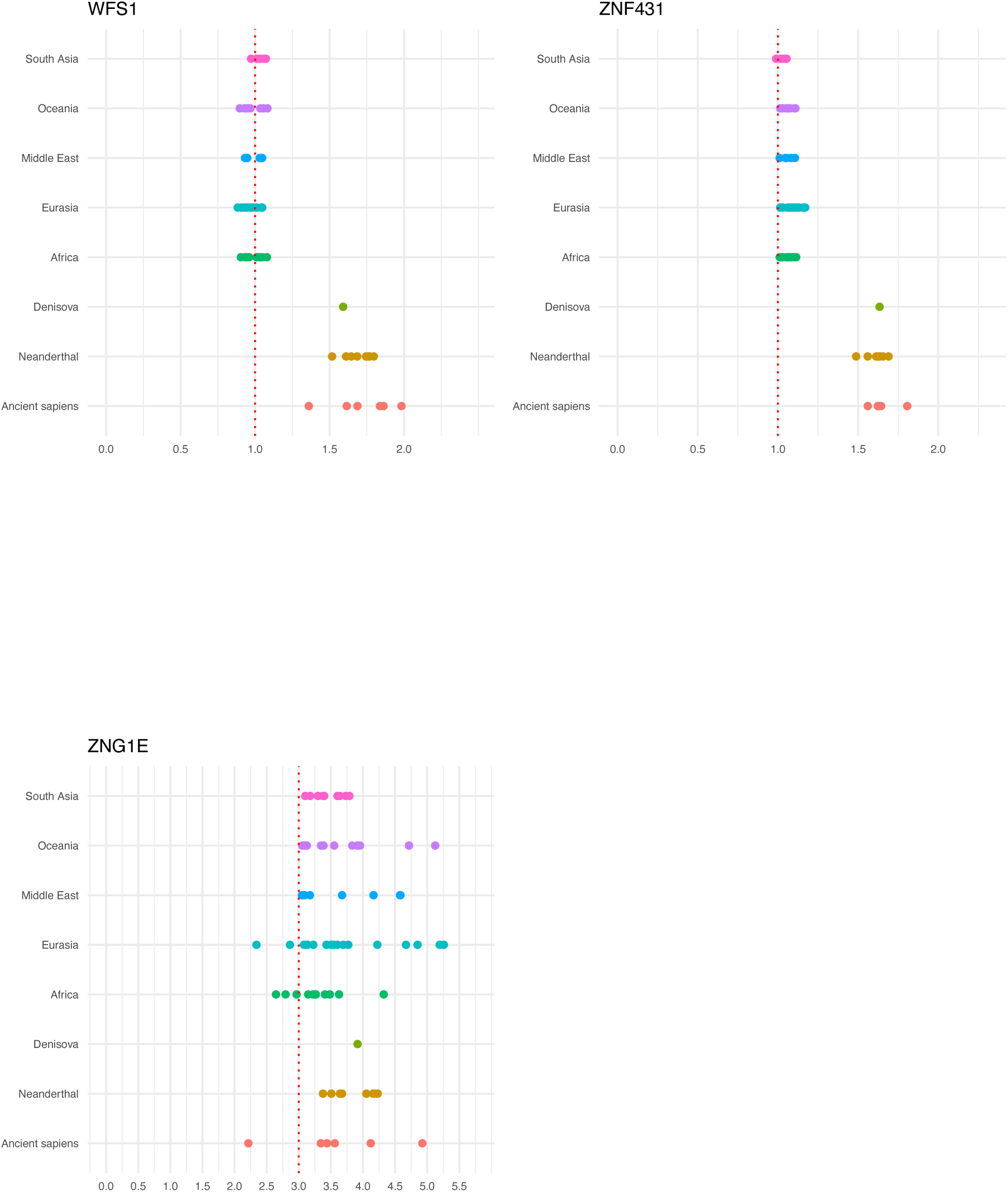
Comparison of CNVs across all modern and ancient individuals in this study, grouped by population. All referenced genes (including diet-related, accessory, and others) are listed in alphabetical order.

**Figure S3.**
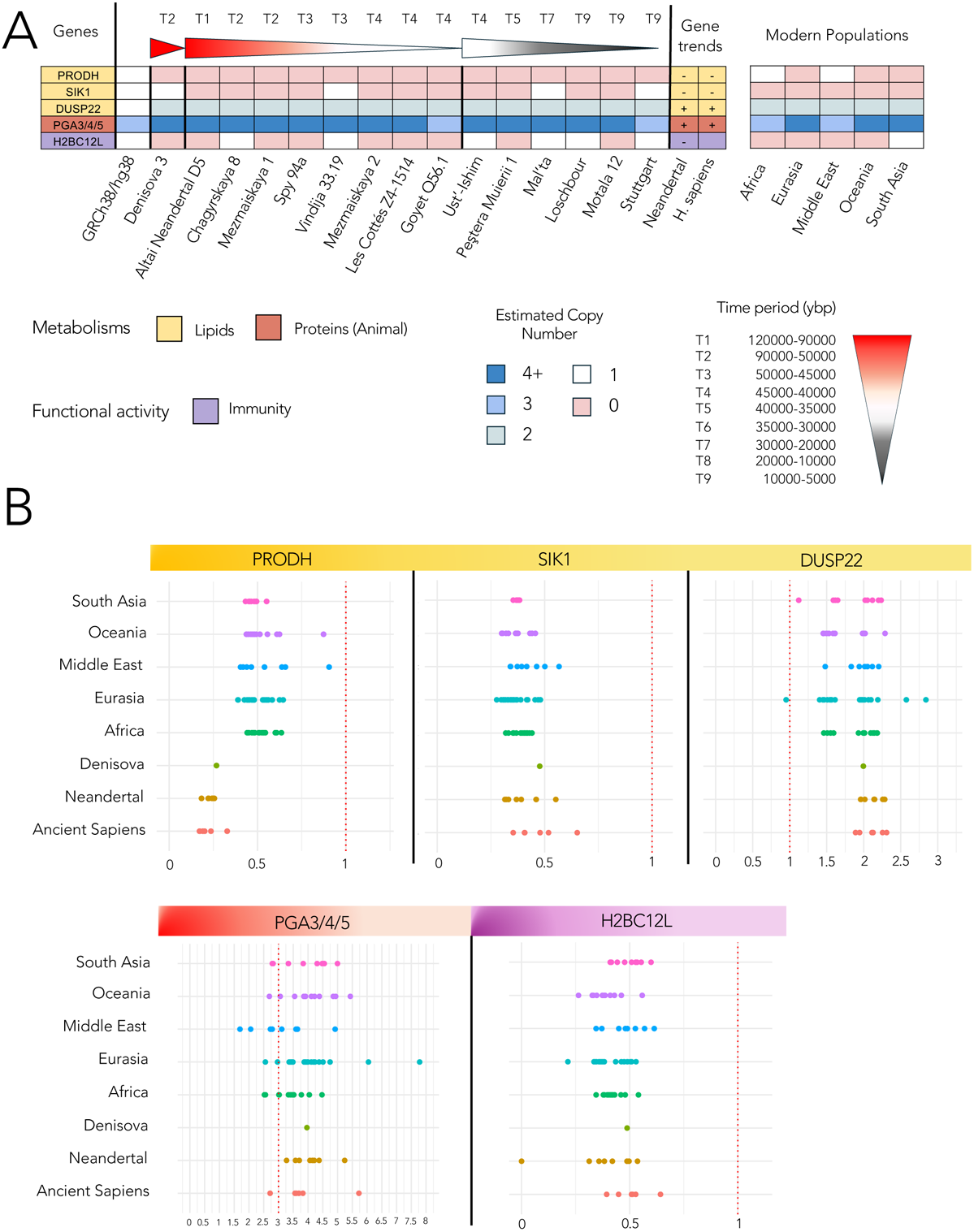
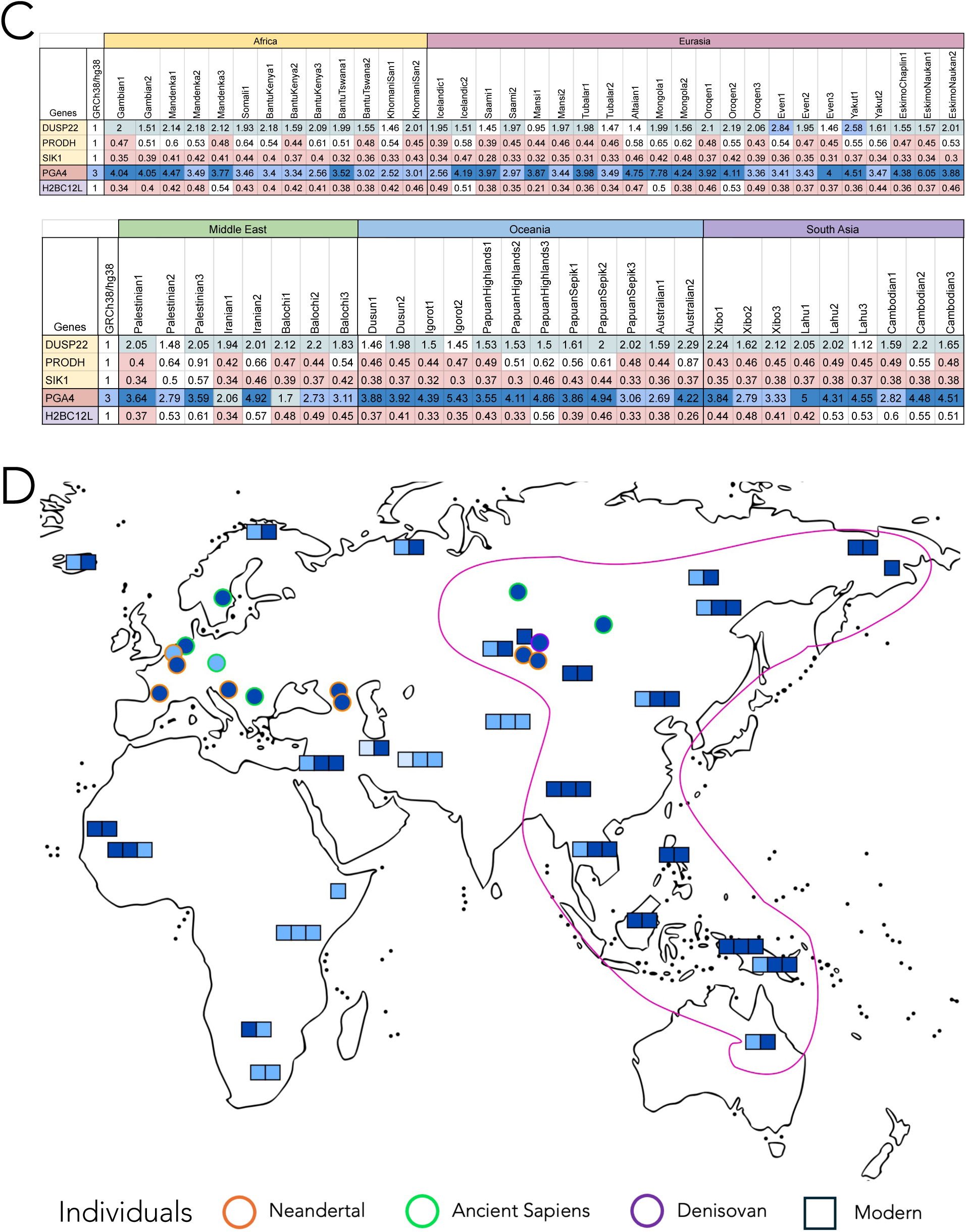
Genes filtered by the analysis of CNV based on CNV distributions on modern populations. Five genes have been filtered out, *PRODH, SIK1, DUSP22, PGA3/4/5* and *H2BC12L*. **A.** Compare with Tables 1 and 2. **B.** Compare with Figure 3. **C.** CNV estimations of the five genes *PRODH, SIK1, DUSP22, PGA3/4/5* and *H2BC12L*. **D.** Visualisation of *PGA3/4/5* haploid CNVs (colors as shown in **A** and **C**) for both modern (square) and ancient (circle) individuals. Different populations are identified by colored borders: *Hn* in orange, *Hs* in green, *Den* in purple, and modern individuals in black. Most modern genomes from the Eastern regions of Eurasia, the South Asia and Oceania (pink shape) exhibit high CNVs, similar to ancient individuals in Eurasia, while modern genomes in the Middle East, and in Africa show lower CNVs.

**Figure S4.**
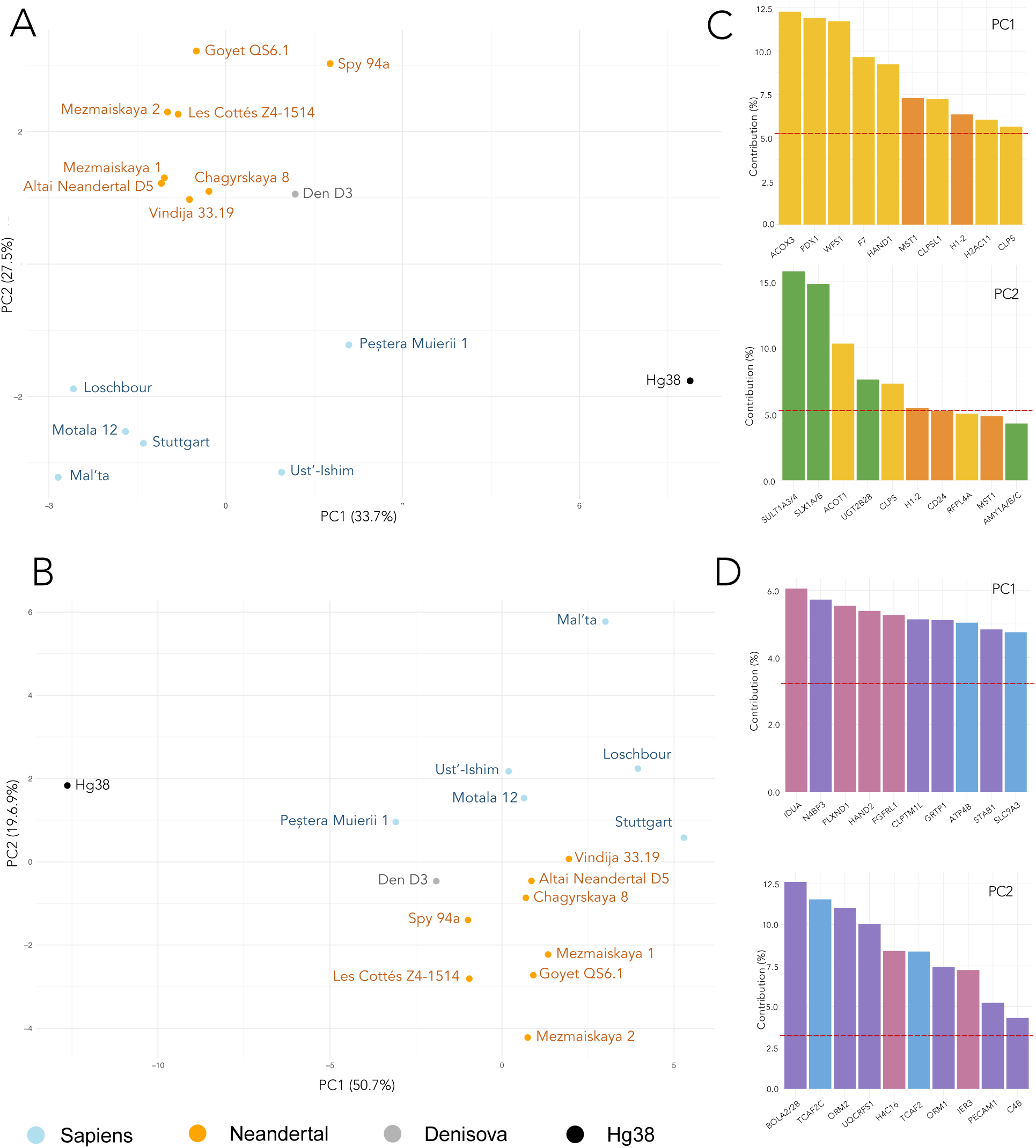
Comparative CNV analysis of *Hn*, *Den* and *Hs* living in the European belt with the reference modern human genome. **A.** Fifteen *Hn*, *Den* and *Hs* individuals and the reference modern genome hg38 are represented in a three-dimensional space obtained by PCA from the 19-dimensional space defined from differential CNV values of the 10 diet-related genes. View of the 2D space where each individual (points) is colored with respect to human populations: *Hn* (orange), *Den* (grey), and *Hs* (light blue). The modern reference genome GRGh38/hg38 is also plotted (blue). PC1 explains the 32.7% of the variance and PC2 the 28.9%. **B.** Same analysis as in A but based on the differential CNV values of the 31 accessory genes. PC1 explains the 49.0% of the variance and PC2 the 20.7%. **C.** Contributions of factors for the definition of each principal component of the 2D space, obtained by PCA for the 19 genes analysis in **A**. Colored bars indicate important contributions from CNV in genes acting in the three overarching metabolic processes (colors as in Figure 2F). **D.** Contributions of factors for the definition of each principal component of the 2D space, obtained by PCA for the 31 genes analysis in **B**. See legend in **C**.

**Figure S5.**
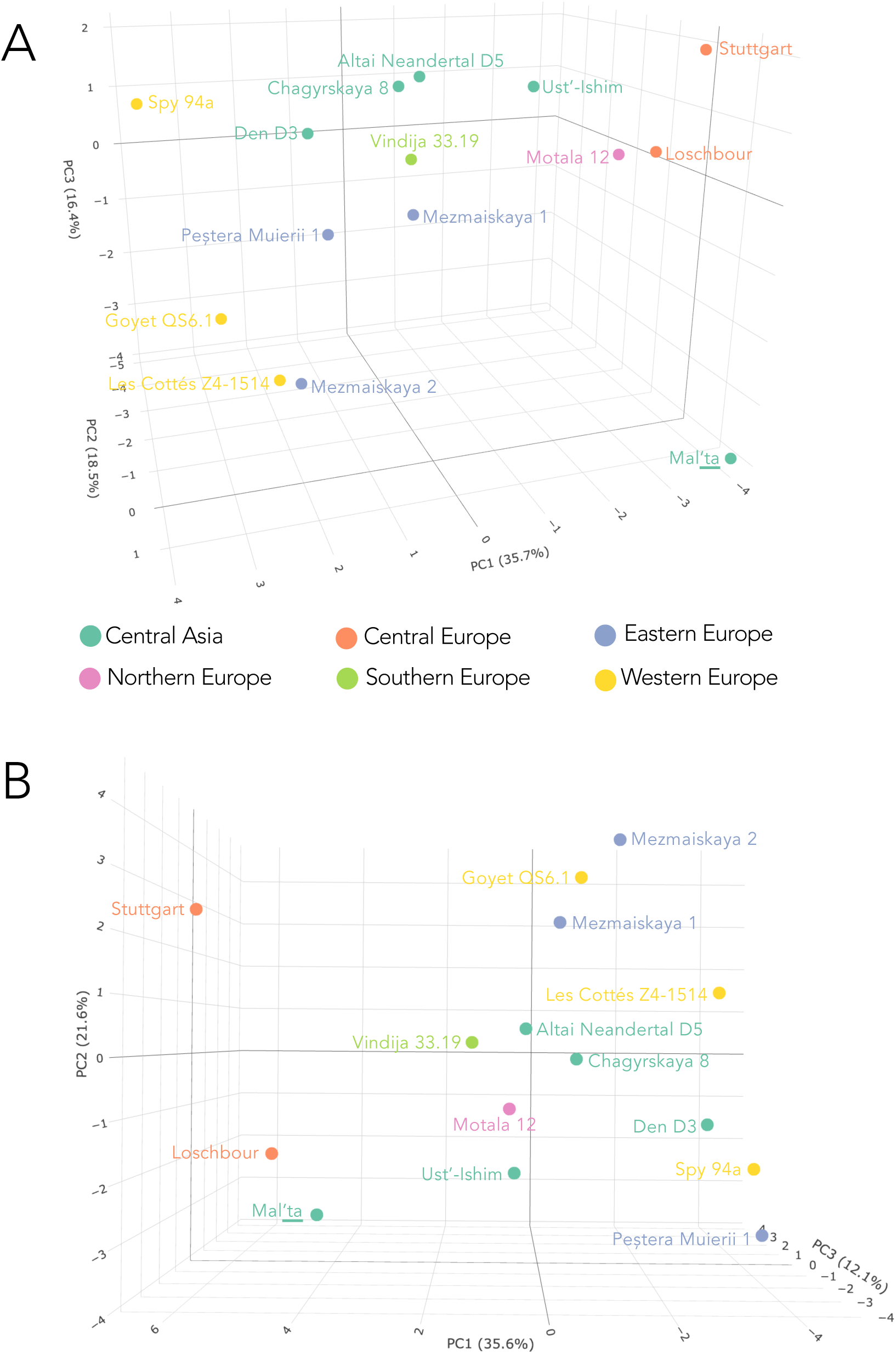
Comparative CNV analysis of *Hn*, *Den* and *Hs* living in the European belt: geographical regions. Complement to Figure 4. **A.** Fifteen *Hn*, *Hs* and *Den* individuals are represented in a three-dimensional space obtained by PCA from the 19-dimensional space defined from diet-related gene CNV (see **Supplementary Text 2**). Individuals are colored with respect to geographical localisation. **B.** As in A but with the 15 individuals plot in the three-dimensional space obtained by PCA from the 31-dimensional space defined from accessory gene CNV.

**Figure S6.**
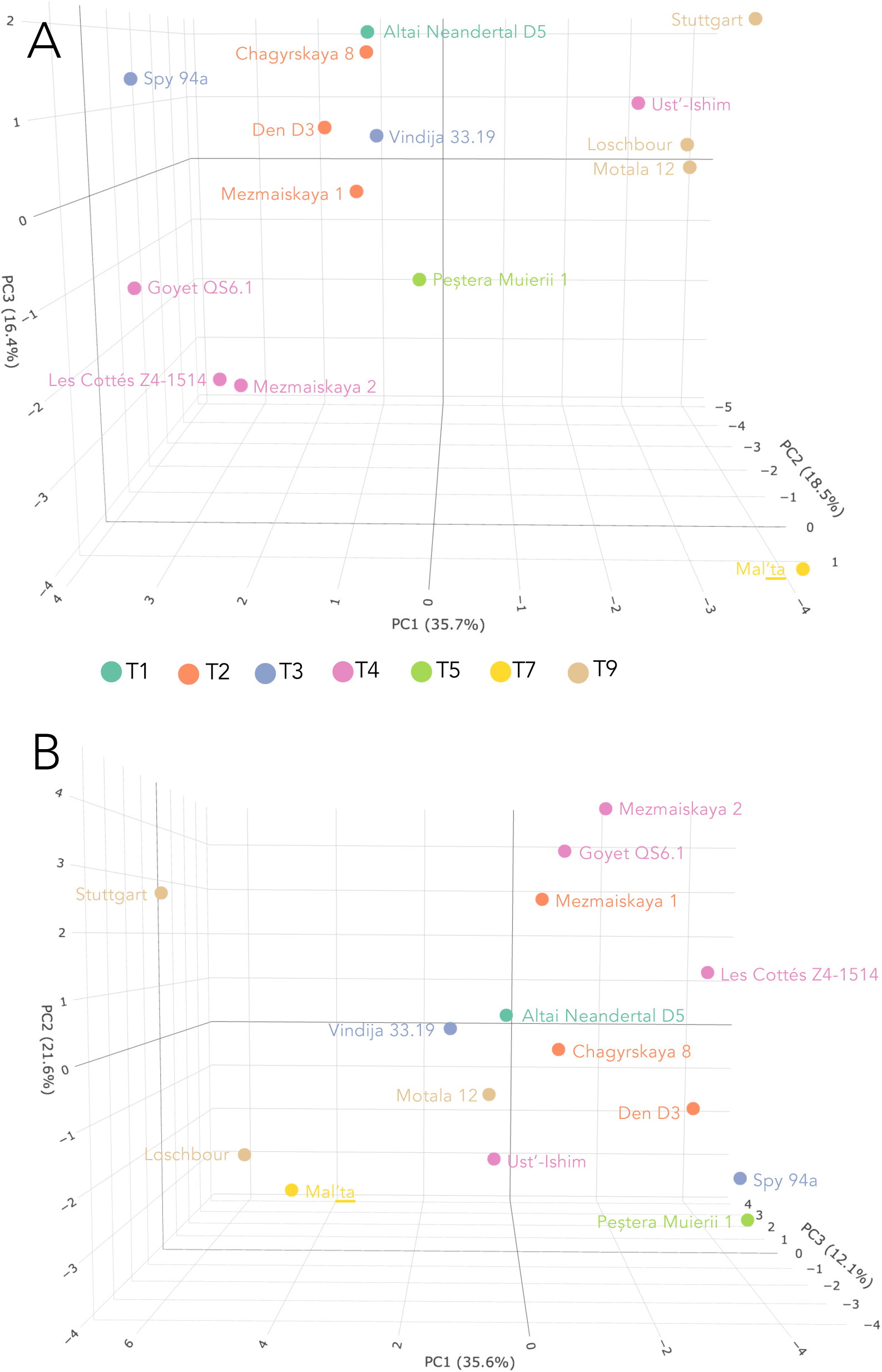
Comparative CNV analysis of *Hn*, *Den* and *Hs* living in the European belt: time period. Complement to Figure 4. **A.** Fifteen *Hn*, *Hs* and *Den* individuals are represented in a three-dimensional space obtained by PCA from the 19-dimensional space defined from diet-related gene CNV (see **Supplementary Text 2**). Individuals are colored with respect to time periods. **B.** As in A but with the 15 individuals plot in the three-dimensional space obtained by PCA from the 31-dimensional space defined from accessory gene CNV.

**Figure S7.**
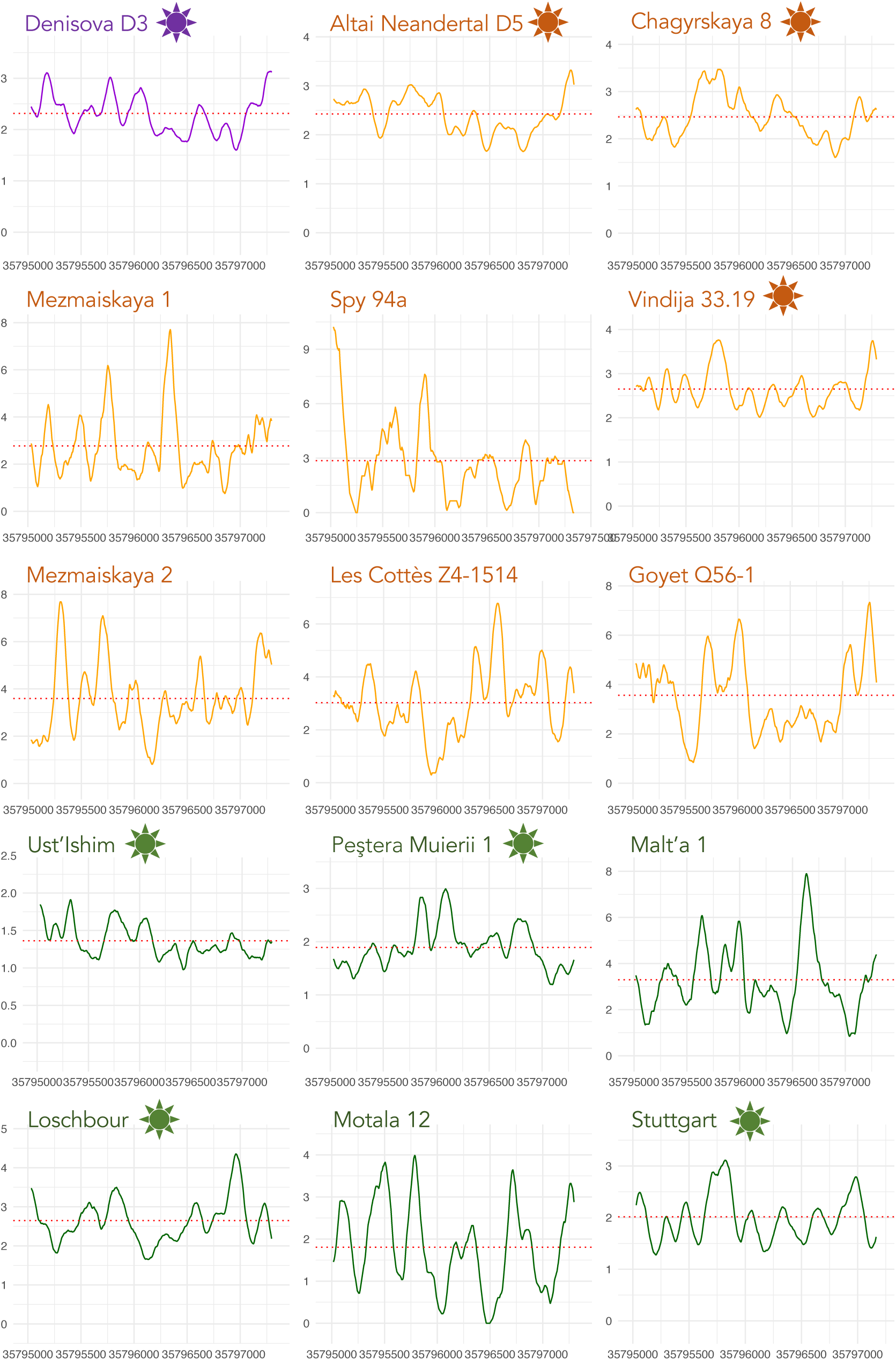
Positional coverage profiles from the read mapping of the CLPS gene. Positional coverage profiles for the CLPS gene across the 15 ancient genomes analyzed in this study, representing Denisovan (purple), Neandertal (orange), and Sapiens (green) populations. The profiles are computed using a sliding window of 100 bp across the gene length. The red dotted line indicates the CNV estimation of CLPS in each genome. The x-axis represents the positions of the CLPS gene on the chromosome, while the y-axis shows the corresponding positional coverage. The star symbol next to the individual’s name indicates a high coverage genome (**Table S1**).

**Figure S8.**
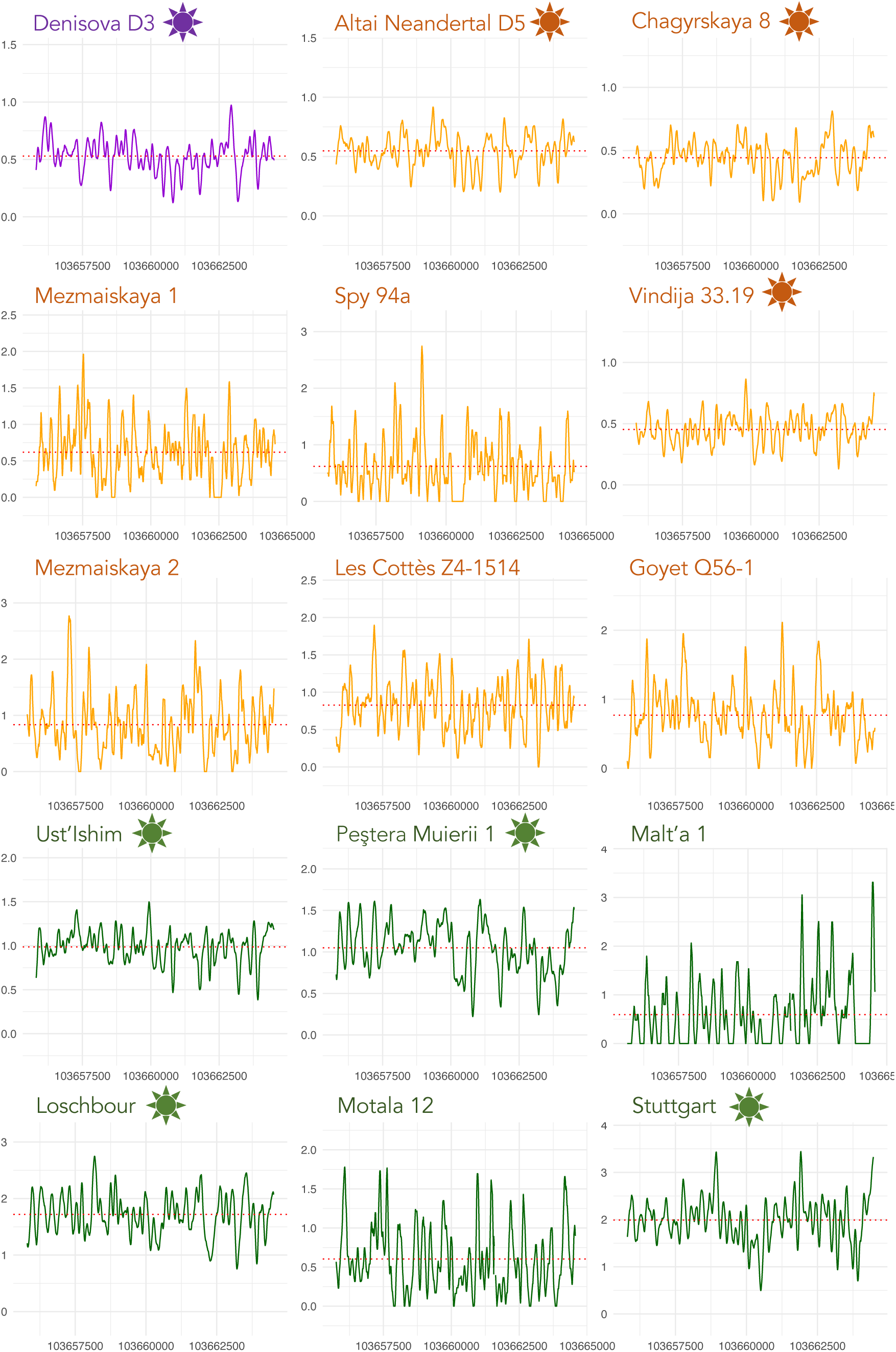
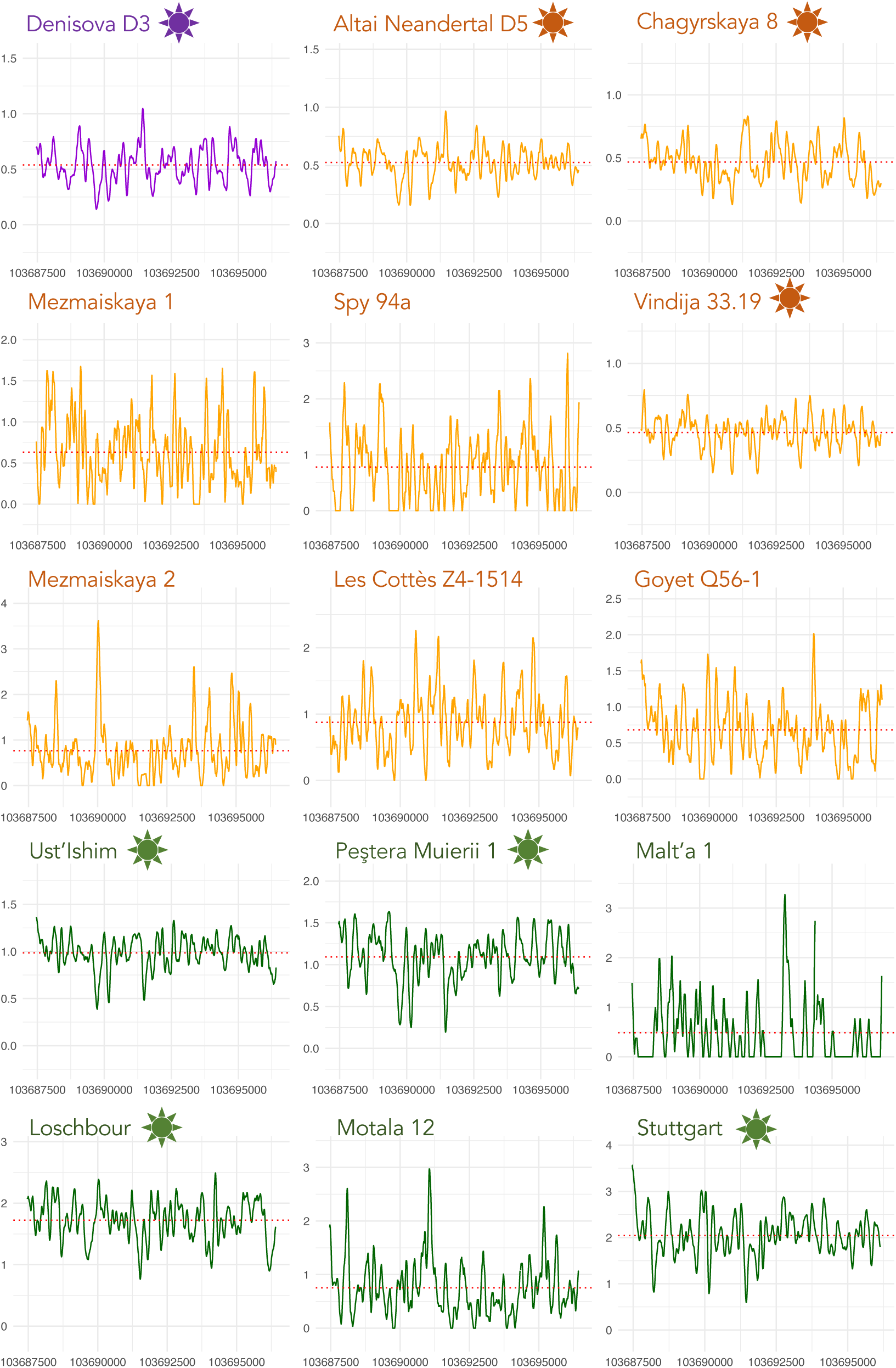
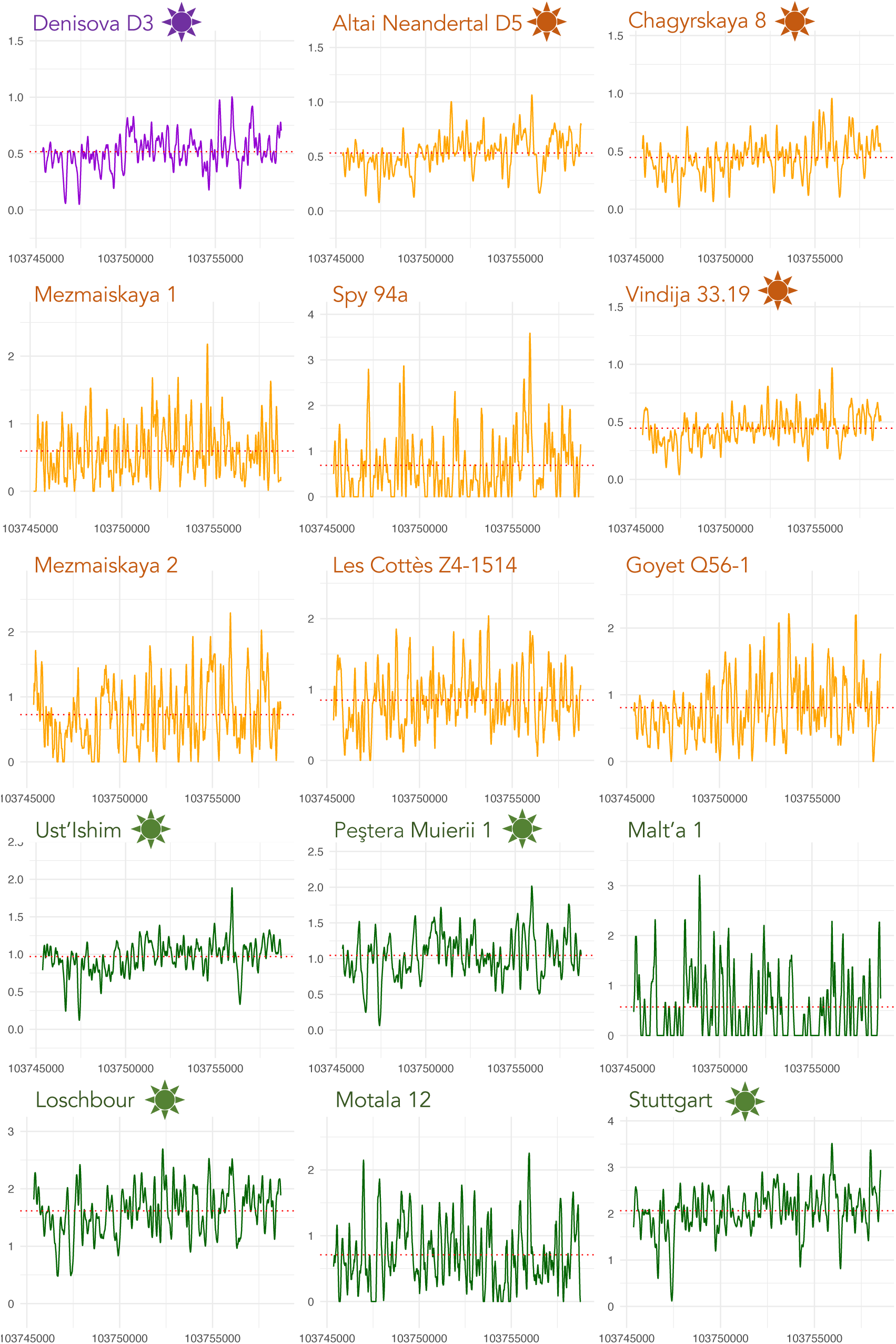
Positional coverage profiles from the read mapping of the the copies of the AMY1A gene. See legend of Figure S7.

## Notes

### Competing Interest Statement

The authors have declared no competing interest.

### Summary of Updates

This revision contain results based on the remapping of all ancient genomes on the human reference genome hg38, where an accurate estimation of gene copy number for the human reference genome is available. By analysing billions of short sequences from 15 archaic human genomes and 64 modern human ones, across the full set of 20,000 human genes, we identify 50 genes whose population-wide discernable CNV trends point to lipid metabolism as being crucial for Neandertal, the efficiency in carbohydrate metabolism for Sapiens' diet, and the importance of lipid metabolism and brown fat metabolism for both Neandertal and early Sapiens compared to modern humans.

